# Localization of adaptive variants in human genomes using averaged one-dependence estimation

**DOI:** 10.1101/229070

**Authors:** Lauren Alpert Sugden, Elizabeth G. Atkinson, Annie P. Fischer, Stephen Rong, Brenna M. Henn, Sohini Ramachandran

**Affiliations:** Department of Ecology and Evolutionary Biology, Brown University, Providence, RI, USA; Center for Computational Molecular Biology, Brown University, Providence, RI, USA; Department of Ecology and Evolution, Stony Brook University, Stony Brook, NY, USA; Division of Applied Mathematics, Brown University, Providence, RI, USA

**Author notes:** correspondence should be addressed to LAS or SR.

## Abstract

Statistical methods for identifying adaptive mutations from population-genetic data face several obstacles: assessing the significance of genomic outliers, integrating correlated measures of selection into one analytic framework, and distinguishing adaptive variants from hitchhiking neutral variants. Here, we introduce SWIF(r), a probabilistic method that detects selective sweeps by learning the distributions of multiple selection statistics under different evolutionary scenarios and calculating the posterior probability of a sweep at each genomic site. SWIF(r) is trained using simulations from a user-specified demographic model and explicitly models the joint distributions of selection statistics, thereby increasing its power to both identify regions undergoing sweeps and localize adaptive mutations. Using array and exome data from 45 ‡Khomani San hunter-gatherers of southern Africa, we identify an enrichment of adaptive signals in genes associated with metabolism and obesity. SWIF(r) provides a transparent probabilistic framework for localizing beneficial mutations that is extensible to a variety of evolutionary scenarios.

## Introduction

Adaptive mutations that spread rapidly through a population, via processes known as selective sweeps, leave distinctive signatures on genomes. These genomic signatures fall into three categories: differentiation among populations, long shared haplotype blocks, and changes in the site frequency spectrum (SFS). Statistics that are commonly used to detect genomic signatures of selective sweeps include *F_ST_*^1^ for measuring population differentiation, iHS^2^ for identifying shared haplotypes, and Tajima’s *D*^3^ for detecting deviations from the neutral SFS. Some approaches, like SweepFinder^4^ and SweeD^5^ integrate information across sites by modeling changes to the SFS. Often, statistical scans for adaptive mutations proceed by choosing a particular genomic signature and a corresponding statistic, obtaining the statistic’s empirical distribution across loci in a genome-wide dataset, and focusing on loci that fall past an arbitrary, but conservative threshold^2,6–11^.

Recently, there has been increased focus on developing composite methods for identifying selective sweeps, which combine multiple statistics into a single framework^12-19^; we refer to the statistics that are aggregated in composite methods such as these as “component statistics”. Most composite methods draw upon machine learning approaches like support vector machinesi^2,15^, deep learning^19^, boosting^14,17^, or random forest classification^18^ in order to identify genomic windows containing selective sweeps. These windows vary in size from 20kb to 200kb, often identifying candidate sweep regions containing many genesi^8,19^. One method, the Composite of Multiple Signals or “CMS”^13,16^, uses component statistics that can be computed site-by-site in pursuit of localizing adaptive variants within genomic windows, but the output from this method cannot be interpreted without comparison to a genome-wide distribution. In addition, CMS must rely on imputation or other methods of compensation when component statistics are undefined, a complication that typically does not arise when using window-based component statistics. In a subset of populations from the 1000 Genomes Project, we found that more than half of variant sites had at least one undefined component statistic (Supplementary Table 1); iHS was frequently undefined because it requires a minor allele frequency of 5% to be computed^2^, and along with XP-EHH^7^, cannot be calculated near the ends of chromosomes or sequenced regions. This poses a particular problem when scanning for complete sweeps, defined here as sweeps in which the beneficial allele has fixed in the population of interest.

Here we introduce a Bayesian classification framework for detecting and localizing adaptive mutations in population-genomic data called SWIF(r) (**SW**eep **I**nference **F**ramework (controlling for **correlation**)). SWIF(r) has three major features that enable genome-wide characterization of adaptive mutations: first, SWIF(r) computes the per-site probability of selective sweep, which is immediately interpretable and does not require comparison with a genome-wide distribution; second, no imputation or compensation mechanisms are necessary in the case of undefined component statistics; and third, we explicitly learn pairwise joint distributions of selection statistics, which gives substantial gains in power to both identify regions containing selective sweeps and localize adaptive variants. Existing composite methods for selection scans have subsets of these features, but SWIF(r) combines all three in a unified statistical framework. Our approach incorporates the demographic history of populations of interest, while being robust to misspecification of that history, and is also agnostic to the frequency of the adaptive allele, identifying both complete and incomplete selective sweeps in a population of interest. We assess SWIF(r)’s performance in simulations against state-of-the-art univariate and composite methods for identifying genomic targets of selective sweeps, and we confirm that we can localize known adaptive mutations in the human genome using data from the 1000 Genomes Project. We then apply SWIF(r) to identify novel adaptive variants in genomic data from the ‡Khomani San, an understudied hunter-gatherer KhoeSan population in southern Africa, representing the most basal human population divergence. Open-source software for training and running SWIF(r) is freely available at https://github.com/ramachandran-lab/SWIFr.

## Results

We first describe the theoretical framework of SWIF(r) and compare SWIF(r) to existing sweep-detection methods using simulated data. We also validate the ability of our method to localize known adaptive mutations in data from the 1000 Genomes. To enable the application of SWIF(r) to single-nucleotide polymorphism (SNP) array datasets from diverse populations, we implement an algorithm for modeling ascertainment bias in simulated data used for training. To illustrate the power of our approach, we apply SWIF(r) to SNP array data from the ‡Khomani San of southern Africa. We show that genes bearing “SWIF(r) signals” — which we define as genomic loci at which SWIF(r) reports a posterior sweep probability greater than 50% — are associated with metabolism and obesity, and we use exome data to show that these genes contain multiple candidate adaptive mutations. Note that we train SWIF(r) on simulations of hard sweeps (Online Methods); our focus here is not on the relative roles of various modes of selection in shaping observed human genomic variation (for recent treatments on this question see^18,20–23^), although we note that SWIF(r) is extensible to multi-class classification, and could be used in future applications to explore multiple modes of selection. In this study, our focus is on localizing genomic sites of adaptive mutations that have spread through populations of interest via hard sweeps.

### Implementation of SWIF(r)

SWIF(r) draws on Bayesian inference and machine learning to localize the genomic site of a selective sweep based on probabilities that incorporate dependencies among component statistics. Unlike genomic outlier approaches, the output of SWIF(r) can be interpreted directly for each genomic site: given a set of n component statistics for a site, SWIF(r) calculates the probability that the site is neutrally evolving or, alternatively, is the site of a selective sweep. We will refer to these two classes as “neutral” and “adaptive” respectively, and these posterior probabilities can be computed as follows:

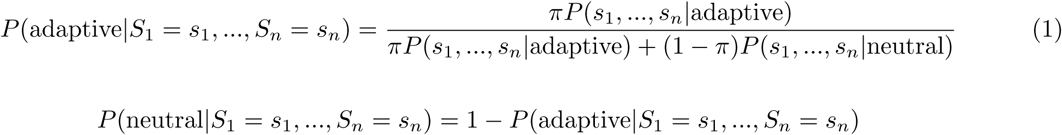

where *s*_1_, …., *s_n_* represent observed values for *n* component statistics such as iHS and *F_ST_*, and *π* is the prior probability of a sweep, which may be altered to reflect different genomic contexts. If a component statistic is undefined at a site, it is simply left out of Equation 1, and does not need to be imputed. The data for learning the likelihood terms, *P*(*s*_1_, …, *s_n_*|adaptive) and *P*(*s*_1_, …, *s_n_*|neutral), come from calculating component statistics on simulated haplotypes from a demographic model with and without simulated selective sweeps comprising a range of selection coefficients and present-day allele frequencies (Online Methods). We note that this general framework is similar to that used by Grossman *et al.*^13^ for CMS, which assumes that the component statistics are independent, and computes the product of posterior probabilities *P*(adaptive|*s_i_*) for each statistic *s_i_*. SWIF(r) strikes a balance between computational tractability and model accuracy by learning joint distributions of pairs of component statistics, thereby relaxing this strict independence assumption.

We base SWIF(r) on a machine learning classification framework called an Averaged One-Dependence Estimator (AODE)^24^, which is built from multiple One-Dependence Estimators (ODEs), each of which conditions on a different component statistic in order to compute a posterior sweep probability at a given site. An ODE conditioning on *S_j_* assumes that all other component statistics are conditionally independent of one another, given the class (neutral or adaptive) *and* the value of *S_j_*. As shown in Equation 2, this assumption effectively reduces the dimensionality of the likelihood terms *P*(*s*_1_,…, *s_n_*|class) in Equation 1. The AODE then reduces variance by averaging all possible ODEs to produce a posterior probability that incorporates all pairwise joint probability distributions (Equation 3; Online Methods).

#### Assumption made by ODE_*j*_

(One-Dependence Estimator conditioning on *S_j_*):

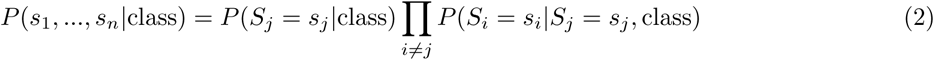

**SWIF(r):**

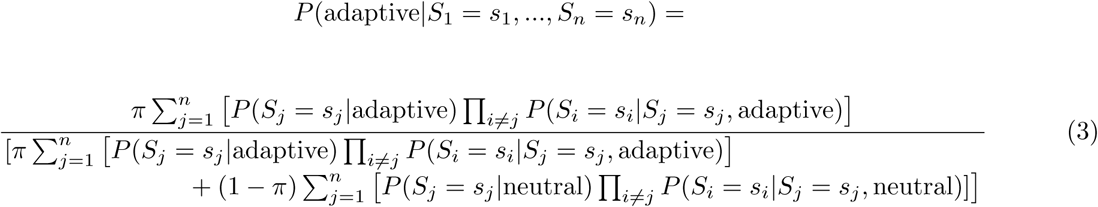

#### Calibration of posterior probabilities calculated by SWIF(r)

A desirable property of probabilities, like those calculated by SWIF(r), is that they be well calibrated: in this context, for the variant positions where the posterior probability reported by SWIF(r) is around 60%, approximately 60% of those sites should contain an adaptive mutation, and approximately 40% should be neutral. We implemented a smoothed isotonic regression scheme to calibrate the probabilities calculated by SWIF(r) (Online Methods). Briefly, when applying SWIF(r) to a given dataset, we calculate the empirical frequencies of neutral and sweep variants that are assigned posterior probabilities between 0 and 1 in simulation, and use isotonic regression^25^ to map the posterior probabilities to their corresponding empirical sweep frequencies (Supplementary Figure 1, Supplementary Figure 2). We then impose a smoothing function that prevents multiple posterior probabilities from being mapped to the same calibrated value (Supplementary Figure 3, Supplementary Figure 1E, Supplementary Figure 2E). This calibration procedure relies on the relative makeup of the training set; a classifier that is calibrated for a training set made up of neutral and sweep variants in equal parts would not be well-calibrated for a training set in which sweep variants only make up 1% of the whole. For each application of SWIF(r) in this study, we calibrated SWIF(r) for a specific training set makeup (Online Methods; see also Supplementary Figure 1 and Supplementary Figure 2).

The calibrated probabilities reported by SWIF(r) can be interpreted directly as the probability that a site contains an adaptive mutation, or fed into a straightforward classification scheme by way of a probability threshold; in this study, we classify sites with a posterior probability above 50% as adaptive SWIF(r) signals. The classifier may be tuned by altering either this threshold or the prior sweep probability *π* (Supplementary Figure 4.

### Performance of SWIF(r) using simulated data

We implemented SWIF(r) using the following component statistics, which can each be calculated site-by-site in a genomic dataset: *F_ST_*^1^, XP-EHH^7^ (altered as in Wagh *et al.*^26^; Supplementary Note), iHS^2^, and difference in derived allele frequency ΔDAF). Training simulations used the demographic model of Europeans, West Africans, and East Asians inferred by Schaffner *et al.*^27^, and simulated selective sweeps within each of those populations (Online Methods). We compared SWIF(r)’s performance against each component statistic, SweepFinder^4^, composite method CMS^13^ (altered by excluding ΔiHH because of non-normality; Supplementary Note, Supplementary Figure 5), and window-based sweep-detection methods evoNet^19^ (using the same component statistics as SWIF(r)), and evolBoosting^14^. We also evaluated the robustness of SWIF(r) to both demographic model misspecification and background selection.

In Figure 1A and Supplementary Figure 6, we evaluate the ability of SWIF(r) to localize the site of an adaptive mutation against that of its component statistic, the composite method CMS, and SweepFinder. The performance of each component statistic varies with different sweep parameters: for example, iHS is most powerful for identifying adaptive mutations that have not yet risen to high frequency within the population of interest, while XP-EHH and ADAF are more effective for those that have (Supplementary Figure 7). This underscores the advantage of composite methods for detecting selective sweeps when the parameters of the sweep are unknown^12–19^. Aggregating over many different sweep parameters, SWIF(r) outperforms each component statistic, as well as CMS and SweepFinder, improving the tradeoff between the false positive rate (fraction of neutral variants incorrectly classified as adaptive) and true positive rate (fraction of adaptive mutations that are correctly classified as such) (Figure 1, Supplementary Figure 6). SWIF(r) also outperforms CMS in distinguishing adaptive mutations from linked neutral variation (Supplementary Figure 8). The performance of SWIF(r) is particularly striking for incomplete sweeps: for example, in Figure 1B, SWIF(r) achieves up to a 50% reduction in the false positive rate relative to CMS for adaptive mutations that have only swept through 20% of the population at the time of sampling (see also Supplementary Figure 6A-C, noting that SweepFinder was designed to identify complete sweeps in a population of interest). For the same incomplete sweep simulations summarized in Figure 1B, Figure 1C shows the performance of each of the individual ODEs (Equation 2); in this particular evolutionary scenario, conditioning on *F_ST_* or ΔDAF results in the best performance. However, the best-performing ODE changes based on the parameters of the selective sweep (Figure 1D). By averaging across all ODEs, SWIF(r) is robust to variable performance of ODEs in the absence of prior knowledge of the true sweep parameters (Figure 1A-C).

**Figure 1.**
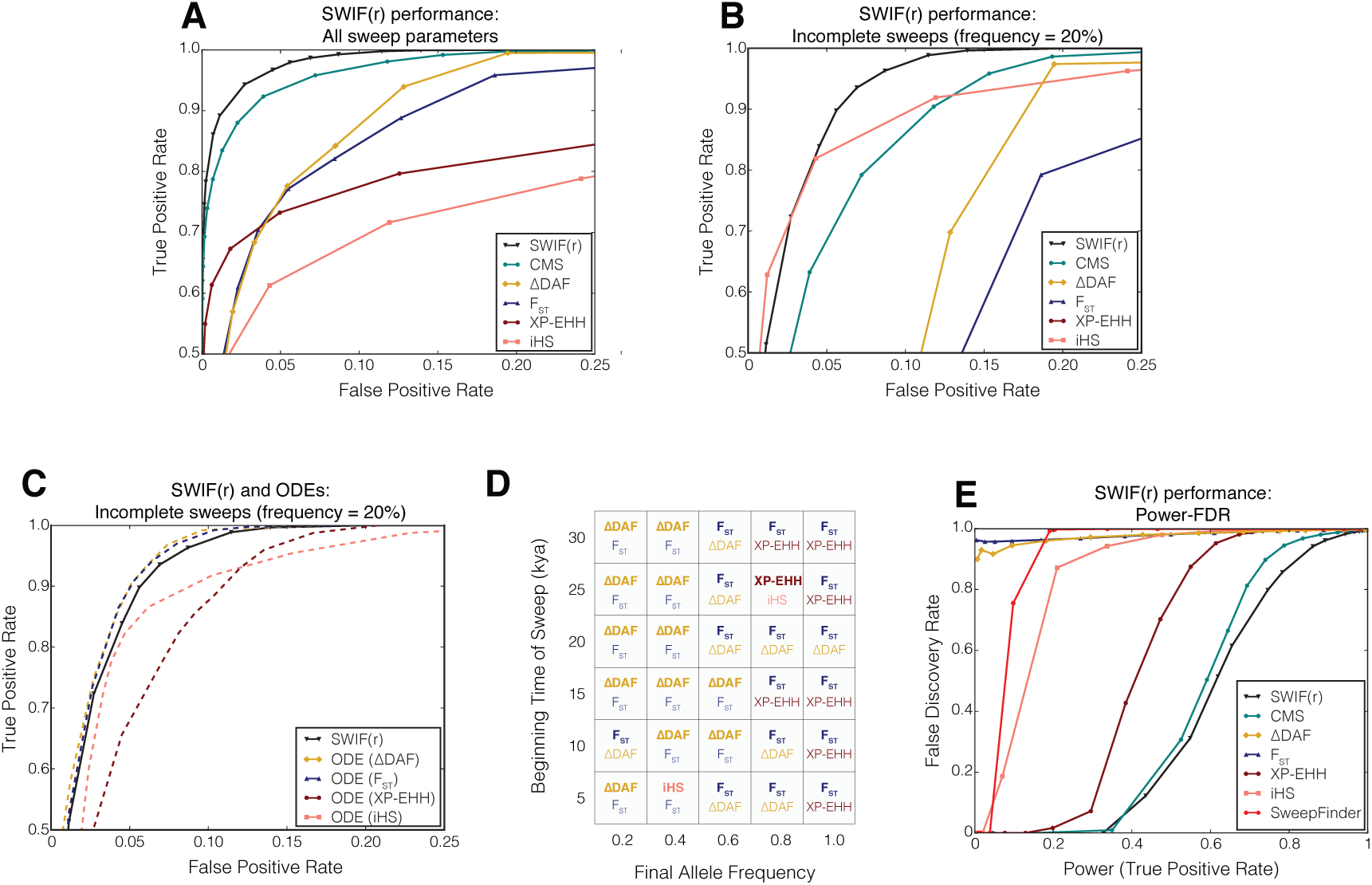
SWIF(r) outcompetes existing site-based sweep-detection methods for a range of sweep model parameters. For comparison against window-based sweep-detection methods, see Supplementary Figure 9 and Supplementary Figure 10. **A)** Receiver operating characteristic (ROC) curves comparing SWIF(r) with CMS^13^, *F_ST_*, iHS, XP-EHH, and ΔDAF across all simulated neutral and sweep scenarios (Online Methods). False positive rate is defined as the fraction of simulated neutral sites that are incorrectly classified as adaptive by a given method, and the true positive rate is defined as the fraction of simulated sites of adaptive mutations that are correctly classified as such. In the context of SWIF(r), sites with posterior sweep probabilities greater than 50% are classified as adaptive. SWIF(r) constitutes an improvement in this tradeoff between true and false positives. For this panel and panel B, the ranges on both axes are such that the ROC curve for SweepFinder^4^ is not visible (Supplementary Figure 6.) **B)** ROC curves for incomplete sweeps in which the beneficial allele has a population frequency of 20%. For these simulations, SWIF(r) reduces the false positive rate by up to 50% relative to CMS. **C)** ROC curves for SWIF(r) and the five component ODEs for incomplete sweeps in which the beneficial allele has a frequency of 20%. SInce the AODE is an average of the ODEs, there will always be individual ODEs that match or outperform SWIF(r); in this case the ODEs conditioned on *F_ST_* and ΔDAF both achieve this. **D)** The two highest-performing ODEs for different sweep parameters. Performance is defined as area under the ROC curve. The statistic that leads to the highest-performing ODE is listed on top in bold, followed by the second best. Text colors correspond to ROC curves in panels A-C. While ODEs conditioned on ΔDAF tend to perform extremely well for sweeps that are at lower frequency, ODEs conditioned on *F_ST_* and XP-EHH tend to perform better for sweeps that are near-complete or complete in the population of interest. By averaging the ODEs, SWIF(r) is robust to uncertainty about the true parameters of the sweep. **E)** Power-FDR curves for SWIF(r), CMS, *F_ST_*, iHS, XP-EHH, ΔDAF, and SweepFinder. Power is equivalent to true positive rate as defined in panels A-C, and false discovery rate is defined as the fraction of sites classified as adaptive that are actually neutral. These curves assume a training set composed of 99.95% neutral variants and 0.05% adaptive variants. See Supplementary Figure 11 for curves based on other training set compositions.

While few composite methods for sweep detection operate site-by-site, there are a handful of machine-learning composite approaches that identify genomic windows containing adaptive mutationsi^4,17–19^. In order to compare SWIF(r) against such methods, we had to alter SWIF(r) to calculate window-based sweep probabilities; there are many potential ways to do this that may be differentially powerful, and here we chose simply to use the highest probability assigned to any variant within a given genomic window as the probability for that window. We compared window-based SWIF(r) to two state-of-the-art composite window-based methods: evolBoosting^14^, which combines 120 statistics using boosted logistic regression, and evoNet^19^, which was developed to jointly infer demography and selection using a deep learning framework (Online Methods). SInce both SWIF(r) and evoNet are frameworks that are designed to incorporate any set of statistics, we implemented evoNet to use the same statistics as SWIF(r). When comparing SWIF(r) with evolBoosting, we used 40kb windows following Lin *et al.*^14^, and show that SWIF(r) outperforms evolBoosting across a range of sweep parameter values (Supplementary Figure 9). For comparison with evoNet, we used 100kb windows following Sheehan *et al.*^19^ (Supplementary Figure 10). We find that SWIF(r) performs similarly to or better than evoNet in this implementation, although we note that this analysis likely downplays the strengths of both methods, as each method has been altered from its original design to enable a direct comparison.

While ROC curves are informative for illustrating the performance of different sweep detection methods, it is important to note that the genome has far more neutral variants than adaptive mutations. Therefore, more relevant performance comparisons can be made by illustrating the predicted false discovery rate (FDR) for a given true positive rate using Power-FDR curves. These curves depend on the composition of the training set, since the false discovery rate rises as the proportion of adaptive variants in the training set decreases. In Figure 1E and Supplementary Figure 6F, we plot Power-FDR curves for all methods using the same training set composition we use for calibration of SWIF(r) for application to the ‡Khomani San dataset (99.95% neutral variants and 0.05% adaptive variants). Curves for other training set compositions can be found in Supplementary Figure 11. Power-FDR comparisons between SWIF(r) and evolBoosting and evoNet can be found in Supplementary Figure 9F and Supplementary Figure 10F assuming that 1% of windows contain a sweep. SWIF(r) performs well relative to its component statistics, CMS, and SweepFinder, however, these analyses illustrate the inherent difficulty of site-by-site detection of adaptive mutations. Because there are so many more neutral variants than adaptive variants in the genome, even a small false positive rate can result in a substantial false discovery rate. Window-based methods, including window-based SWIF(r), may appear to have lower false discovery rates (since there are many fewer windows than variants, and thus fewer opportunities for false positives to arise; see Supplementary Figure 9 and Supplementary Figure 10), but this comes at the cost of a longer list of putative SNP targets, since each classified window contains a large number of individual variants.

#### Robustness of SWIF(r) to demographic model misspecification

We first assessed the sensitivity of SWIF(r) to the demographic model used in training simulations using two very different demographic models of West Africans, Europeans, and East Asians, from Schaffner *et al.*^27^ (“Schaffner model”) and Gronau *et al.*^28^ (“Gronau model”). These demographies differ in multiple evolutionary parameter estimates: the Schaffner model includes an ancient population expansion and post-divergence bottlenecks, features not included in the Gronau model. Divergence times differ almost two-fold in some cases between the two models, with the Yoruban/Eurasian split at 47kya in the Gronau model and 88kya in the Schaffner model. Furthermore, the Schaffner model allows migration between East Asian, European, and West African populations, while the Gronau model does not. Effective population sizes also differ dramatically, in some cases over six-fold (see Supplementary Figure 12 for a full comparison).

We trained SWIF(r) using simulations from the Gronau model (including simulation of ascertainment bias; Online Methods), and tested SWIF(r) on simulated haplotypes drawn from the Schaffner model (with and without selective sweeps). As shown in Supplementary Figure 13, even with dramatic demographic differences and thinning to simulate ascertainment bias, SWIF(r) is quite robust to this misspecification.

Both the Schaffner model and the Gronau model, as implemented, do not include very recent population expansion, so we implemented a third demographic model from Gravel *et al.*^29^ that includes exponential population growth within the last 23,000 years. SInce cosi cannot simulate selective sweeps overlapping with demographic changes, we only simulated sweeps beginning 5kya, allowing for 18,000 years of exponential expansion (Online Methods). In Supplementary Figure 13, we show that SWIF(r) is also robust to this recent population expansion.

#### Robustness of SWIF(r) to background selection

We also assessed the sensitivity of SWIF(r) to background selection by generating a set of training simulations containing neutral regions, selective sweeps, and exonic regions, using forward simulator `slim`^30^ (Online Methods). We trained SWIF(r) on both neutral and sweep simulations, and tested the ability of SWIF(r) to distinguish between exonic and sweep sites, relative to its ability to distinguish between neutral and sweep sites. We find that SWIF(r) is fully robust in this scenario, meaning that we would not expect background selection in genic regions to result in false positive sweep signals (Supplementary Figure 14). These results are aligned with those of Enard *et al.*^31^, who have shown that background selection has little to no effect on haplotype-based statistics iHS and XP-EHH (and in fact makes iHS more conservative).

### SWIF(r) correctly localizes canonical adaptive mutations in humans

For application to data from phase 1 of the 1000 Genomes Project, we used training simulations from the Schaffner demographic model^27^, calibrated SWIF(r) for a training set composed of 0.01% sweep variants and 99.99% neutral variants (Supplementary Figure 1), and applied it to SNP array data from West African (YRI), East Asian (CHB and JPT), and European (CEU) populations. SWIF(r) reports high sweep probabilities at multiple SNPs within known and suspected selective sweep loci in each of these populations (Supplementary Table 2, Supplementary Table 3). Figure 2 illustrates the ability of SWIF(r) to localize sites of adaptive mutations within genomic regions containing canonical sweeps. Adaptive SNPs have been determined via functional experiments in *SLC24A5*^32^, *DARC*^33^, and *HERC2*^34^; we find that modeling the dependency structure among component statistics within SWIF(r) enables statistical localization of these experimentally identified adaptive mutations (Figures 2A,C,D). Methods that treat component statistics as independent, as CMS does, cannot localize these experimentally identified adaptive SNPs (Supplementary Figure 15, Supplementary Figure 16). In CHB and JPT, SWIF(r) recovers a strong adaptive signal in the vicinity of *EDAR*, offering new hypotheses for targets of selection in this genomic region. Whole-genome results with gene annotations can be found in Supplementary Figure 17, Supplementary Figure 18, Supplementary Figure 19 and Supplementary Table 2 (see also Supplementary Figure 20). False discovery estimates can be found in Supplementary Table 4. Of the 126 genes across these populations with SWIF(r) signals (i.e. at least one variant within the gene has posterior hard sweep probability greater than 50%), 63% were identified in at least one positive selection scan conducted in humans (Supplementary Table 3).

**Figure 2.**
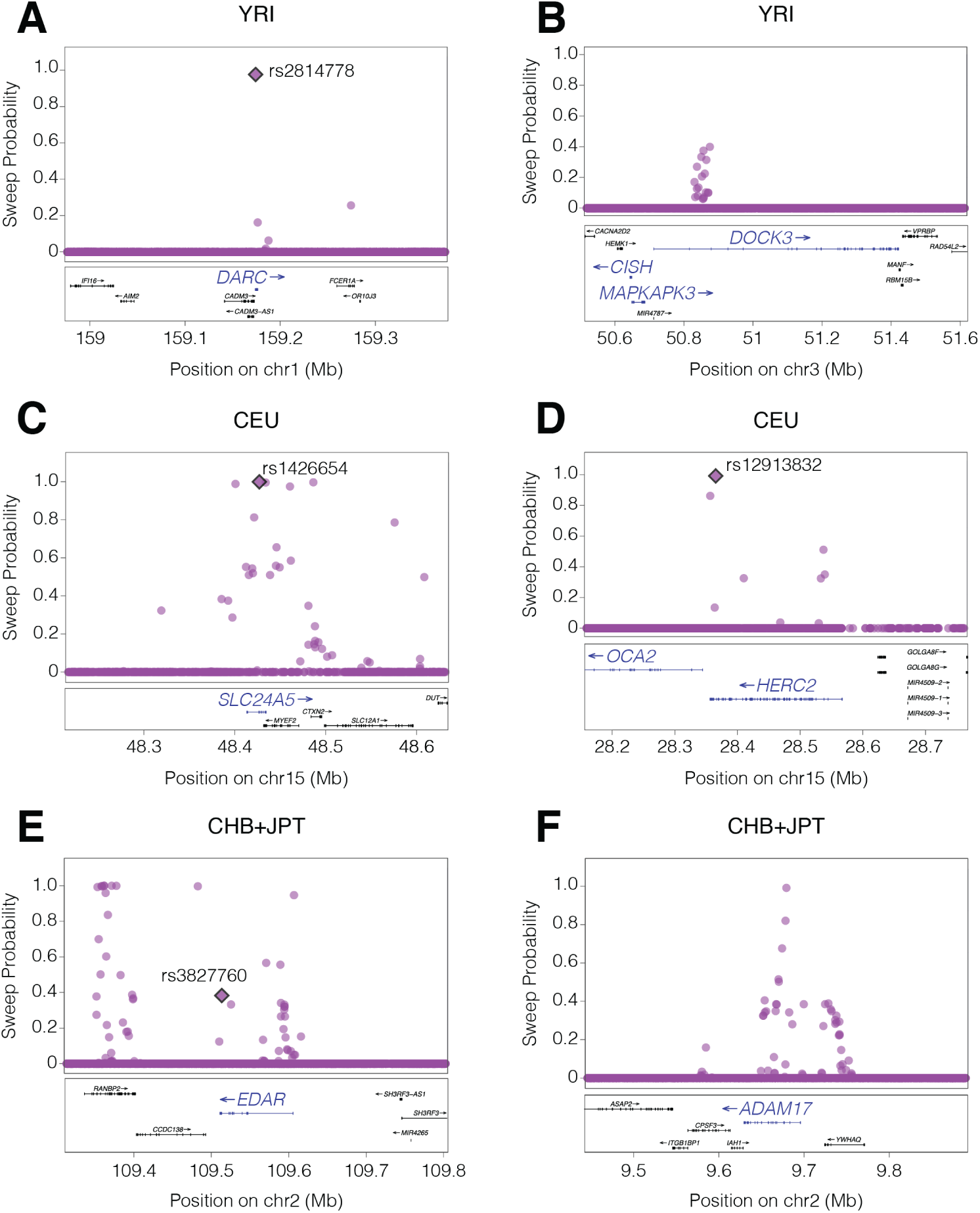
SWIF(r) localizes canonical adaptive mutations and provides additional evidence for suspected sweeps in YRI, CHB+JPT, and CEU^35^. Plotted points indicate the calibrated posterior sweep probability calculated by SWIF(r) at each site, using prior sweep probability **π** = 10^−5^. Plots were made with LocusZoom^36^. Where available, functionally verified adaptive SNPs are depicted as filled diamonds and labeled with rsids. **A-B)** In YRI, two loci where SWIF(r) reports high sweep probabilities are *DARC* and *DOCK3. DARC* encodes the Duffy antigen, located on the surface of red blood cells, and is the receptor for malaria parasites. The derived allele of the causal SNP shown has been determined to be protective against *Plasmodium vivax* malaria infection^33^. *DOCK3*, along with neighboring genes *MAPKAPK3* and *CISH*, are all associated with variation in height, and have previously been shown to harbor signals of selection in Pygmy populations^37^. *CISH* may also play a role in susceptibility to infectious diseases, including malaria^38^. **C-D)** In CEU, we uncover multiple loci in genes involved in pigmentation, including rsi426654 in *SLC24A5*, which is involved in light skin color^32^, and rsi2913832 in the promotor region of *OCA2*, which is functionally linked with eye color and correlates with skin and hair pigmentation^34^. rs1426654 has the highest sweep probability reported by SWIF(r) in *SLC24A5* (0.999173321, after smoothed calibration; see Supplementary Table 2); note each panel depicts genomic windows containing multiple genes. **E-F)** In CHB and JPT, SWIF(r) recovers a strong adaptive signal in the vicinity of *EDAR*; multiple GWA studies have shown rs3827760 to be associated with hair and tooth morphology^39–41^. SWIF(r) also identifies variants with high sweep probability in *ADAM17*, which is involved in pigmentation^42^, and has been identified in other positive selection scans in East Asian individuals^43,44^.

### Adaptive loci in the ‡Khomani San are enriched for metabolism- and obesity-related genes

We applied SWIF(r) to samples from the ‡Khomani San, a formerly hunter-gatherer KhoeSan population of the Kalahari desert in southern Africa, using Illumina SNP array data for 670,987 phased autosomal sites genotyped in 45 individuals^45^ (Supplementary Note). The KhoeSan have likely occupied southern Africa for ∼100,000 years, and maintain the largest long-term *N_e_* of any human population^46,47^, a feature that facilitates adaptive evolution. We trained SWIF(r) on simulations from the Gronau demographic model^28^ (Supplementary Figure 21, Supplementary Table 5, and Online Methods), and implemented an ascertainment modeling scheme to produce a training dataset with population-level site frequency spectra similar to the observed array data. Briefly, for each simulated haplotype, SNPs were subsampled to match the empirical three-dimensional unfolded SFS for YRI, CEU, and CHB+JPT individuals in the 1000 Genomes Project on the chips used to genotype the ‡Khomani San (Online Methods, Supplementary Figure 22, Supplementary Figure 23). We calibrated SWIF(r) for this dataset based on a training set composed of 0.05% sweep variants and 99.95% neutral variants (Supplementary Figure 2). After applying SWIF(r) to SNP data, we then examined whether genomic regions identified by SWIF(r) contain annotated functional mutations identified in high-coverage exome data from the same 45 individuals^48^ (Supplementary Note).

SWIF(r) identifies a number of genomic regions bearing signatures of selective sweeps in the ‡Khomani San, driven by extreme values in multiple component statistics that together produce a posterior sweep probability greater than 50% (Figure 3A,B; see also Supplementary Figure 24, Supplementary Figure 25, and Supplementary Table 6). These signals comprise 108 SNPs, of which 94 are distributed across 80 genes, and the remaining 14 are intergenic, defined as genomic variants that do not land within 50kb of an annotated gene (Supplementary Table 7). We observe an abundance of SWIF(r) signals within the Major Histocompatibility Complex (MHC), a region of immunity genes for which studies have indicated ongoing selection in many populations^49–52^, including an iHS outlier scan in the ‡Khomani San^53^. We show in Supplementary Figure 26 that the SWIF(r) signals in this region are not qualitatively different from the SWIF(r) signals we see throughout the genome, despite the fact that balancing selection is typically thought to be the primary mode of selection in the MHC^54^.

**Figure 3.**
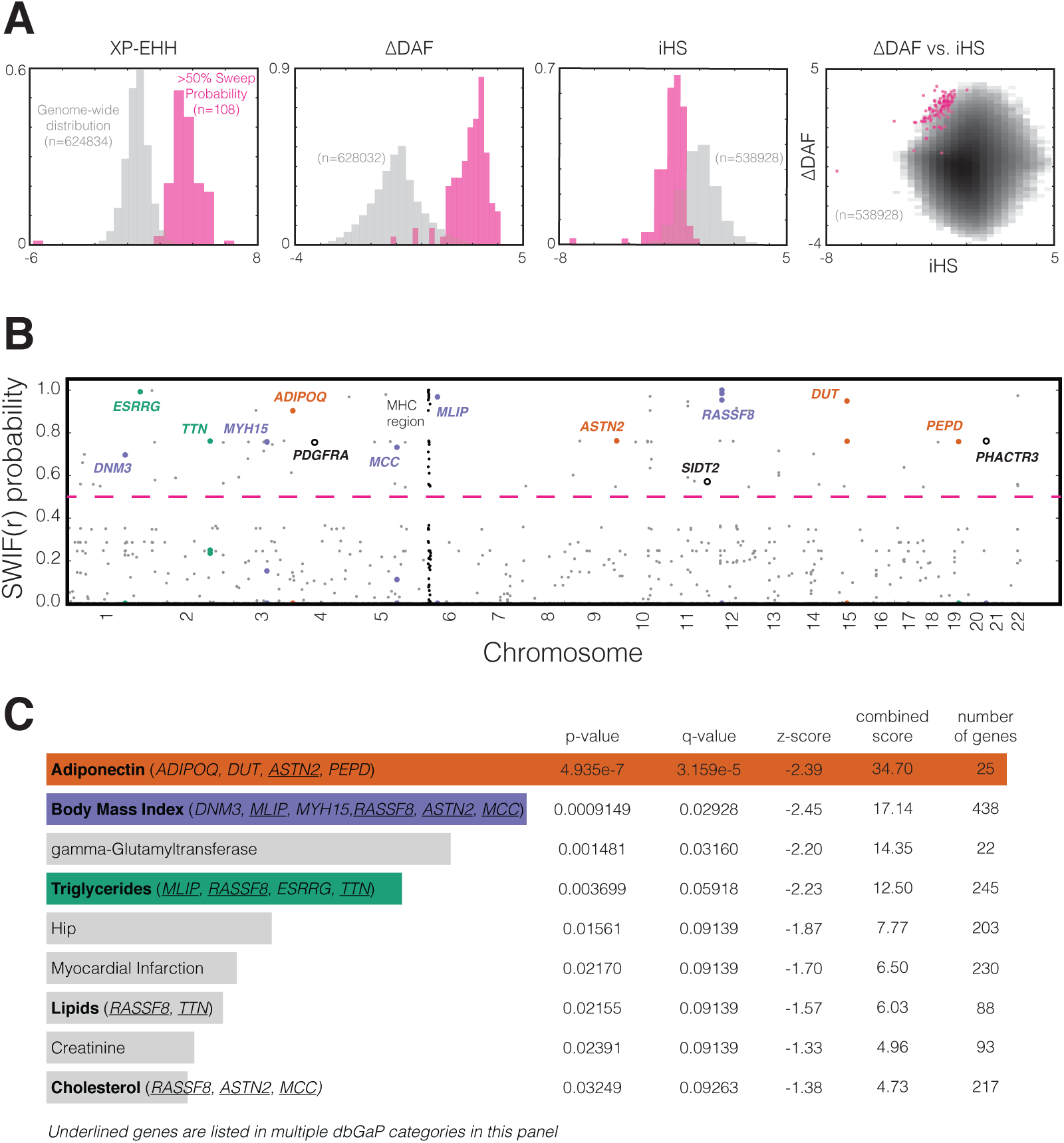
Genome-wide SWIF(r) scan for adaptation in ‡Khomani San SNP array data. **A)** The empirical genome-wide univariate distributions of three of the component statistics, XP-EHH, ADAF, and iHS are shown in gray, as is the empirical joint distribution of ADAF and iHS (where darker bins have more observations than lighter bins). The number of sites in each genome-wide univariate distribution differs due to some component statistics being undefined more often than others. In pink are the corresponding distributions for the 108 variants that SWIF(r) identifies as having posterior sweep probabilities of greater than 50% (variants above the dashed pink line in panel B). The full set of empirical univariate and joint distributions of component statistics for this scan can be found in Supplementary Figure 25. **B)** The value plotted for each position along the genome is the calibrated posterior probability of adaptation computed by SWIF(r) (per-site prior for a selective sweep is *π* = 10^−4^ to detect signals of relatively old sweeps given the high long-term *N_e_* of the ‡Khomani San); only SNPs with a calibrated posterior sweep probability greater than 1% are plotted and the horizontal line indicates a probability cutoff of 50%. A strong signal of adaptation over the Major Histocompatibility Complex on chromosome 6 is shown in black. Gene names are listed for genes previously associated with metabolism- and obesity-related traits (colors match categories in panel C; open circles denote genes of interest that are not in any category in C). **C)** We used gene set enrichment analysis tool Enrichr^55^ to identify categories that had an overrepresentation of genes containing SWIF(r) signals (Supplementary Table 7). We found multiple enriched dbGaP categories related to metabolism and obesity, including Adiponectin (a protein hormone that influences multiple metabolic processes, including glucose regulation and fatty acid oxidation), Body Mass Index, and Triglycerides. Genes in these categories containing SWIF(r) signals are listed next to category names. p-values, q-values, and the total number of genes are shown for each category, and categories are ranked by a combined score computed by Enrichr^55^, which combines p-value and z-score information and is shown to return optimal ranking^60^. Adiponectin, Body Mass Index, and gamma-Glutamyltransferase all have q-values below 5%.

We tested for a common functional or phenotypic basis among the 80 genes bearing SWIF(r) signals by conducting a gene ontology enrichment analysis across public databases with Enrichr^55^. We find that these genes are significantly enriched for dbGaP categories related to adiponectin, body mass index, and triglyceride phenotypes (Figure 3C, Table 1). Specifically, SNPs in genes related to adiponectin (*ADIPOQ, PEPD, DUT*, and *ASTN2*) have among the highest posterior sweep probabilities (all ≥ 75%). SWIF(r) also identified SNPs within three other genes (*PDGFRA, SIDT2*, and *PHACTR3*) that have previously been associated with obesity and metabolism phenotypes (Figure 3B, Table 1). Some of the genes highlighted in Figure 3B are also involved in muscle-based phenotypes (Supplementary Note), but here we focus on the substantial evidence supporting the association of the highlighted genes with obesity and metabolism phenotypes in prior GWA and functional studies (Table 1).

**Table 1.** Multiple published functional and association studies link genes identified by SWIF(r) to metabolism- and obesity-related phenotypes. The second column contains all variants within the genes listed that have posterior sweep probability ≥50% as calculated by SWIF(r). Column 3 shows the derived allele frequency (DAF) in the |Khomani San at the SNP in column 2, and column 4 shows the calibrated posterior sweep probability calculated by SWIF(r) at that site. For genome-wide association (GWA) studies, GWA *p*-values are given for the strongest SNP associations. Bold rsid indicates a result about the specific SNP identified by SWIF(r) in column 2. All genes highlighted in Figure 3B are included in this table except *MCC* and *MLIP*, for which additional associations to metabolism- and obesity-related phenotypes could not be found beyond the dbGaP categories in Figure 3C.

| Gene | SNP(s) | DAF | P(sweep) | Studies relating gene to metabolism-obesity phenotype | refs |
| --- | --- | --- | --- | --- | --- |
| *DNM3* | rs12121064 | 54% | 70% | associated with waist hip ratio in Europeans (rs1011731, GWA $p = 7.5 \times 10^{-9}$)<br>associated with waist circumference in Hispanic women (GWA replication $p = 1.5 \times 10^{-3}$)<br>associated with weight loss after gastric bypass surgery | 61<br>62<br>63 |
| *ESRRG* | rs11808388 | 62% | 99.2% | regulates adipocyte differentiation by modulating the expression of adipogenesis-related genes<br>candidate obesity-susceptibility gene based on epigenetic profile and association with BMI<br>may be involved in increasing the potential for energy expenditure in brown adipocytes<br>mediates hepatic gluconeogenesis<br>contributes toward maintenance of hepatic glucose homeostasis<br>necessary for metabolic maturation of pancreatic β-cells<br>significantly up-regulated under treatment with cholesterol drug fenofibrate | 64<br>65<br>66<br>67<br>68<br>69<br>70 |
| *TTN* | rs16866534 | 44% | 76% | isoform composition in cardiac tissue is regulated by insulin signaling, possibly contributing to altered diastolic function in diabetic cardiomyopathy | 71 |
| *MYH15* | rs3957559 | 49% | 76% | variant rs3900940, along with four other variants, contributes to elevated risk for coronary heart disease | 72 |
| *ADIPOQ* | rs6444174 | 56% | 90% | **rs6444174** associated with adiponectin levels in African American women<br>associated with plasma adiponectin levels in Europeans (rs17366568, GWA $p = 4.3 \times 10^{-24}$)<br>associated with plasma adiponectin levels in African Americans (rs4686807, GWA $p = 1.6 \times 10^{-11}$)<br>associated with plasma adiponectin levels in East Asians (rs822391, GWA $p = 1.6 \times 10^{-10}$)<br>associated with coronary heart disease, body mass index (BMI), childhood obesity, metabolic syndrome, and Type II Diabetes<br>serum adiponectin levels are associated with metabolic health and cardiovascular risk | 57<br>73, 74<br>75<br>75<br>57, 76–80<br>81, 82 |
| *PDGFRA* | rs4530695 | 64% | 75% | used in molecular biology as a marker for white adipocytes<br>controls pancreatic β-cell proliferation<br>plays a role in the link between obesity and inhibited placental development | 83<br>84<br>85 |
| *ASTN2* | rs16934033 | 50% | 76% | contributes to genetic variation of plasma triglyceride concentrations<br>marginally associated with childhood obesity in Hispanic individuals (GWA $p = 2.4 \times 10^{-6}$) | 86<br>87 |
| *SIDT2* | rs11605217 | 27% | 57% | important regulator of insulin secretion<br>mice without the gene are glucose intolerant and have decreased serum insulin<br>associated with triglyceride levels (rs1242229, GWA $p = 3.1 \times 10^{-20}$) | 88<br>89, 90<br>91 |
| *RASSF8* | rs16929850<br>rs16929965<br>rs2729646<br>rs956627 | 61%<br>64%<br>68%<br>64% | 95.3%<br>99.9%<br>98.4%<br>99.9% | expression is significantly altered by fasting in mice | 92 |
| *DUT* | rs11637235 | 33% | 76% | missense variant causes a syndrome characterized in part by early onset diabetes mellitus | 93 |
| *PEPD* | rs12975240 | 62% | 76% | associated with adiponectin levels in multiple populations (rs731839, rs4805885, rs8182584, rs889139, rs889140, GWA p-values between $1.1 \times 10^{-9}$ and $2.2 \times 10^{-13}$)<br>associated with Type II Diabetes (rs3786897, GWA $p = 1.3 \times 10^{-9}$)<br>associated with fasting insulin levels (rs731839, GWA $p = 5.1 \times 10^{-12}$)<br>associated with serum lipid levels<br>expression is modulated by n-3 fatty acids | 75, 94<br>95<br>96<br>97<br>98 |
| *PHACTR3* | rs1182507 | 54% | 76% | regulation in adipose tissue is BMI-dependent<br>candidate obesity gene based on epigenetic profile | 58<br>65 |

One variant that SWIF(r) identifies, rs6444174, has a calibrated sweep probability of 90%, driven by extreme values at this SNP in *F_ST_*, XP-EHH, and ΔDAF (Figure 3A; empirical *p*-values 4.4 × 10^−4^, 4.0 × 10^−4^, 5.5 × 10^−4^ respectively). This variant lies in *ADIPOQ*, which is expressed predominantly in adipose tissue^56^, and codes for adiponectin, a regulator of glucose and fatty acid metabolism. In a study of associations between *ADIPOQ* variants and adiponectin levels and obesity phenotypes in 2968 African American participants, rs6444174 was found to be associated with serum adiponectin levels in female participants (*p* = 6.15 × 10^−5^), and with body mass index in all normal-weight participants (*p* = 3.66 × 10^−4^). The allele at high frequency in the ‡Khomani individuals studied here corresponds to decreased adiponectin levels and increased BMI, respectively^57^.

#### Exome-based support for targets of selection identified by SWIF(r)

This SWIF(r) scan was performed using SNP array data ascertained from primarily Eurasian polymorphisms, a common feature of commercial SNP array platforms. Thus, the observed SWIF(r) signals are likely tagging haplotypes common in the ‡Khomani San, and may not themselves be causal polymorphisms. We examined high-coverage exome data^48^ within each gene to identify putatively functional mutations near the sites identified by SWIF(r) (see Supplementary Table 8 for full results). This allows us to identify variants not captured on SNP array platforms, including variants that are unique to the ‡Khomani San. We note that we did not include the MHC genes in this exome analysis, because of potential issues with mapping and phasing of exome sequence data in the MHC region. In *ADIPOQ,* we identify a missense mutation, rs13716447, for which the nearest SNP that is present on the SNP array is rs6444174 (less than 1kb away); rs6444174 has a calibrated SWIF(r) sweep probability of 90%, the highest in *ADIPOQ* (Figure 4). The missense T allele at rs13716447 is at high frequency in the ‡Khomani San relative to all other populations sequenced in the 1000 Genomes Project (27% vs. <0.5%; Figure 4). Furthermore, in the SImons Genome Diversity Project (SGDP), whose samples are drawn from 130 diverse and globally distributed human populations, only four copies of the missense allele at rs13716447 are found: two copies in a ‡Khomani San individual, and one copy each in a Namibian San individual and a Ju|’hoansi San individual. This SNP defines the two major haplogroups within the *ADIPOQ* gene in a median-joining haplotype network for the gene region (Supplementary Figure 27), providing some support for selection at this SNP.

**Figure 4.**
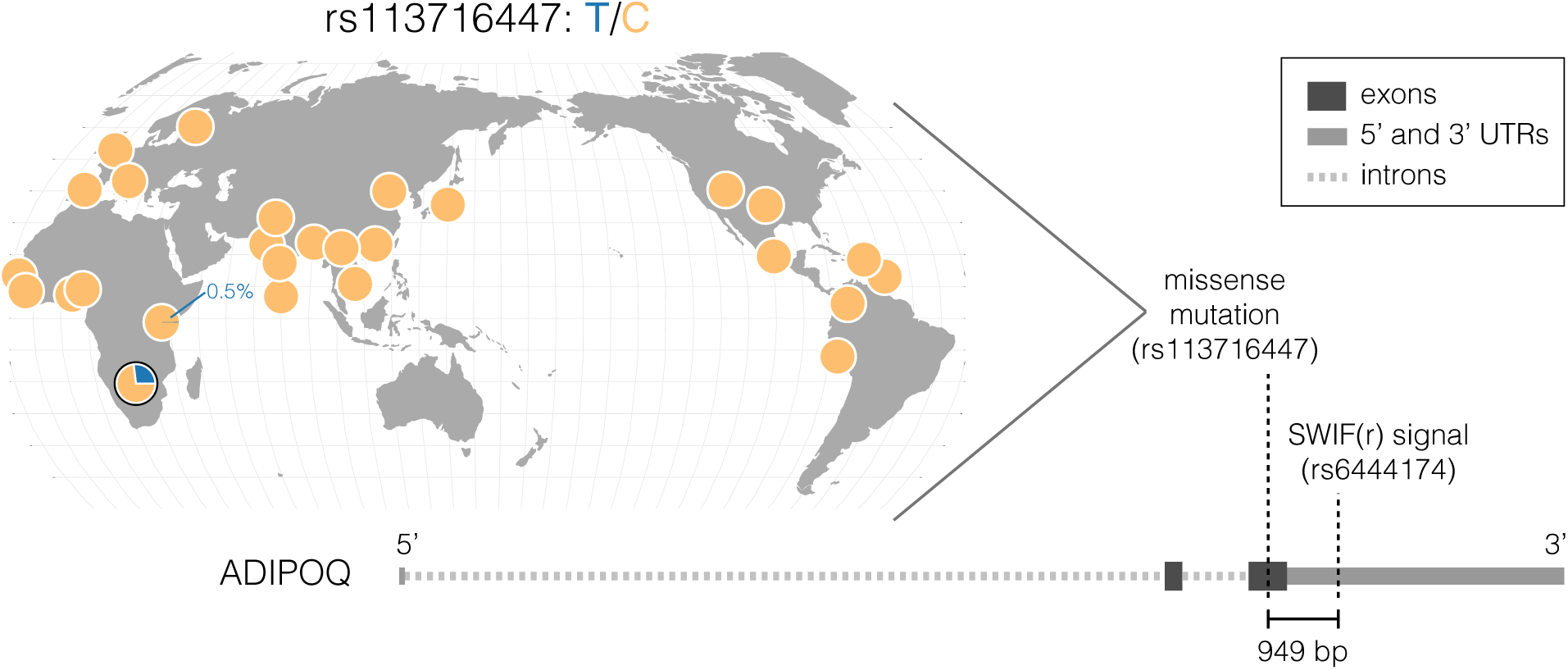
Exome analyses reveal potential beneficial mutations in genes identified by SWIF(r). Missense mutation rs1316447 is a potential causal mutation in *ADIPOQ*; A) Worldwide distribution of rs1316447 generated by the Geography of Genetic Variants Browser^99^ (http://popgen.uchicago.edu/ggv/) shows that the **T** allele carried by 27% of ‡Khomani San individuals (pie chart outlined in black) is extremely rare throughout phase 3 of the 1000 Genomes^35^, at a maximum of 0.5% in the Luhya population of Kenya. Diagram of *ADIPOQ* highlights the positions of the variant identified by SWIF(r) (rs6444174) and the nearby missense variant (rs1316447). These two variants are within 1kb of each other, suggesting that the SWIF(r) signal at rs6444174 is tagging this missense variant.

Two other genes highlighted in Figure 3B harbor promising polymorphisms that may be related to the underlying causal haplotypes. In *PEPD,* we identify a novel polymorphism at 10% frequency in the ‡Khomani (chr19:33882361) which is a missense mutation approximately 42kb from the SNP identified by SWIF(r). We also identify a missense mutation in the first exon of *PHACTR3* at 38% frequency in this sample, which is at < 2% frequency in other global populations including other Africans sequenced as part of the 1000 Genomes Project. Because the SNP array density is low, we expect that SWIF(r) signals in this population may in many cases be somewhat removed from the causal variants that these signals tag. We note that intronic variants in both *PEPD* and *PHACTR3* have been identified as cis-eQTLs that affect RNAseq expression in adipose tissue, in two independent northern European cohorts^58^.

For some of the genes identified by SWIF(r), exome data either was not generated, or did not reveal nearby functional polymorphisms with differential allele frequencies between the ‡KhomaniSan and other worldwide populations. One such gene is *RASSF8,* previously annotated as under positive selection in the Namibian and ‡Khomani San populations relative to western Africans using XP-EHH^59^; in our SNP array analysis, we detect a cluster of four SNPs in *RASSF8* within 70kb of each other, each with SWIF(r) sweep probability >98%. *RASSF8* is present in the BMI, Triglycerides, Lipids, and Cholesterol dbGaP categories (Figure 3C), yet functional mutations underlying this SWIF(r) signal remain elusive.

## Discussion

In this paper, we have presented both a new method for selective sweep detection, SWIF(r), and new insight into adaptive evolution in the ‡Khomani San. Not only does SWIF(r) outperform existing SNP-based component statistics and composite methods when detecting both complete and incomplete sweeps in simulation (Figure 1), it also localizes experimentally validated adaptive mutations *using genomic data alone* (Figure 2). SWIF(r) accounts for the confounding effect of neutral population histories when detecting sweeps by generating training simulations based on a demographic model, and we find it is robust to misspecification of the demographic model underlying the testing data (Supplementary Figure 13). We outline an algorithm for modeling SNP ascertainment in training simulations, thereby enabling the application of SWIF(r) to selection scans using genotype array data from diverse understudied populations like the ‡Khomani San (Online Methods). While some of the component statistics we use here may be fairly robust to ascertainment bias, this algorithm also enables future use of component statistics that are more vulnerable to ascertainment bias, such as SFS statistics, within SWIF(r)’s framework. When analyzing genotype and exome data from 45 ‡Khomani San individuals, we find that SWIF(r) signals tagging functional variants are enriched in genes associated with metabolism and obesity (Figure 3, Table 1, Figure 4).

Composite classification frameworks such as SWIF (r) quantitatively ground a common qualitative approach used in scans for adaptive sweeps based on summary statistics: evidence for selection at a locus is considered stronger when extreme values are observed for more than one statistic (Figure 3A). Furthermore, machine-learning approaches like SWIF(r) that incorporate joint distributions of selection statistics can detect sweep events that individual univariate statistics cannot (Supplementary Figure 28). SWIF(r) additionally reports calibrated probabilities assessing evidence for selective sweeps site-by-site, resulting in a transparent probabilistic framework for localizing adaptive mutations. These features allow for the localization of specific adaptive variants rather than adaptive regions, and minimize bias arising from undefined component statistics. While approaches such as Approximate Bayesian Computation can exploit higher-dimensional correlations in order to distinguish between selective sweep modes at candidate loci^100^, this comes at the cost of genome-scale tractability, and can be vulnerable to the curse of dimensionality^19^. The AODE framework allows us to transparently calculate probabilities without the need for imputation of undefined statistics, and our priors are made explicit, allowing for clearer interpretation. Future applications of SWIF(r) can easily incorporate new site-based summary statistics as they are developed (Supplementary Figure 29), and can assign variable site-specific prior sweep probabilities according to genomic annotations: for example, one could assign a smaller prior for synonymous variants relative to non-synonymous variants, or a higher prior in regulatory regions relative to intergenic regions^31^.

In order for the class probabilities reported by SWIF(r) to be practically interpretable, we calibrated SWIF(r), such that *k*% of variants with a posterior sweep probability of *k*% are indeed sweep variants. We have implemented a calibration scheme based on isotonic regression for SWIF(r) that maps the posterior sweep probabilities to their empirical sweep proportions in simulated data (Supplementary Figure 1, Supplementary Figure 2, Supplementary Figure 3), but importantly, this calibration relies on the composition of the training set used. While for some classifiers, the proportions of classes are known, or can be reliably estimated (e.g. see Durand *et al.*^101^ and Scheet and Stephensi^102^), the proportion of sites throughout the human genome that are adaptive is unknown. For calibrating SWIF(r), we chose training training sets made up overwhelmingly of neutral variants; while our calibration of SWIF(r) always preserves the rank order of posterior probabilities (Supplementary Figure 3), the specific choice of training set makeup can have a dramatic effect on the calibration. Therefore, a direct interpretation of the posterior probabilities reported by SWIF(r), or any other classifier that calculates probabilities, must incorporate knowledge of the scenarios used for training and calibration.

One caveat for interpretation of the SWIF(r) results presented here is that we train SWIF(r) on hard selective sweeps. In simulation, we find that SWIF(r) is also sensitive to sweeps from standing variation with a low initial frequency (Supplementary Figure 30); indeed, the sweep in West Africans in the gene *DARC,* for which SWIF(r) calculates a high sweep probability (Figure 2A) has recently been shown to have originated from standing variation in the ancestral population^103^. Given the multiple metabolism- and obesity-related sweep targets identified by SWIF(r) in the ‡Khomani San (Figure 3), we also suspect that some putative adaptive mutations identified by SWIF(r) may be components of polygenic adaptation.

Genes with SWIF(r) signals in our high-throughput genomic scan for selective sweeps in the ‡Khomani San have been independently identified in multiple GWA studies and functional experiments as associated with metabolism- and obesity-related phenotypes (Table 1). One way to interpret this signal is through the lens of the “thrifty gene” hypothesis, which posits that ready fat storage was positively selected for in hunter-gatherer populations due to the survival advantage it conferred in unreliable food cyclesi^104^. The hypothesis further states that modern disease phenotypes such as type 2 diabetes and obesity are the consequence of a radical shift in diet from ancestral environments and forager subsistence strategies to a contemporary environment with abundant food in the form of simple sugars, starches, and high fat, though this is a subject of much debate^105,106^. Although most indigenous Khoe and San groups of the Kalahari are classically considered small and thin, populations such as the Khoekhoe cow/goat pastoralists are characterized by steatopygia (i.e. extensive fat accumulation along the buttocks and thighs in women), as notoriously described by early European explorers and anthropologists^107–109^. While the thrifty gene hypothesis would predict an increase in metabolic pathology for these individuals, studies have shown that accumulated subcutaneous gluteofemoral fat, found in patients exhibiting steatopygia^110,111^, is protective against diabetes and other metabolic disordersi^112,113^. The mutations and genes identified by our SWIF(r) scan, such as *ADIPOQ,* are natural targets for functional assays to determine the origins and consequences of subcutaneous versus visceral fat; future studies could merge such assays with phenotypic data on diabetes and metabolic syndromes in KhoeSan groups to gain new insight into the “obesity-mortality paradox”^114^.

In this selection scan, we also see an abundance of SWIF(r) signals in the MHC region involved in immunity. It is possible that this signal reflects balancing selection, which is the mode of selection canonically thought to be occurring within this region^54^. Indeed, it has been shown that the signatures of balancing selection and incomplete or recurrent sweeps may be similar to signatures of positive selection^115,116^. We note, however, that other studies using different methodologies have detected signatures of directional selection in the MHC in human populations^117,118^, and others have noted that fluctuating directional selection is a possible mechanism for pathogen-mediated selection in this region^119^.

The probabilistic framework of SWIF(r) suggests two natural extensions for future applications. First, while the use of SNP-based component statistics enabled us to localize adaptive mutations, SWIF(r) could easily incorporate region-based component statistics, including composite likelihood approaches like XP-CLR^120^, and SFS-based measures^3,121^ in order to help detect older selection events, for which haplotype-based statistics are less powerful. Second, future studies can exploit the flexibility and interpretability of SWIF(r) to conduct multi-class classification. Supplementary Figure 31 illustrates a preliminary extension of SWIF(r) that classifies sweeps based on the start time of positive selection. Recent methods have attempted multi-class sweep classification using hierarchical binary classification or other machine learning approaches^14,18,22^, but without the benefits of a transparent probabilistic framework in which priors are made explicit. Using the probabilistic framework of SWIF(r), future studies could determine the mode of adaptive evolution at genomic sites, including background selection or sweeps from standing variation or recurrent mutation^21,22^, or infer the timing or selective strength of an adaptive event^100^. Thus, SWIF(r) offers a technical advance in genome-wide sweep detection that can yield new insight into the modes and roles of selection in shaping population-genomic diversity.

## Online Methods

### SImulation of haplotypes for 1000 Genomes analysis

SImulations based on the demographic model of African, Asian, and European populations outlined in Schaffner *et al.*^27^ were carried out with the following alterations necessary for allowing the simulation of recent selective sweeps (within the last 30ky): no modern population growth (within the last 30 generations), and migration ending 500 generations following the Asian/European split instead of continuing to the present. We carried out 100 simulations of 1Mb regions from this neutral demographic model using `cosi`, which resulted in ∼400,000 neutral training points. We generated a new recombination map for each simulation with the recosim package within `cosi`, using a hierarchical recombination model that assumes a regional rate drawn from the observed distribution of rates in the deCODE genetic map^122^, and then randomly generates recombination hotspots with randomly drawn local rates^27^. For each simulation, we generated 120 1Mb-long haplotypes from each of the three populations.

Selective sweeps continued until the time of sampling, and were simulated for a range of sweep parameters: start time ranging over [5, 10, 15, 20, 25, 30kya], final allele frequency ranging over [0.2, 0.4, 0.6, 0.8, 1.0], and population of origin ranging over [African, Asian, European]. Note that these sweeps cover a range of incomplete as well as complete sweeps in a population of interest. The selection coefficients for each parameter set are fully determined by the effective population size, sweep start time, and final allele frequency, and are displayed in Supplementary Table 5. We calculated these selection coefficients using Equation 4 for complete sweeps (by which we mean sweeps where the beneficial mutation has reached fixation in the population of interest), and Equation 5 for incomplete sweeps, where *t*_1_ is the sweep start time and *t*_2_ is the sweep end time, both measured in generations from the present, *N_e_* is the effective population size, 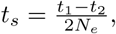 and *ϕ* is the present-day frequency of the beneficial allele, 0 < *ϕ* ≤ 1^123^. The range of selection coefficients corresponds to a range of *α* = 2*Ns* of ∼100-4500. For each set of sweep parameters, we carried out 100 simulations with the adaptive allele located halfway along the 1MB region, for a total of 9,000 sweep training points.

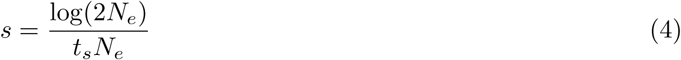

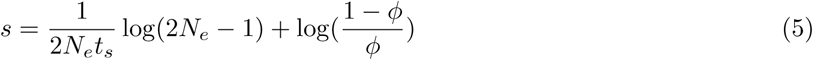

### SImulation of haplotypes for ‡Khomani San analysis

To train our classifier to identify selective sweeps in the ‡Khomani San, we used the demographic model inferred by Gronau *et al*.^28^ based on six diploid whole-genome sequences, one from each of six populations: European, Yoruban, Han Chinese, Korean, Bantu, and San. We used the inferred population sizes, coalescent times, and migration rates reported by Gronau *et al.^28^,* which are calibrated based on a 6.5 million year human-chimpanzee divergence, the presence of migration between the Yoruban and San populations, and 25 years/generation to construct a demographic model for the Han Chinese, European, Yoruban, and San, shown in Supplementary Figure 21.

Because Uren *et al.*^45^ showed that the ‡Khomani San have experienced recent gene flow from both Western Africa and Europe, we replaced migration in the Gronau model with two pulses of recent migration: one pulse from the Yoruban population with migration rate 0.179 at 7 generations ago, and one pulse from the European population with migration rate 0.227 at 14 generations ago. We found that these rates resulted in present-day admixture levels that matched those found in Uren *et al.*^45^ (Supplementary Note).

Using `cosi`^27^, we simulated 1Mb genomic regions, comprising both neutral and sweep scenarios as described earlier, with sample sizes matching the number of individuals in the filtered 1000 Genomes dataset, and the number of San individuals in our study.

Because the ‡Khomani have been isolated for so long, we included additional sweep scenarios, with sweeps beginning and ending between 30 and 60kya. We called these “old sweeps” and trained the classifiers on three classes: neutral, “old sweeps”, and “recent sweeps” (those occurring within the last 30ky). SInce we found that the classifiers did not have enough power to reliably distinguish between old and recent sweeps (Supplementary Figure 31), in applications to data, we only considered the total probability of a sweep, given by the sum of the posterior probabilities for each sweep class.

### SImulation of haplotypes including recent population growth

To test SWIF(r)’s robustness to misspecification of recent population growth, we implemented a set of simulations using a demographic model from Gravel *et al.*^29^ that estimates recent exponential population expansions with rates of 0.38% for Europe and 0.48% for East Asia over the last 23,000 years. SInce the simulation software `cosi`^27^ cannot simulate sweeps and population-level changes simultaneously, we allowed the expansion to last from 23,0000 years ago to 5,000 years ago, to allow for sweeps beginning 5,000 years ago. We also included the migration rates inferred by Gravel *et al.*^29^ between Europe, East Asia, and Africa, and between Africa and the ancestral population of Europe and East Asia. As in other analyses, We simulated selective sweeps spanning a range of present-day allele frequencies from 20% to 100%.

### SImulation of background selection

For evaluating SWIF(r)’s robustness to background selection, we generated 3 sets of simulations of 1Mb each using forward simulator `slim`^30^: neutral regions, regions with a hard sweep, and genic regions. For genic regions, we followed Messer and Petrov^21^ to simulate gene structure: each simulation had one gene with 8 exons of 150bp each, separated by introns of 1.5kb, and flanked by a 550bp 5′UTR and a 250bp 3′UTR. Within exons and UTRs, 75% of sites were assumed to be functional. Mutations were assumed to be codominants, and fitness effects across different sites were assumed to be additive. Functional sites were divided into 40% “strongly deleterious” sites with selection coefficient −0.1, and 60% “weakly deleterious” sites with selection coefficients between −0.01 and −0.0001. The mutation rate was set at 2.5 × 10^−8^ per site per generation, and the recombination rate at 10^−8^. Note that for testing the robustness of SWIF(r), we only considered sites from these simulations that landed in exons or UTRs.

For all three sets of simulations, we simulated two populations with *N_e_* = 5000, which split from each other 40,000 years ago. For sweep simulations, we drew selection coefficients for the beneficial allele from an exponential distribution with mean 0.03, and sweeps begin 10,000 years ago. SInce forward simulations are much more computationally intensive than coalescent simulations, we rescaled our parameters by a factor of 10 (10 times larger for mutation rate, recombination rate, and selection coefficients, 10 times smaller for population sizes and 10 times shorter for all times) to make the simulations feasible^30^.

### Implementation of the Classifiers

For ease of comparison, we built SWIF(r) using the same statistics that comprise the Composite of Multiple SIgnals (CMS)^13,16^: the fixation index (*F_ST_*), cross-population extended haplotype homozygosity (XP-EHH) (adapted for improved performance on incomplete sweeps; Supplementary Note), the integrated haplotype score (iHS), and change in derived allele frequency (ΔDAF). AiHH was excluded in applications to real data 469 because of non-normality (Supplementary Note, Supplementary Figure 5).

#### Implementation of SWIF(r)

To avoid over-fitting the joint distributions modeled in the AODE framework (Equation 3), we fit Gaussian mixture models with full covariance matrices (i.e. containing nonzero off-diagonal entries) to the joint probability distributions of each pair of statistics *S_i_* and *S_j_* within each scenario C (neutral or sweep), *P*(*S_i_* = *si*, *S_j_* = *si*|*C*), with the number of components ranging between three and five based on Bayesian Information Criterion (BIC) curves. Joint probabilities were learned using sites for which both component statistics were defined. We used the python package `scikit-learn`^124^ to compute the BIC curves and fit the mixture models. These mixture models capture the salient features of each pairwise joint distribution, as illustrated in the example in Supplementary Figure 32. Given the smoothed joint distribution learned for a pair of statistics (*S_i_*, *S_j_*), we calculate the conditional probability distributions *S_j_*|*s_i_* as one-dimensional gaussian mixtures:

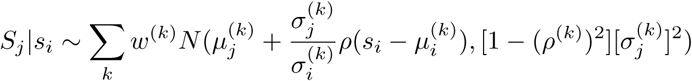

where *N*(*μ*, *σ*^2^) denotes the normal distribution with mean *μ* and variance *σ*^2^, *k* indexes the components in the joint Gaussian mixture, *w*^(*k*)^ is the weight assigned to each component, *μ_i_* and *μ_j_* are the components of the joint mean, and *σ_i_*, *σ_j_*, and *ρ* are taken from the joint covariance matrix Σ.

We find that SWIF(r) loses little power to identify sites with adaptive mutations when one of the component statistics is undefined, but each component statistic differentially influences the power of SWIF(r) (Supplementary Figure 33).

#### Calibration of SWIF(r) probabilities

There are a few techniques for calibrating probabilities returned by a binary classifier so that of all of the data points that are given a *k*% probability of belonging to class A by the classifier, *k*% of those are indeed drawn from class A, and (100 – *k*)% are drawn from class B^25,125^. Isotonic regression (IR) is a popular method because it makes no assumptions about the mapping function beyond requiring that it be monotonically increasing^126^. In the case of SWIF(r) probabilities, IR calibration works by grouping sweep and neutral variants from a training dataset into posterior probability bins, and mapping each bin to the empirical proportion of variants in the bin that are sweep variants. We used 10 bins for calibration, because we found that using more bins increased the risk of overfitting. This can be mitigated by performing more simulations, but in our case, even with 1000 neutral simulations of 1Mb each, mid to high posterior probabilities were extremely rare at neutral variants. For sweep site localization in data from the 1000 Genomes Project, we calibrated SWIF(r) based on a training dataset composed of 99.99% simulated neutral variants and 0.01% simulated sweep variants, and for application to the ‡Khomani San SNP array data, we calibrated SWIF(r) based on a dataset composed of 99.95% simulated neutral variants and 0.05% simulated sweep variants (Supplementary Figure 1, Supplementary Figure 2, Supplementary Figure 3). The slightly larger fraction of sweep simulations in the ‡Khomani San training set relative to the 1000 Genomes training set allowed for more sensitivity to older sweeps, and accounted for the sparser SNP density of this dataset. In both data, we restricted the simulated sweep variants to those with present-day allele frequencies over 50%, since we have the most power in this realm (Supplementary Figure 6), and wanted to avoid overcorrection of strong signals.

A downside to IR is that by its nature, it maps a range of input values to the same output value, which removes some information about which probabilities are larger than others. We implemented a “smoothed” isotonic regression for calibration that interpolates the piecewise constant mapping function learned by IR (Supplementary Figure 3). In practice, we find that both methods of calibration produce equally well-calibrated classifiers; that is, after either method, the data points in our simulated dataset that have a calibrated posterior sweep probability of *k*% are made up of approximately *k*% sweep simulations and (100 – *k*)% neutral simulations (Supplementary Figure 1, Supplementary Figure 2). Unlike IR alone, however, smoothed isotonic regression has the advantage of preserving strict monotonicity of posterior probabilities.

#### Implementation of CMS

We implemented CMS following the algorithm described in Grossman *et al.*^13^ and personal communication with the authors. Based on the simulations and component statistics described above, CMS is computed as the product of individual posterior distributions:

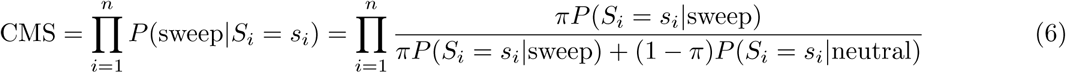

where *π*, the prior probability of a sweep, is 10^−6^.

When one or more component statistics are undefined at a locus, CMS is not well-defined. If statistics are simply left out of the product, this artificially inflates the reported score. Some compensation is thus required to avoid such a bias, which is not discussed by Grossman *et al*. We implemented a conservative compensation scheme: if statistic *S_i_* is undefined at a locus, we set its value to the mean of the distribution for that statistic learned from neutral simulations.

For the purpose of evaluating CMS using YRI, CEU, and CHB+JPT samples from the 1000 Genomes, we use CMS Viewer (https://pubs.broadinstitute.org/mpg/cmsviewer/; use date: 04/26/2016), an interactive tool designed by Grossman *et al.*^16^ for visualizing genome-wide CMS scores.

### Implementation of window-based methods

We implemented evolBoosting using the R package released by Lin *et al.*^14^ (http://www.picb.ac.cn/evolgen/softwares/) using default settings. We trained and tested evolBoosting on the simulations of YRI, CEU, and CHB+JPT described above, including all sweep durations and present-day allele frequencies, splitting the simulations in two equally sized groups for training and testing. We used the middle 40kb of each 1Mb simulation, and generated window-based SWIF(r) probabilities by taking the maximum posterior sweep probability for all SNPs within the 40kb window.

For evoNet^19^ we used the same simulations as above, but using the central 100kb windows of each 1Mb simulation (following Sheehan *et al.*^19^). The software released by Sheehan *et al.*^19^ is generalized, so that any component statistics may be used for implementing the deep learning framework. Therefore, we implemented evoNet using the component statistics we use here for SWIF(r): *F_ST_*, XP-EHH, iHS and ΔDAF. evoNet exploits the signatures at three distances from the beneficial allele (0-10kb, 10-30kb, and 30-50kb away), so we used the mean values for each component statistic in each of these windows, for a total of 12 component statistics. For this comparison, we generated window-based SWIF(r) probabilities by taking the maximum posterior sweep probability for all SNPs within the 100kb window.

### ROC analysis

To generate the ROC curves for CMS, SweepFinder, and the component statistics (Figure 1A-C), we varied the threshold for classifying a mutation as adaptive in order to cover the range from ∼0% false positive rate to ∼100% true positive rate. For SWIF(r), and the ODEs, we varied the prior *π*, and sites with scores greater than 0.5 were classified as adaptive (Supplementary Figure 4). To generate Figure 1D, we partitioned all simulations by present-day frequency of the adaptive mutation and sweep start time. For each pair of these parameters, we approximated the area under the ROC curves (AUROC) by summing the areas of the trapezoids defined by each pair of neighboring points in the ROC plane, then identified the summary statistics with the highest and second-highest AUROC. ROC curves for window-based SWIF(r), evoNet, and evolBoosting, were generated in much the same way, except that we varied the threshold for classifying a window as containing an adaptive variant.

### Ascertainment modeling

For our selection scan in the ‡Khomani San population, we use genotype data from two SNP arrays^45^; the ascertainment bias of these arrays means that the simulated haplotypes we generate from the four populations (‡Khomani San, YRI, CEU, CHB+JPT) for training SWIF(r) differ dramatically from the observed data for these populations at the sites genotyped on the arrays. To account for this, we implemented an ascertainment-modeling algorithm that prunes sites from simulated haplotypes in order to provide SWIF(r) with simulations for training that match the site frequency spectrum (SFS) of the observed data as closely as possible. The key to this algorithm is to define regions of joint SFS space that are similar in terms of representation on the SNP arrays (e.g. SNPs with low derived allele frequency in all populations are fairly common, while SNPs that are highly differentiated across multiple populations are relatively rare). Defining these “equivalence classes” (hereafter referred to as “SFS regions”) in joint SFS space allows us to learn the density of SNPs from each SFS region along the SNP arrays, and then to thin simulations in order to re-create those densities. This first requires smoothing of the joint SFS to account for sparsity. The full algorithm is as follows:

1. Learn the empirical 3D SFS for YRI, CEU, and CHB+JPT individuals in the 1000 Genomes Project, restricted to SNPs present in the overlap between the Illumina OmniExpress and OmniExpressPlus platforms (Supplementary Note) This results in a three-dimensional array of SNP counts for each triplet of derived allele frequencies (DAF_YRI_, DAF_CEU_, DAF_CHB+JPT_). For this dataset, given 87 YRI individuals, 81 CEU individuals, and 186 CHB and JPT individuals, the dimensions of this three-dimensional array are 175 × 165 × 373 (2*n* + 1 in each dimension for n individuals).
2. To account for sparseness in the empirical 3D SFS, subdivide each axis into 40 evenly-spaced bins to create a new 40 × 40 × 40 array where each entry is the average SNP count within that 3-dimensional bin; this array approximates the original empirical 3D SFS. Use the one-dimensional histogram of average SNP counts across all 40^3^ bins to define five intervals that span the range of counts, then assign each bin to its interval (Supplementary Figure 34). Groups of bins belonging to the same interval will be hereafter referred to as “SFS regions.” We note that we choose a 40 × 40 × 40 array for smoothing because it resulted in SFS regions with well-defined boundaries in 3-dimensional space (Supplementary Figure 34); these dimensions may need to be altered for other datasets to achieve well-defined boundaries as in Supplementary Figure 34.
  a. In most SFS regions, the SNP counts in the 3-dimensional SFS are relatively invariant; however, in the SFS region with the highest SNP counts (the region in red in Supplementary Figure 34, corresponding predominantly to SNPs with low derived allele frequency in all populations), there is a wide range of SNP counts (this is analogous to the higher variability in counts of low-frequency variants in the 1-dimensional site frequency spectrum relative to that of medium- and high-frequency variants). To account for this increased variability, apply a similar procedure as above: subdivide each bin in the highest SFS region by 2 in each dimension (resulting in 8 sub-bins), re-learn the average SNP count within that sub-bin, use a histogram of average SNP counts across sub-bins to again define five intervals, and assign each sub-bin to its interval, thereby defining an additional set of SFS regions that gives better resolution in higher-density areas.
3. For each 1Mb block along the SNP array, count the number of SNPs that fall in each SFS region, based on the observed derived allele frequencies at each SNP for YRI, CEU, and CHB+JPT. This provides a measure of SNP density (counts per Mb) for each SFS region. Applying this over a sliding window of 1Mb across the entire SNP array results in a distribution of densities for each SFS region.
4. Within each 1Mb block of simulated sequence data, assign each simulated SNP to its SFS region. For each SFS region, draw a value from the distribution of SNP densities learned in step 3, then randomly down-sample the number of simulated SNPs that fall in that region to match this value. In the rare case in which downsampling is not possible for a given SFS region (i.e. there are fewer simulated SNPs in that region than the value drawn from the distribution of densities), retain all simulated SNPs that belong to the SFS region.
5. For training the classifiers, restrict the simulated ‡Khomani San genotype data (as well as the simulated data from the 1000 Genomes populations) to the downsampled set of SNPs.

## Software and Data availability

SWIF(r) repository: https://github.com/ramachandran-lab/SWIFr; selscan repository: https://github.com/szpiech/selscan 1000 Genomes phase 1 data:
ftp://ftp.1000genomes.ebi.ac.uk/vol1/ftp/phase1/analysis_results/integrated_call_sets/.

‡Khomani San genotype data were first described by Uren *et al.*^45^, and ‡Khomani San exome data were first described by Martin *et al.*^48^. Queries regarding access to ‡Khomani San data analyzed here should be sent to the South African San Council for research and ethics review by contacting both Leana Snyders and.

## Acknowledgments

We thank Dean Bobo, Barbara Engelhardt, Chris Gignoux, David Guertin, Erik Sudderth, Zachary Szpiech, Jeremy Mumford, Lorin Crawford, Paul Norman, Sara Mathieson, and the Ramachandran Lab for helpful discussions; we also thank Shari Grossman, Ilya Shlyakhter, and Pardis Sabeti for discussing details regarding the implementation of CMS. We are grateful to Ryan Hernandez for multiple discussions regarding analysis of exome sequences and false discovery estimates, as well as to three reviewers for multiple suggestions that improved the manuscript. This research was supported by the Pew Charitable Trusts (S Ramachandran is a Pew Scholar in the Biomedical Sciences), and US National Institutes of Health (NIH) grant R01GM118652 (to S Ramachandran) and COBRE award P20GM109035. S Ramachandran is an Alfred P. Sloan Research Fellow and also acknowledges support from National Science Foundation (NSF) CAREER Award DBI-1452622. S Rong is supported by an NSF Graduate Research Fellowship. APF was supported by an REU supplement to NSF CAREER Award DBI-1452622. EGA was supported by NIH grant K12-GM-102778.

## Author Contributions

L.A.S. and S. Ramachandran conceived the study and L.A.S. implemented the methods. Sequence data from the ‡Khomani San were generated and processed by E.G.A. and B.M.H., and L.A.S., E.G.A., B.M.H., and S. Ramachandran contributed to analysis of SWIF(r) results. A.P.F. annotated SWIF(r) results and contributed to comparison of SWIF(r) with existing methods. S. Rong contributed simulations of different modes of selection. L.A.S, E.G.A., B.M.H., and S. Ramachandran wrote the manuscript.

---

**Supplementary Figure 1.**
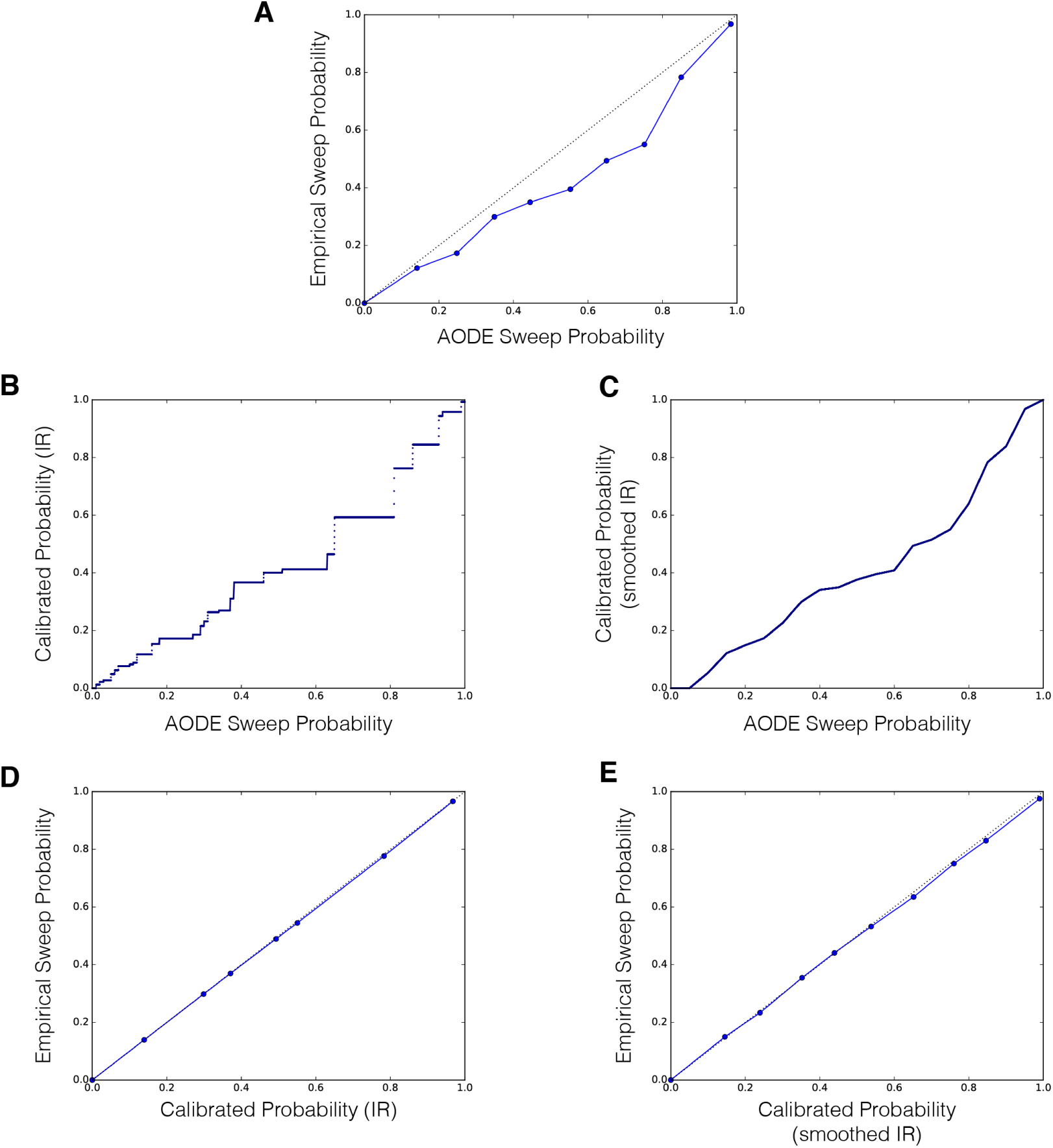
Calibration of SWIF(r) for analysis of data from the 1000 Genomes. For sweep site localization in data from the 1000 Genomes Project, we calibrated SWIF(r) based on a training dataset made up of 99.99% simulated neutral variants and 0.01% simulated sweep variants. We restricted the simulated sweep variants to those with present-day allele frequencies over 50%, since we have the most power in this realm, and wanted to avoid overcorrection of strong signals. **A)** Reliability curve^1^ for uncalibrated SWIF(r) posterior probabilities using 10 evenly-spaced bins between 0 and 1; the x-axis plots the mean posterior probability within each bin, and the y-axis plots the fraction of sites within the bin that are sweep variants (“empirical sweep probability”). Uncalibrated, the probabilities that SWIF(r) calculates are slightly inflated. **B)** Isotonic regression (IR) mapping of probabilities in each bin to their corresponding empirical sweep probabilities, based on panel A. **C)** Our smoothed IR map, learned by interpolating between the midpoints of the piecewise constant segments inferred by isotonic regression in panel B (see also Supplementary Figure 3). **D)** Reliability curve for sweep probabilities calibrated with isotonic regression from panel B. **E)** Reliability curve for sweep probabilities calibrated with smoothed isotonic regression from panel C.

**Supplementary Figure 2.**
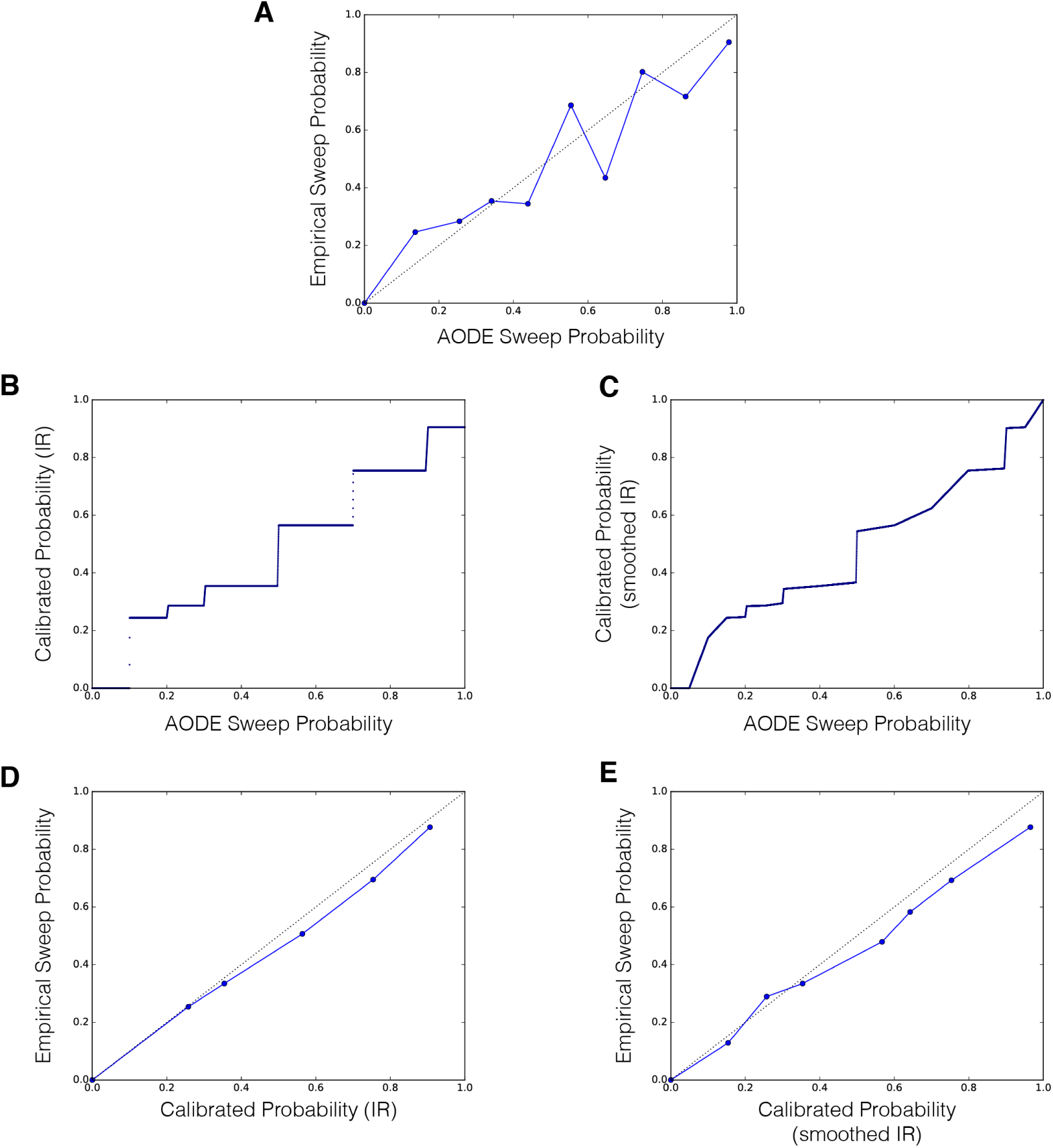
Calibration of SWIF(r) for analysis of SNP array data from the ‡Khomani San. For sweep site localization in array data from the ‡Khomani San, we calibrated SWIF(r) based on a training dataset made up of 99.95% simulated neutral variants and 0.05% simulated sweep variants with present-day allele frequencies over 50%. The slightly larger fraction of sweep simulations relative to the 1000 Genomes calibration is to allow for more sensitivity to older sweeps, and to account for the sparser SNP density of this dataset compared to the 1000 Genomes (phase 1). **A)** Reliability curve^1^ for uncalibrated SWIF(r) posterior probabilities using 10 evenly-spaced bins between 0 and 1; the x-axis plots the mean posterior probability within each bin, and the y-axis plots the fraction of sites within the bin that are sweep variants (“empirical sweep probability”). **B)** Isotonic regression (IR) mapping of probabilities in each bin to their corresponding empirical sweep probabilities, from panel A. **C)** Our smoothed isotonic regression map, learned by interpolating between the midpoints of the piecewise constant segments inferred by isotonic regression in panel B (see also Supplementary Figure 3). **D)** Reliability curve for sweep probabilities calibrated with isotonic regression from panel B. **E)** Reliability curve for sweep probabilities calibrated with smoothed isotonic regression from panel C.

**Supplementary Figure 3.**
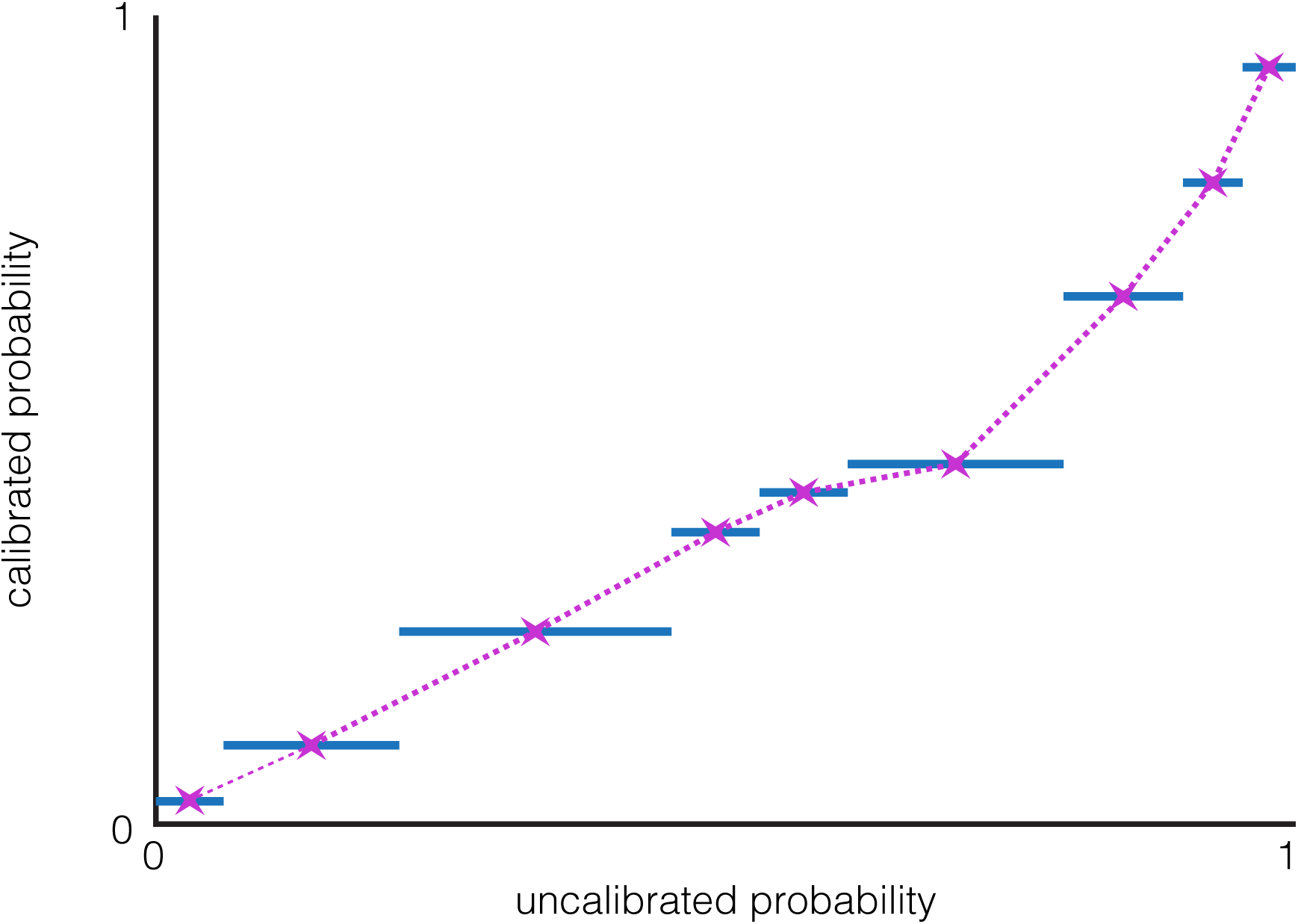
Schematic of smoothed isotonic regression for probability calibration. There are a few techniques for calibrating probabilities returned by a classifier so that of all of the data points that are given a k% probability of belonging to class *A* by the classifier, *k*% of those are indeed drawn from class *A*. Isotonic regression is a popular method because it makes no assumptions about the mapping function beyond requiring that it be monotonically increasing^2^. A downside, however, is that by nature, isotonic regression maps a range of input values to the same output value, which removes some information about which probabilities were larger than others. We implemented a “smoothed” isotonic regression for calibration that takes the piecewise constant mapping produced by isotonic calibration (solid blue line segments), and creates a mapping that preserves the strict monotonicity of the input data (purple dotted line). To obtain this new mapping, we interpolate between the midpoints (purple stars) of each piecewise constant segment. In practice, we find that both methods of calibration produce equally well-calibrated classifiers; that is, after either calibration method, the data points in our simulated dataset that have a calibrated posterior sweep probability of *k*% are made up of approximately *k*% sweep simulations and (100 – *k*)% neutral simulations (see Supplementary Figure 1D-E, Supplementary Figure 2D-E). However, smoothed isotonic regression has the advantage of preserving strict monotonicity of posterior probabilities.

**Supplementary Figure 4.**
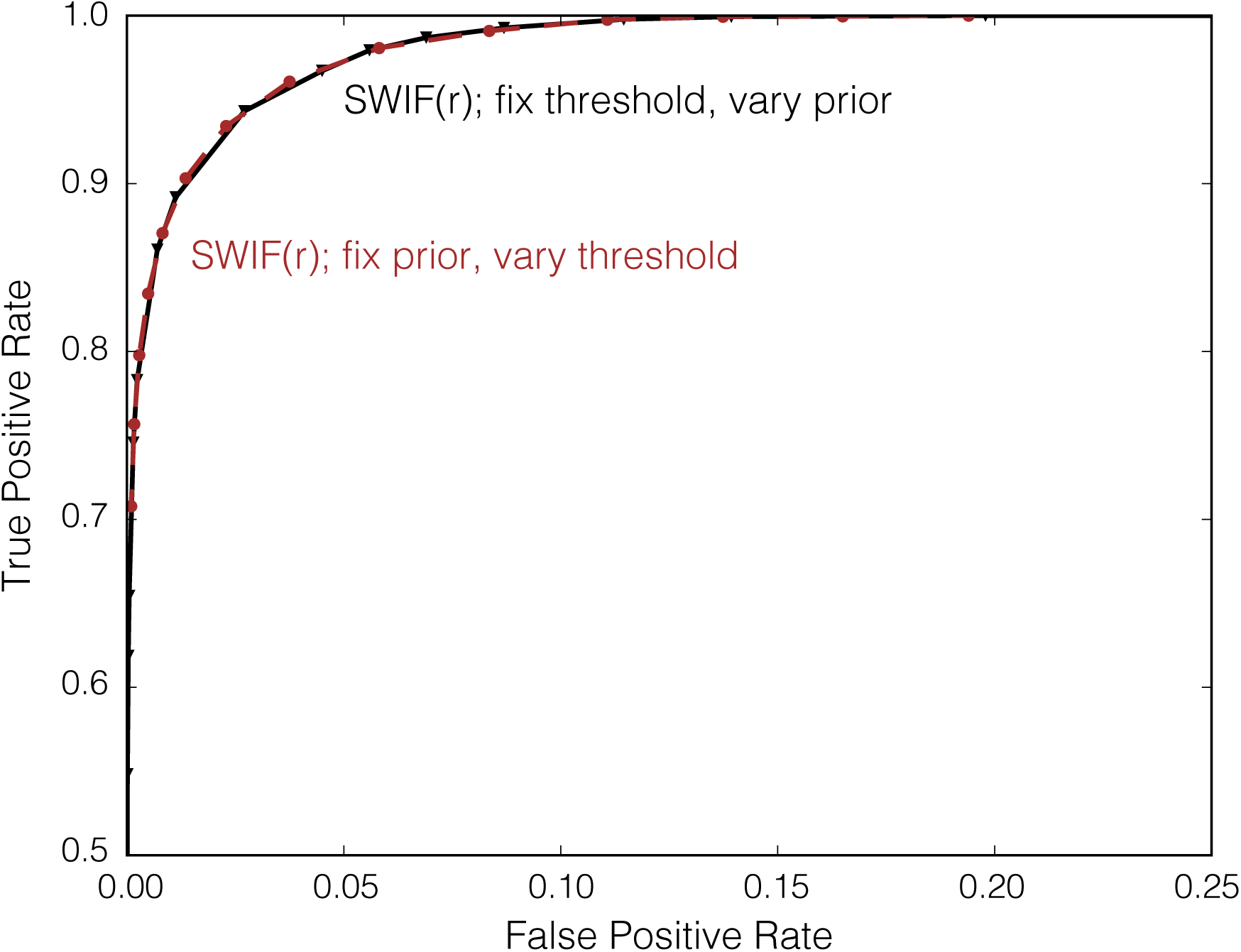
ROC curves for SWIF(r), varying the prior and varying thresholds. We plot ROC curves for SWIF(r) by fixing the threshold for classifying a site as adaptive at 50% posterior probability and varying the prior in Equation 3 (black curve with points denoted as triangles). However, an equally valid method would be to fix the prior and vary the posterior probability cutoff for determining which sites are classified as neutral and which are classified as adaptive (brown curve with circular points, prior fixed at 10^−5^). We show below that both methods result in equivalent ROC curves.

**Supplementary Figure 5.**
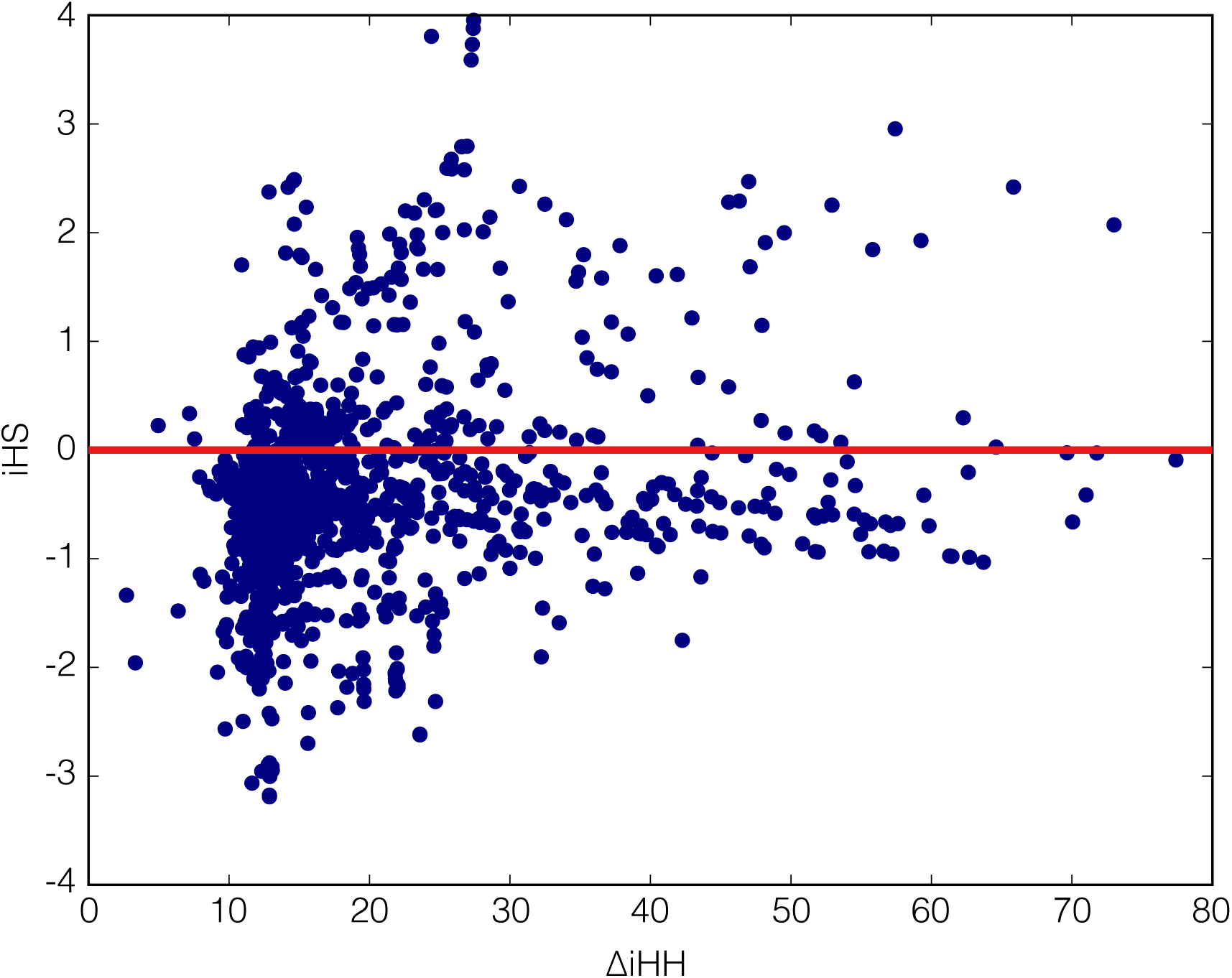
ΔiHH produces many false positives in 1000 Genomes data. Points represent ΔiHH and iHS values for sites classified as adaptive mutations (sweep probability ≥ 50%) in the 1000 Genomes data (here, ΔiHH included as a component statistic in SWIF(r)). The red line denotes iHS = 0, to emphasize the large number of these sites for which ΔiHS is positive. SInce positive values of iHS provide evidence *against* positive selection for the derived allele, these sites are likely false positives driven by ΔiHH.

**Supplementary Figure 6.**
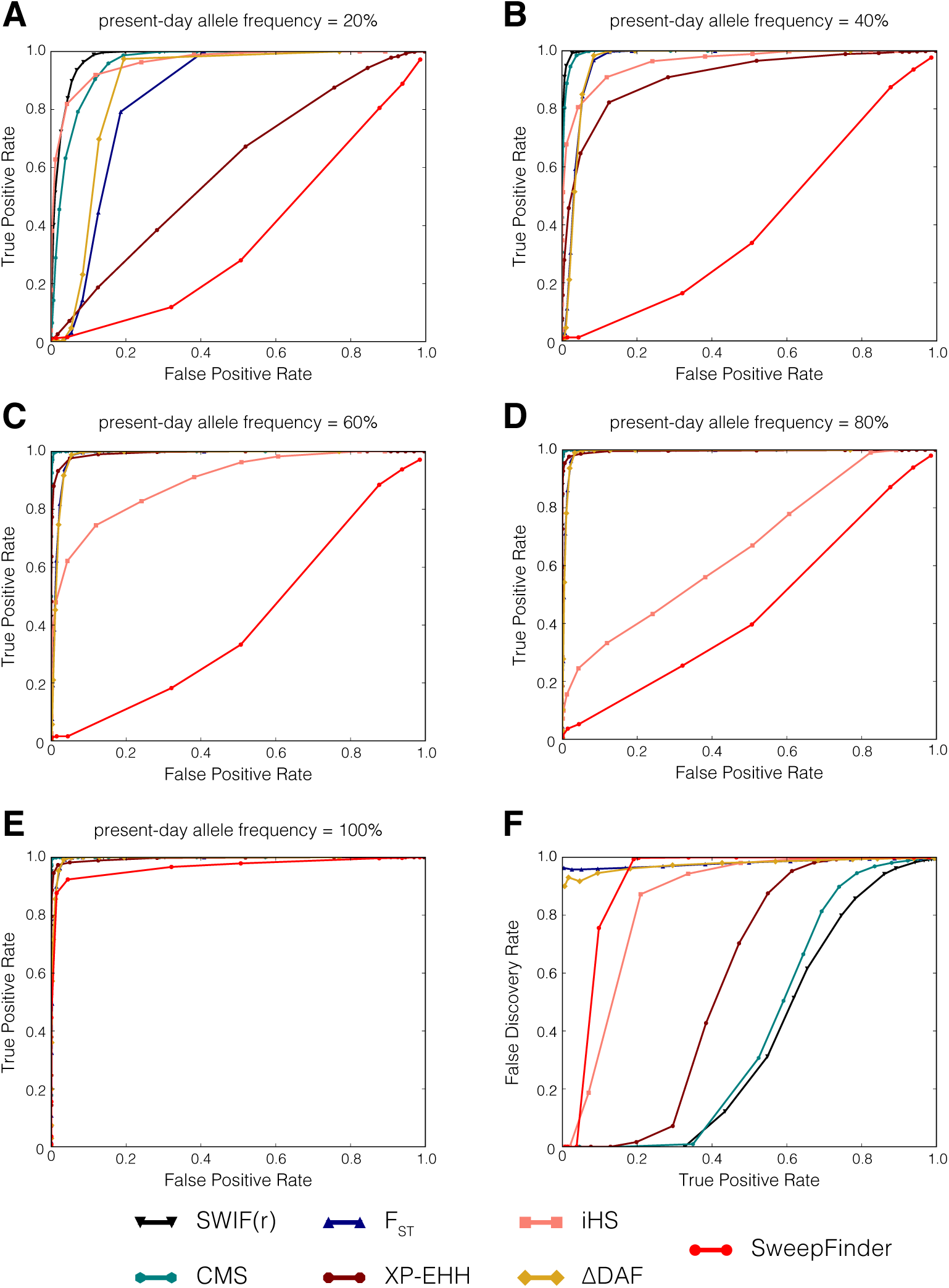
SWIF(r) outperforms sweep-detection method SweepFinder. SweepFinder^3^ is a composite-likelihood method for detecting selective sweeps that is designed to be robust to recombination rate and demography. SweepFinder uses a theoretical model of changes in the allele frequency spectrum after a sweep that is dependent on the distance at each site to the site of the selective sweep, and is designed to detect completed sweeps (present-day beneficial allele frequency of 100% in the population of interest). **A-E)** As expected given SweepFinder’s underlying model, the ROC curves for detecting incomplete sweeps (present-day beneficial allele frequencies from 20% to 80%) show very poor performance of SweepFinder relative to other methods. SweepFinder performs better for completed/nearly-completed sweeps relative to incomplete sweeps, but still lags behind other methods including SWIF(r). We note that iHS is missing from the plot of completed sweeps (panel E), because iHS cannot be calculated at such sites^4^. **F)** Power-FDR curves aggregated over all sweep parameters, assuming a training set composed of 99.95% neutral variants and 0.05% adaptive variants.

**Supplementary Figure 7.**
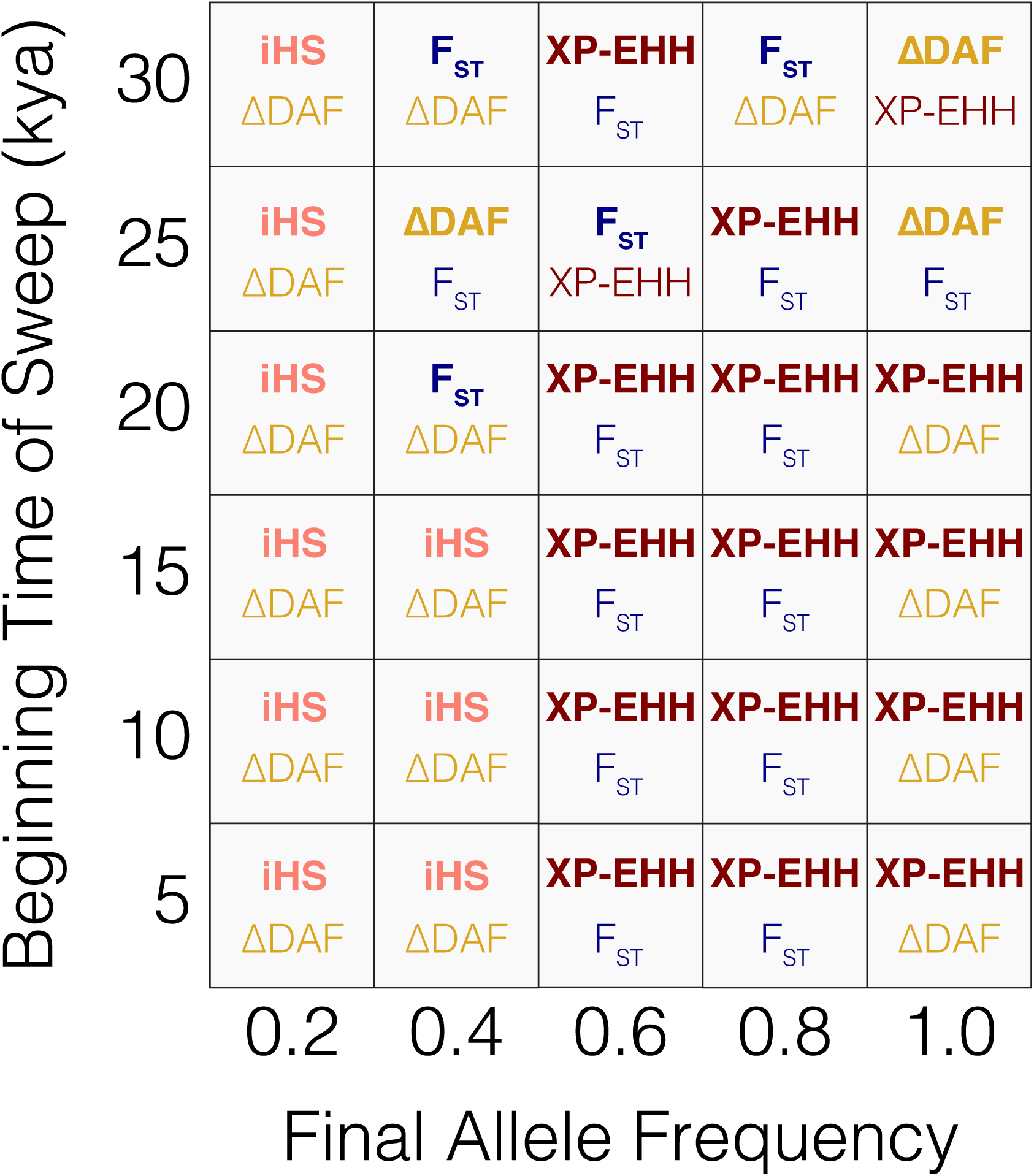
Component selection statistic rankings per sweep model parameters used in this study. Each box contains the top two performing component statistics (listed in rank order) for each pair of sweep start time and present-day frequency of the adaptive allele in the population of interest. Performance is evaluated as the area under the ROC curve. Colors correspond to those in Figure 1.

**Supplementary Figure 8.**
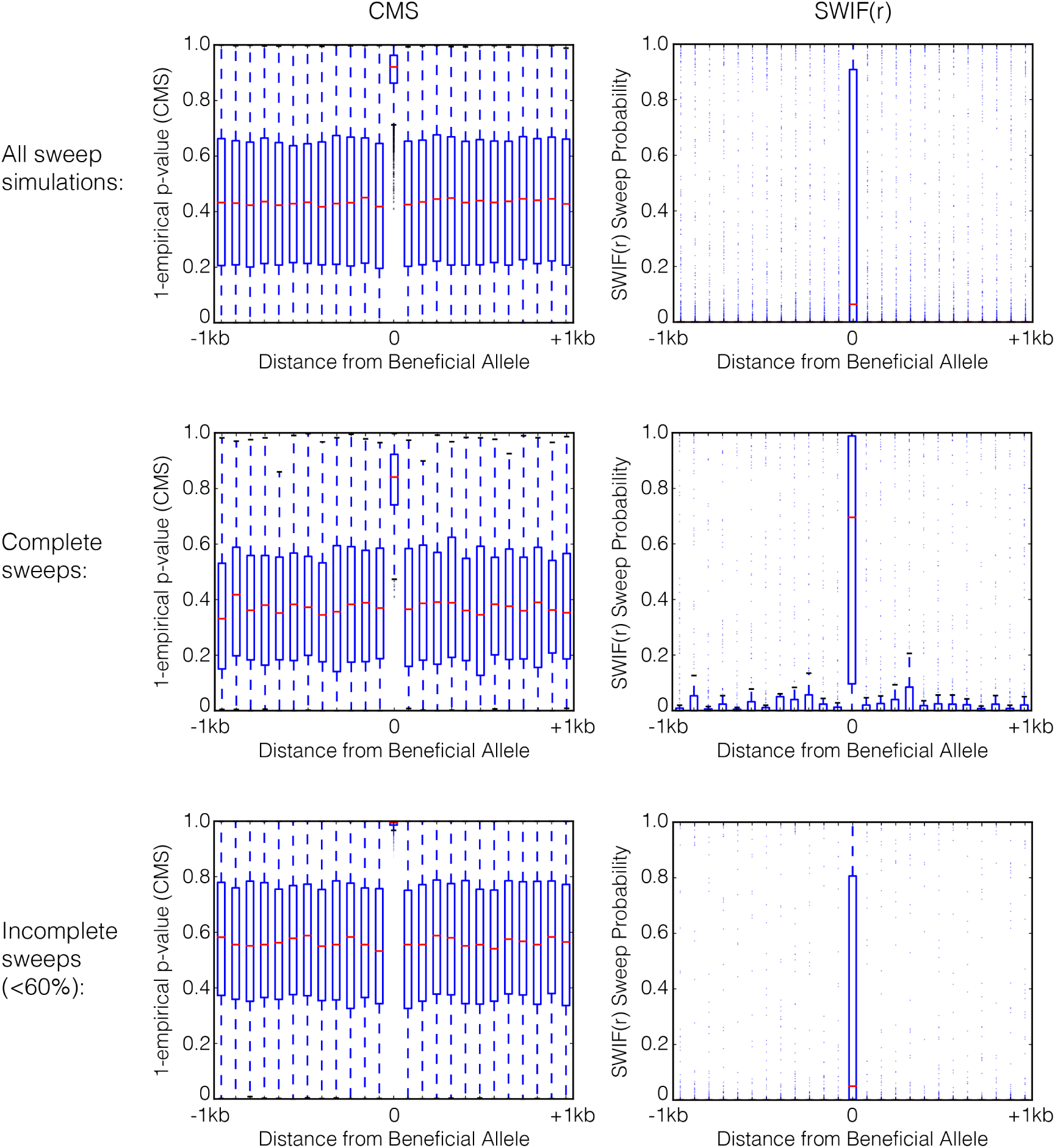
SWIF(r) outperforms CMS in localization of adaptive mutations in simulation. Boxplots show empirical *p*-values calculated with respect to all neutral simulations for CMS (left) and sweep probabilities returned by SWIF(r) (right). Center boxplots contain only adaptive sites, and all other boxplots evenly bin genomic coordinates 1kb up- and downstream of the adaptive SNP. Results are shown for **A)** all sweep parameters, **B)** complete sweeps (beneficial allele has reached a frequency of 100% in the population of interest), and **C)** incomplete sweeps where the beneficial allele has reached a maximum frequency of 60%. In each case, the distribution of CMS scores at the beneficial allele differs from the distributions at neighboring sites, but with significant overlap, making it difficult to confidently and quantitatively localize the adaptive site. In contrast, the distributions of SWIF(r) probabilities at neighboring loci are extremely low with little variance, with only outliers overlapping the range of probabilities seen at adaptive sites.

**Supplementary Figure 9.**
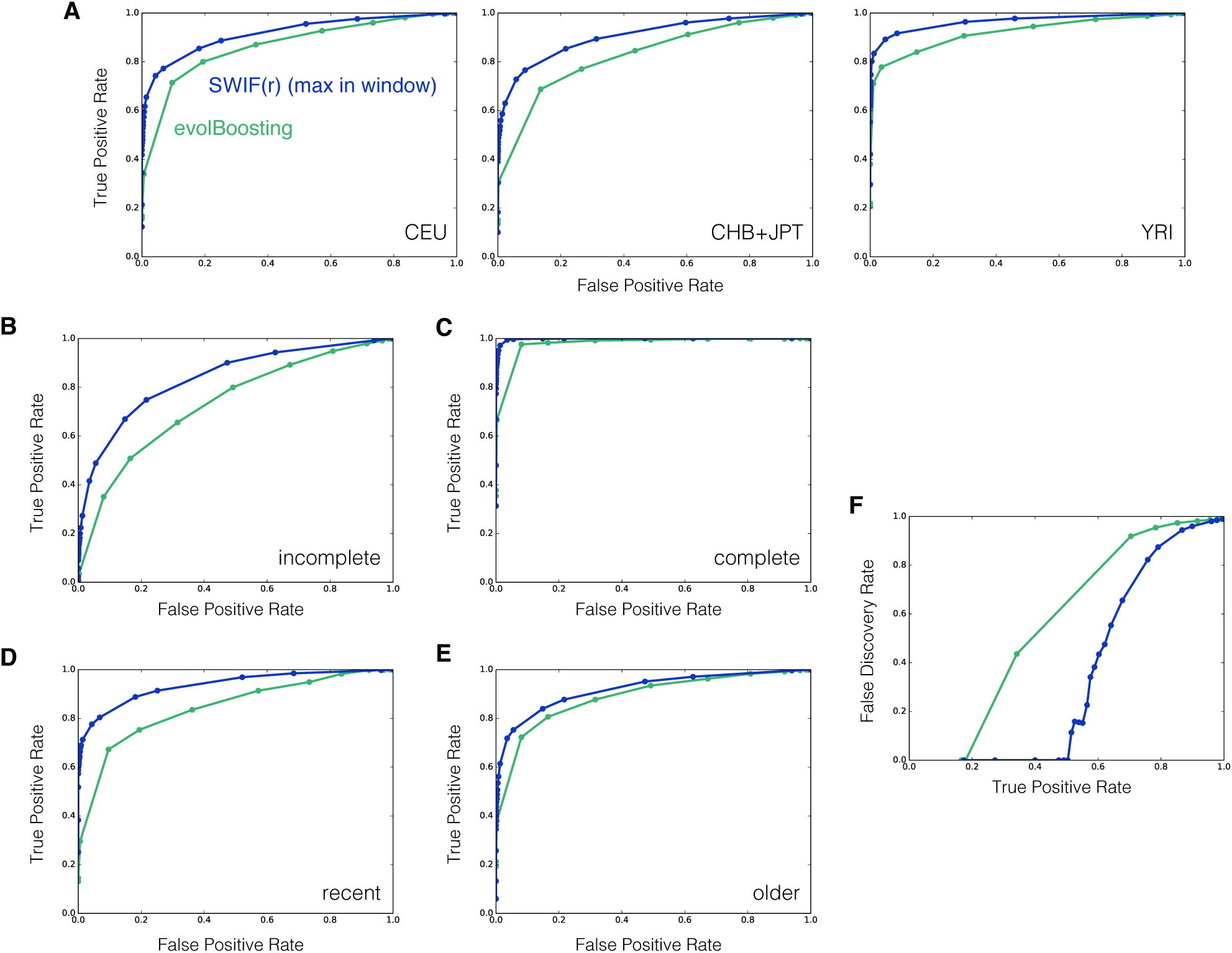
SWIF(r) outperforms evolBoosting^5^. Following Lin *et al.* ^5^, we implemented their software evolBoosting (http://www.picb.ac.cn/evolgen/softwares/) and applied it to simulated data in 40kb windows. For comparison, we calculated “window-based” SWIF(r) scores by taking the maximum site-based SWIF(r) probability in each 40kb window. We tested and trained evolBoosting on the middle 40kb of the 1Mb neutral simulations and sweep simulations described in Online Methods, for three populations modeled after CEU (Europe), CHB and JPT (East Asia), and YRI (West Africa). For a given threshold *α*, the false positive rate is defined as the fraction of neutral windows with a score above *α*, and the true positive rate is the fraction of windows containing a sweep with a score above *α*. The ROC curves below are obtained by varying *α*. **A)** Aggregated over all sweep parameters (beginning time of sweep between 5 and 30kya, present-day allele frequency between 20 and 100%), window-based SWIF(r) outperforms evolBoosting in all three populations. Results in remaining panels are averaged over populations. **B)** ROC curves for incomplete sweeps (present-day beneficial allele frequencies of 20-40%). **C)** ROC curves for complete and near-complete sweeps (present-day beneficial allele frequencies of 80-100% in the population of interest). **D)** ROC curves for sweeps beginning 5-10kya. **E)** ROC curves for sweeps beginning 25-30kya. The performance of the two methods is closest for this parameter set representing older sweeps, likely reflecting the decreased power of haplotype-based statistics, which SWIF(r) relies on, relative to SFS-based statistics, which evolBoosting relies on. **F)** Power-FDR curves aggregated over all sweep parameters, assuming 1% of windows contain a sweep.

**Supplementary Figure 10.**
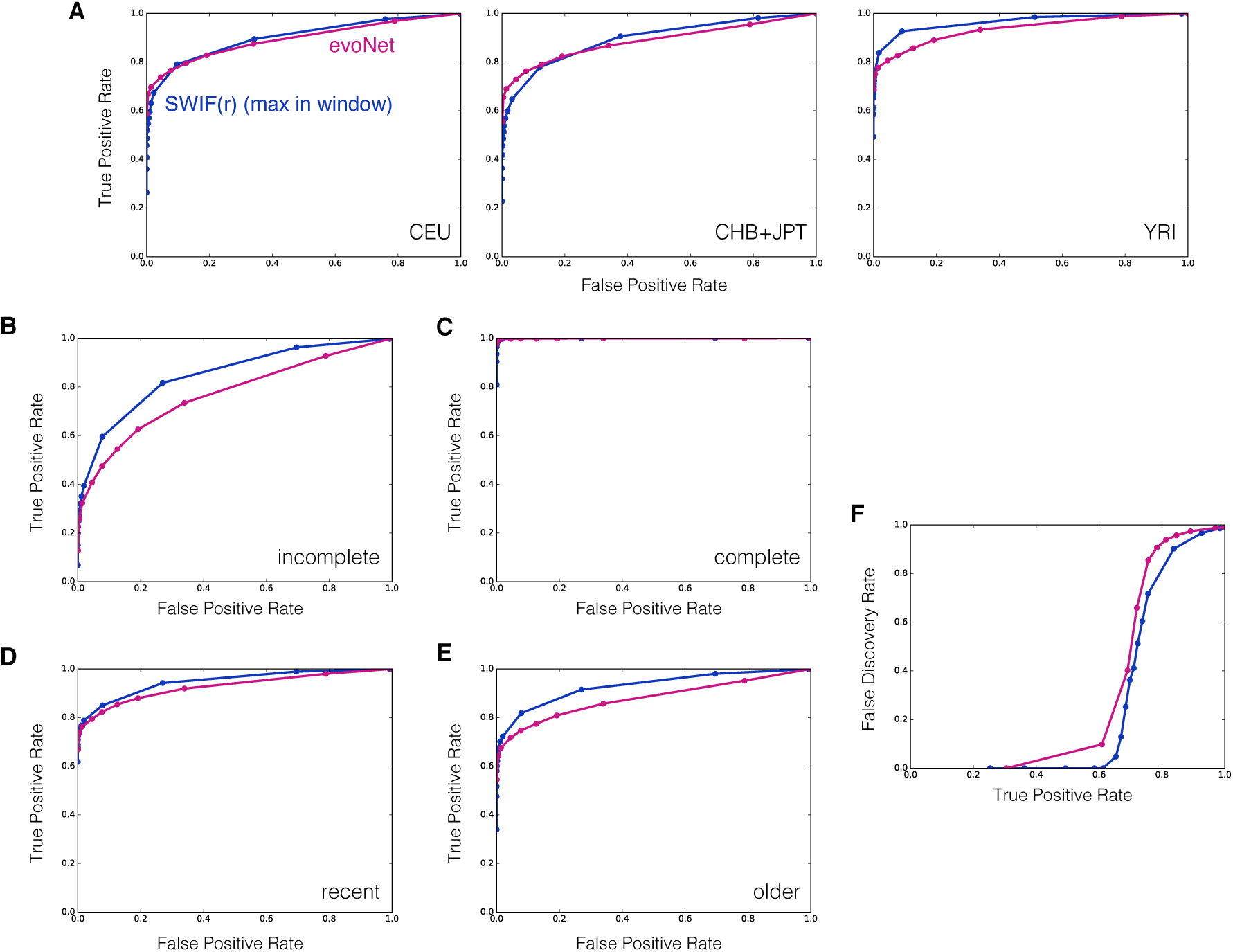
Performance comparison of SWIF(r) against evoNet^6^. evoNet^6^ is a deep learning framework for simultaneous demographic and selection inference; however, the software can be implemented for selection inference only. Following Sheehan *et al.* ^6^, we implemented evoNet in 100kb windows, using the central 100kb of the 1Mb simulations described in Online Methods. For comparison, we calculated “window-based” SWIF(r) scores by taking the maximum site-based SWIF(r) probability in the same 100kb windows. Both evoNet and SWIF(r) are frameworks that can take in any summary statistics, so we trained evoNet using the same component statistics used for SWIF(r). SInce evoNet is a window-based method that makes use of the values of statistics close to the beneficial allele (within 10kb), at mid-range (between 10 and 30kb away), and far away (between 30 and 50kb away), we computed average component statistic values in these three genomic windows for *F_ST_*, XP-EHH, iHS, and ΔDAF. For a given threshold a, the false positive rate is defined as the fraction of neutral windows with a score above a, and the true positive rate is the fraction of windows containing a sweep with a score above a. The ROC curves below are obtained by varying a. We note that these analyses likely downplay the strengths of both methods, as each method has been altered from its original design to enable a direct comparison.**A)** Aggregated over all sweep parameters (beginning time of sweep between 5 and 30kya, present-day allele frequency between 20 and 100%), window-based SWIF(r) performs on a par with evoNet in all three populations. Results in remaining panels are averaged over populations. **B)** ROC curves for incomplete sweeps (present-day beneficial allele frequencies of 20-40%). **C)** ROC curves for complete sweeps (present-day beneficial allele frequencies of 80-100% in the population of interest). **D)** ROC curves for sweeps beginning 5-10kya. **E)** ROC curves for sweeps beginning 25-30kya. **F)** Power-FDR curves aggregated over all sweep parameters, assuming 1% of windows contain sweeps.

**Supplementary Figure 11.**
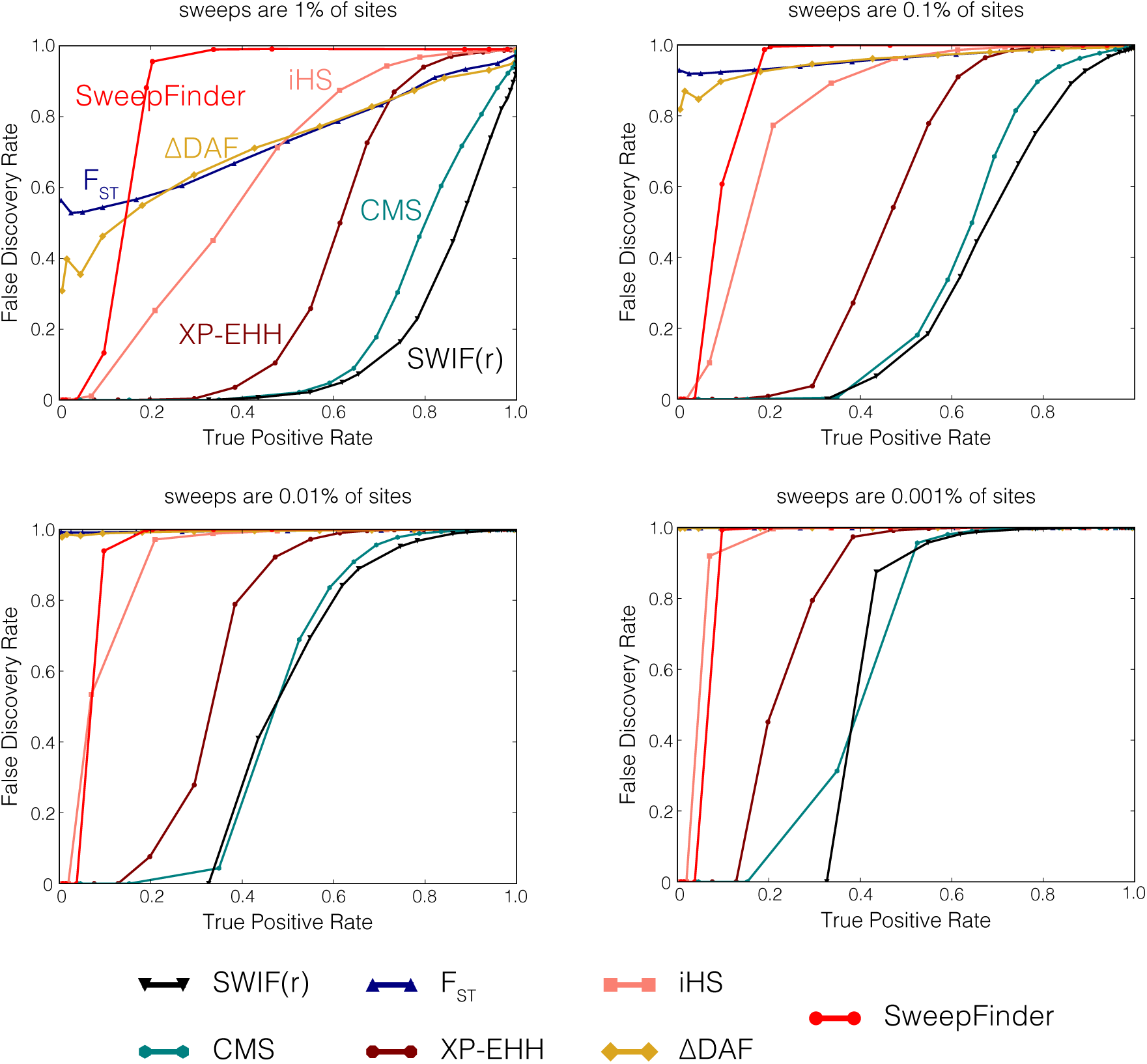
Power versus false discovery rate (FDR) for various training set compositions. Using the same sets of simulations for testing and training that we use to generate the ROC curves in Figure 1, we calculated the tradeoff between true positive rate (power; the fraction of simulated sweep sites correctly classified as a sweep) and the false discovery rate (the fraction of sites classified as a sweep that are actually neutral). We note that the shape of these curves depends heavily on the makeup of the training set, since the false discovery rate depends on the total number of true positives relative to false positives. Below we provide four sets of curves for training sets that are 1%, 0.1%, 0.01%, and 0.001% sweep sites respectively, with the remainder of the training set made up of neutral simulated sites. SWIF(r) performs well relative to CMS and the component statistics across training set compositions, in particular achieving a false discovery rate near zero for a moderate true positive rate of ∼30%. We note, however, that depending on the training set makeup, the false discovery rates for all methods can be quite high; we believe that this illustrates the difficulty of detecting selection reliably at a genome scale. We note that these curves were generated before calibration of posterior probabilities, but that calibrating SWIF(r) would not alter these curves, since the Power-FDR curves themselves are transformation invariant^7^.

**Supplementary Figure 12.**
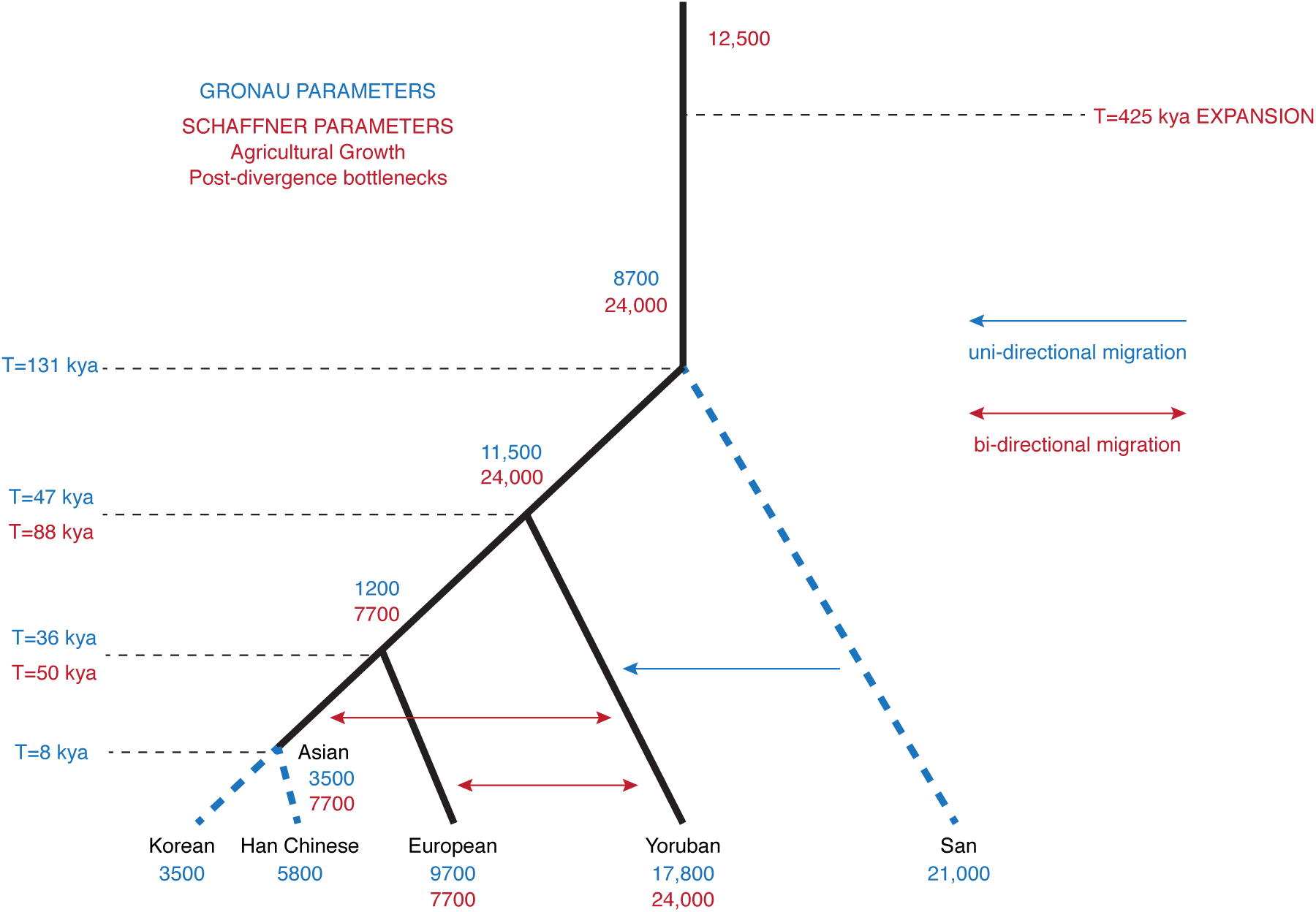
Illustration of differences between Gronau^8^ and Schaffner^9^ demographic models. Black lines indicate lineages inferred by both models, and blue dotted lines are lineages only inferred by Gronau *et al.*^8^. The models differ in many evolutionary parameters, including population sizes, divergence times, and migration rates. In addition, the Schaffner model^9^ includes exponential population growth due to agriculture and post-divergence bottlenecks, while the Gronau model^8^ assumes constant population size.

**Supplementary Figure 13.**
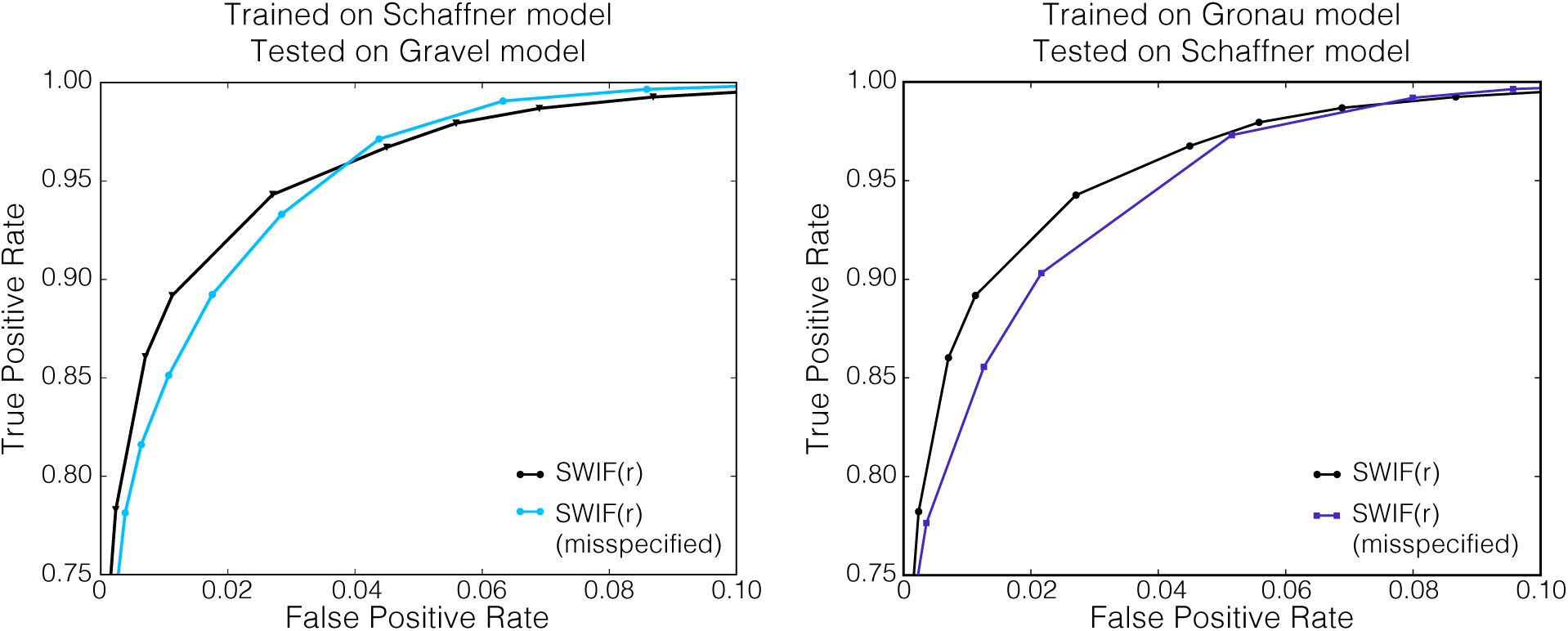
SWIF(r) is robust to demographic misspecification. We conducted two tests of robustness to demographic misspecification. On the left, we trained SWIF(r) on simulations from the Schaffner *et al.*^9^ demographic model, and tested it on simulations from the Gravel *et al.*^10^, which incorporates recent exponential population expansion. On the right, we trained SWIF(r) on simulations from the Gronau model^8^, (including ascertainment modeling), and tested it on simulations from the Schaffner model (see Supplementary Figure 12). In each case, the black curve represents the performance of SWIF(r) when trained and tested on simulations from the same model, and the blue curve represents the performance when misspecified. These results show that misspecification of the demographic model does not have a large effect on the overall performance of SWIF(r). Note that false positive rate ranges from 0% to 10% here.

**Supplementary Figure 14.**
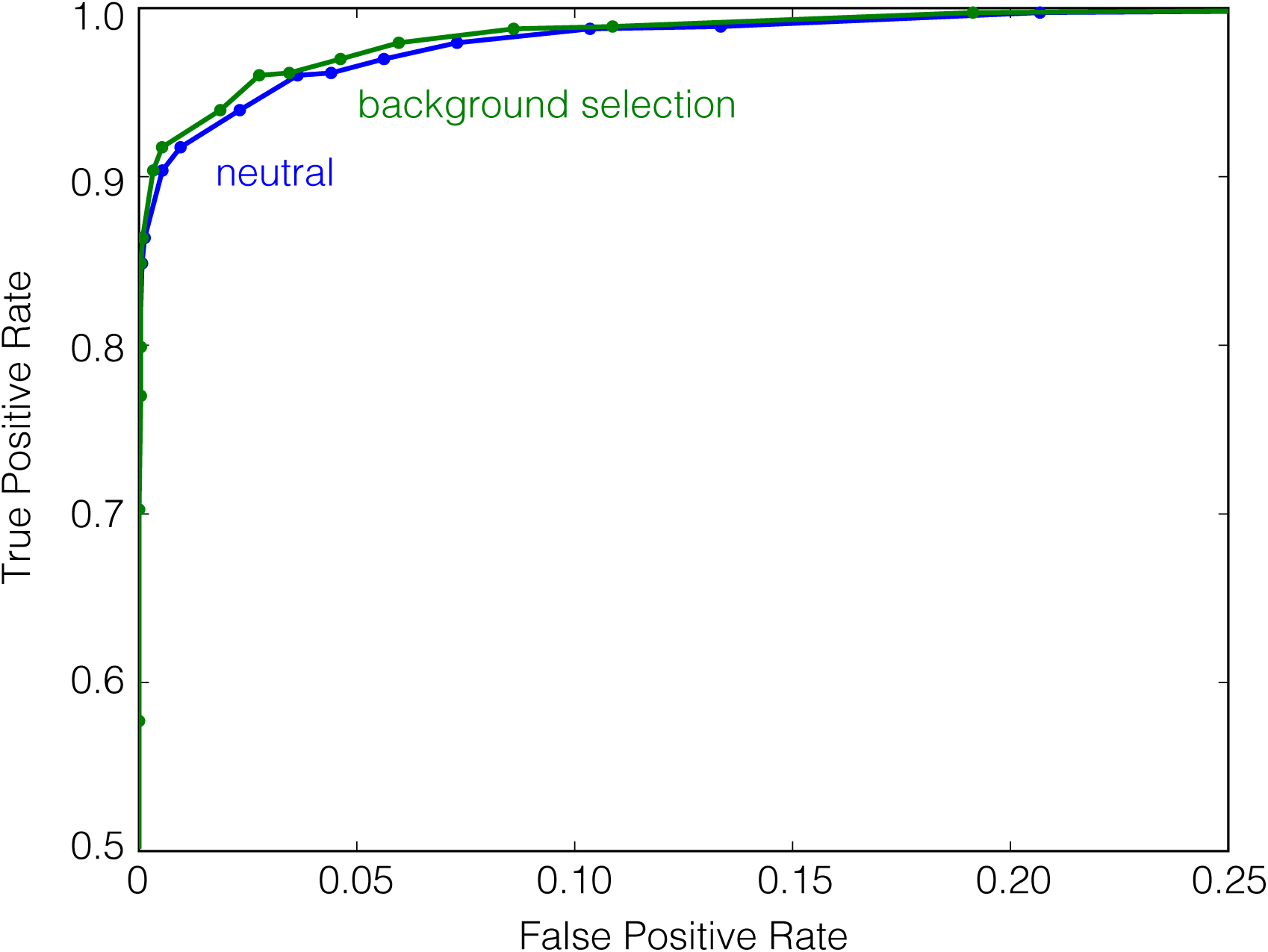
SWIF(r) is robust to background selection in simulation. To test the ability of SWIF(r) to differentiate between selective sweeps and background selection, we generated simulated data for neutral, sweep, and background selection regions using forward simulator `slim`^11^. We followed Messer *et al.*^12^ for simulating background selection: briefly, we simulated genes with 8 exons of 150bp each, separated by introns of 1.5kb each, and surrounded by 550bp and 250bp 5′ and 3′ UTRs, respectively. We assumed that 75% of sites in exons and UTRs were functional (subject to purifying selection), that mutations were codominant, and that fitness effects at different sites were additive. 40% of functional sites were modeled as strongly deleterious (*s* = −0.1), and the rest were weakly deleterious with selection strengths ranging between −0.01 and −0.0001 (see Messer *et al.*^12^). For sweep simulations, selection coefficients were drawn from an exponential distribution with mean 0.03, and sweeps began 10,000 years ago. For all simulations, we modeled two populations with *N_e_* = 5000, with a population split 40,000 years ago, and we used a mutation rate of 2.5 × 10^−8^ and recombination rate of 10^−8^ (also following Messer *et al.*^12^). We trained SWIF(r) using neutral and sweep simulations, and then tested the performance of the resulting classifier in distinguishing neutral variants from sweep variants (blue), and distinguishing exon/UTR variants from sweep variants (green). True positive rate is the fraction of simulated sweep variants that are correctly classified, and false positive rate is the fraction of neutral (blue) or exon/UTR (green) variants that are incorrectly classified as adaptive. These results indicate that SWIF(r) is completely robust to background selection, which is a function of the component statistics it uses; Enard et al.^13^ have shown that iHS and XP-EHH are robust to background selection, and since deleterious alleles are unlikely to rise to high frequency (indeed, are less likely to do so than neutral variants), we would expect that *F_ST_* and ΔDAF would also be robust.

**Supplementary Figure 15.**
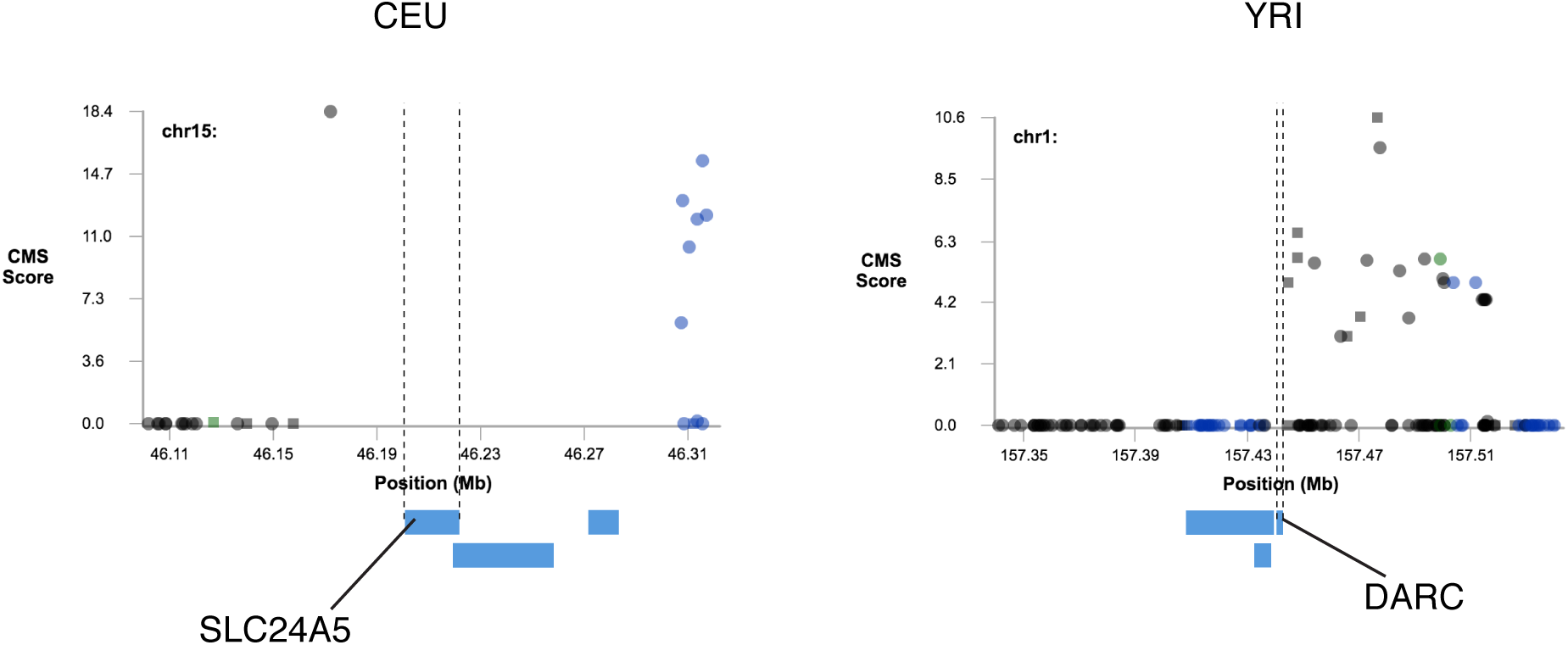
CMS scores on known targets of selective sweeps. Plots made with CMSviewer (https://pubs.broadinstitute.org/mpg/cmsviewer/ use date 04/26/2016) show CMS scores^14^ around *SLC24A5* in CEU and *DARC* in YRI. In each case, SNPs within the gene (region shown between dotted lines) are not assigned CMS scores, possibly due to undefined component statistics. Indeed, in our scan for selection using SWIF(r), iHS and ΔiHH are undefined at the causal mutation in *DARC* and throughout *SLC24A5.* We were unable to obtain plots for *OCA2* and *EDAR* from CMSviewer on subsequent use dates.

**Supplementary Figure 16.**
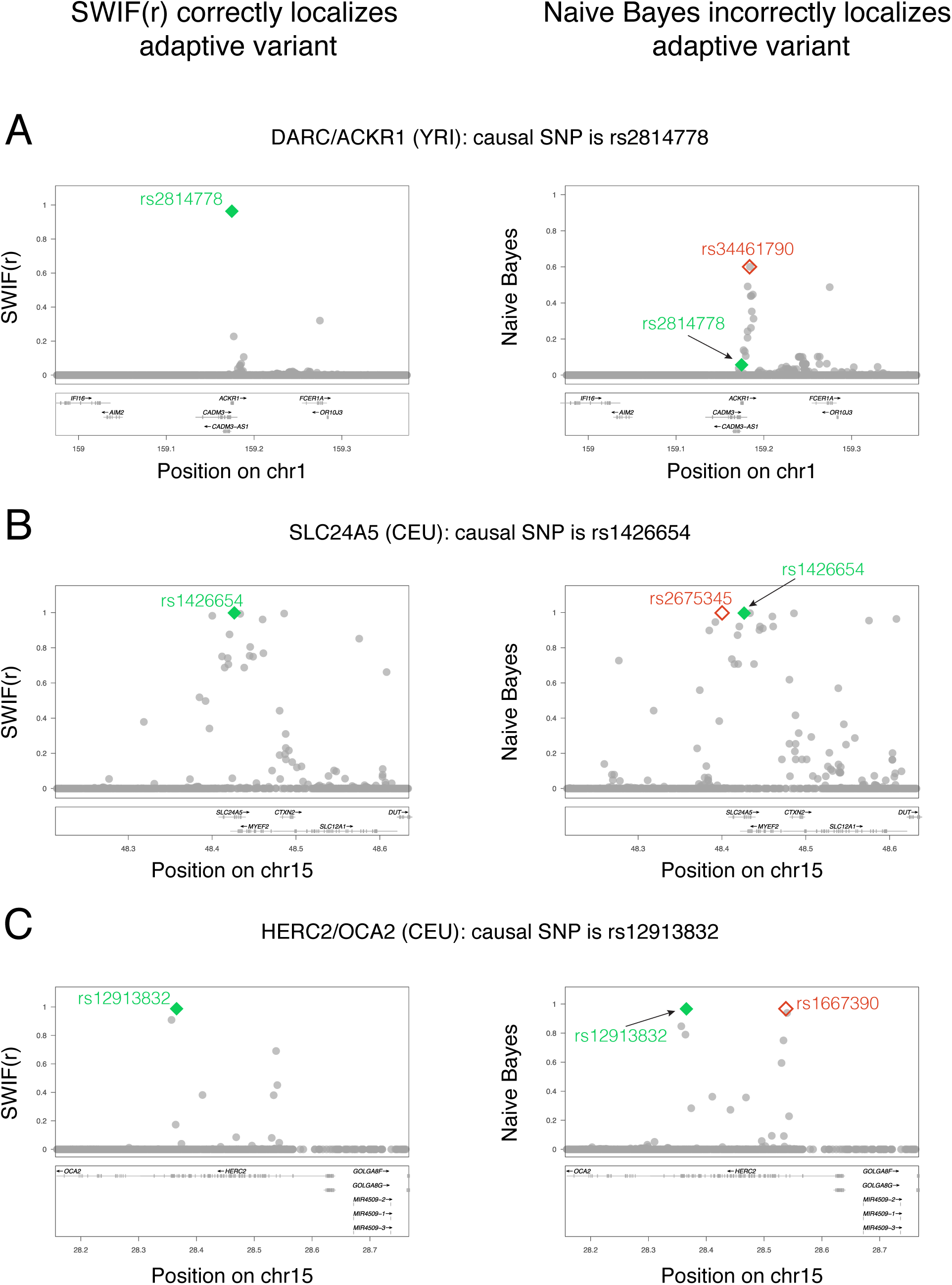
Joint distributions are necessary for localization of previously validated adaptive SNPs. We note that if we do not learn joint distributions and only univariate distributions, then the AODE reduces down to a Naive Bayes classifier, which is very similar to CMS^15^ except that the Naive Bayes classifier returns a probability. Here we compare SWIF(r) to a Naive Bayes classifier in order to illustrate what is gained by learning joint distributions. Posterior probabilities based on SWIF(r) and a Naive Bayes classifier are shown for three genes with validated adaptive SNPs, **A)** *DARC,* **B)** *SLC24A5,* and **C)** *HERC2.* In each case, SWIF(r) assigns the highest sweep probability in the region to the adaptive SNP (in green filled diamond, left column), while the Naive Bayes classifier does not. This is most dramatic for *DARC,* where the Naive Bayes we implemented classifier assigns the causal SNP a very low probability (in green diamond, right column, denoted by arrow), and a different SNP (in red open diamond) is assigned the highest probability. For *SLC24A5* and *HERC2*, the Naive Bayes classifier assigns high probabilities to the causal SNPs, but cannot distinguish these from other similarly high-scoring SNPs. See Figure 2 caption for citations concerning functional validation of these adaptive SNPs.

**Supplementary Figure 17.**
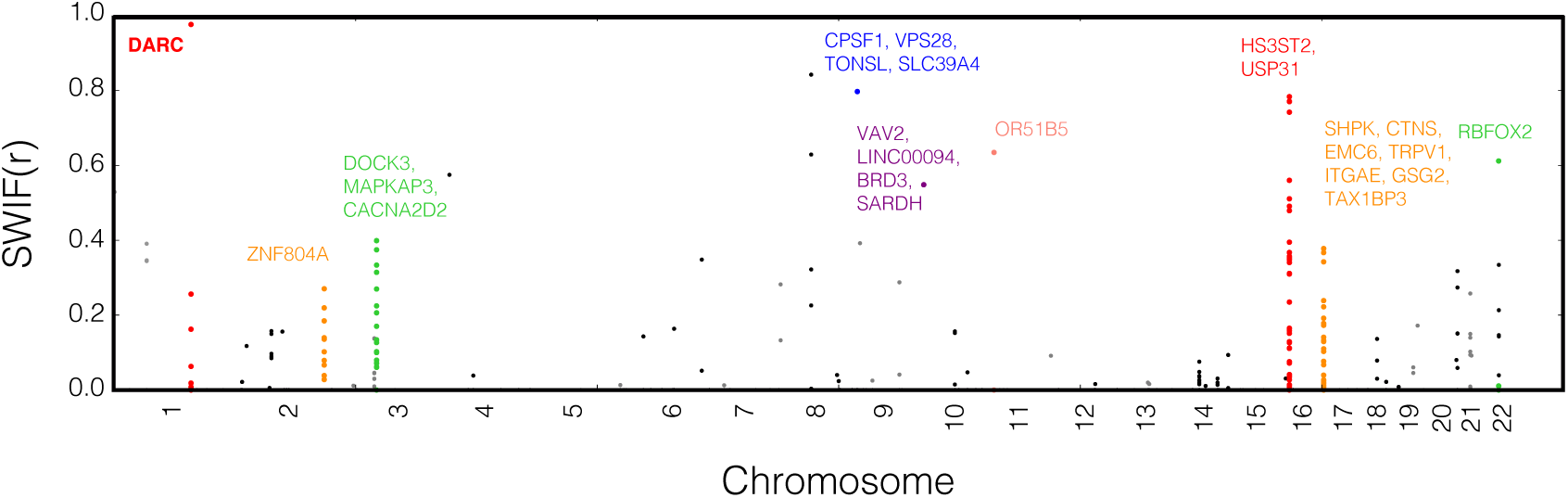
Full selection scan using SWIF(r) in the YRI population (1000 Genomes phase 1^16^). The value plotted for each position along the genome is the calibrated posterior sweep probability calculated by SWIF(r), with per-site prior *π* = 10^−5^ (Supplementary Figure 1). Only SNPs with sweep probability greater than 1% are plotted. Genes containing a single SNP with sweep probability over 50%, or containing multiple SNPs with sweep probability over 10% are annotated. We note that a paralog of *HS3ST2 (HS3ST3A1)* has been linked to malaria resistance^17^.

**Supplementary Figure 18.**
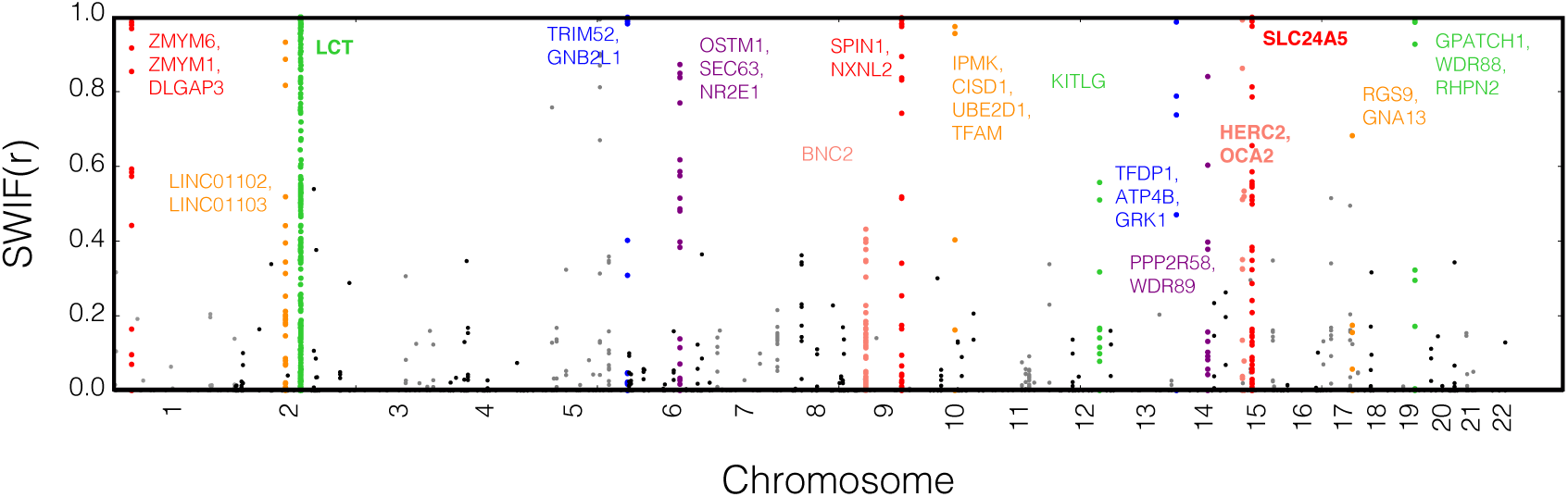
Full selection scan using SWIF(r) in the CEU population (1000 Genomes phase 1^16^). The value plotted for each position along the genome is the calibrated posterior sweep probability calculated by SWIF(r), with per-site prior *π* = 10^−5^ (Supplementary Figure 1). Only SNPs with sweep probability greater than 1% are plotted. Genes containing a single SNP with sweep probability over 50%, or containing a cluster of SNPs with sweep probability over 10% are annotated. In addition to *SLC24A5* and *OCA2,* SWIF(r) also identifies other canonical targets of positive selection in Europeans, including LCT^18^, *BNC2*^19^, and *KITLO*^20^.

**Supplementary Figure 19.**
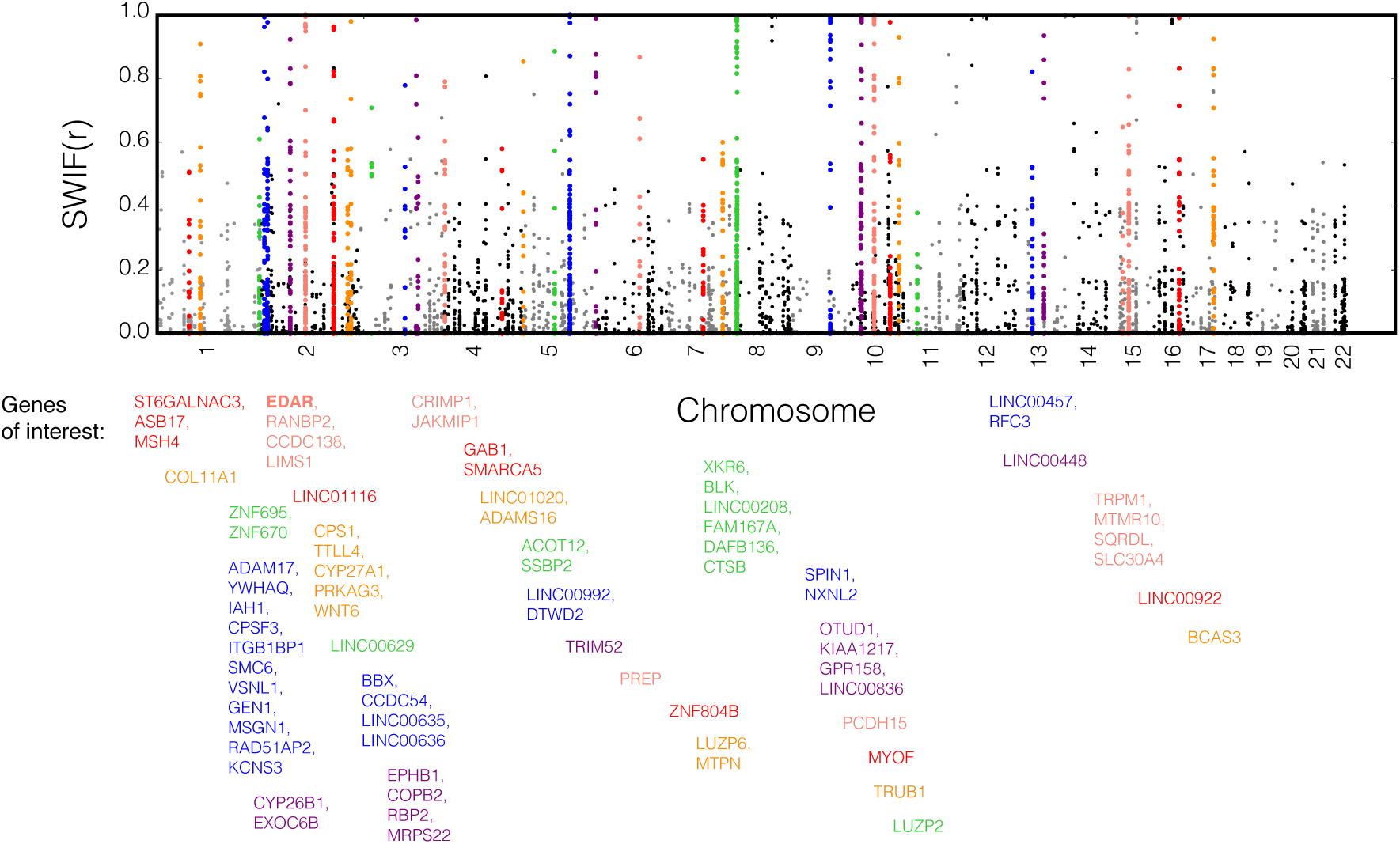
Full selection scan using SWIF(r) in the CHB and JPT populations (1000 Genomes phase 1^16^). The value plotted for each position along the genome is the calibrated posterior sweep probability calculated by SWIF(r), with per-site prior *π* = 10^−5^ (Supplementary Figure 1). Only SNPs with sweep probability greater than 1% are plotted. Genes containing a single SNP with sweep probability over 50%, or containing a cluster of SNPs with sweep probability over 10% are annotated. We suspect that the abundance of selection signals in this population is partly a consequence of longer LD blocks in East Asian populations relative to European and West African populations, and partly a consequence of the fact that the simulation software `cosi`^9^ precludes modeling demographic events during the duration of a sweep, and so very recent population expansions are not modeled in our simulations; Supplementary Figure 20 shows that the distributions of component statistics across populations differs more in observed 1000 Genomes data than in our demographic simulations. We also note that this abundance of signal in East Asian populations has been previously observed^21^.

**Supplementary Figure 20.**
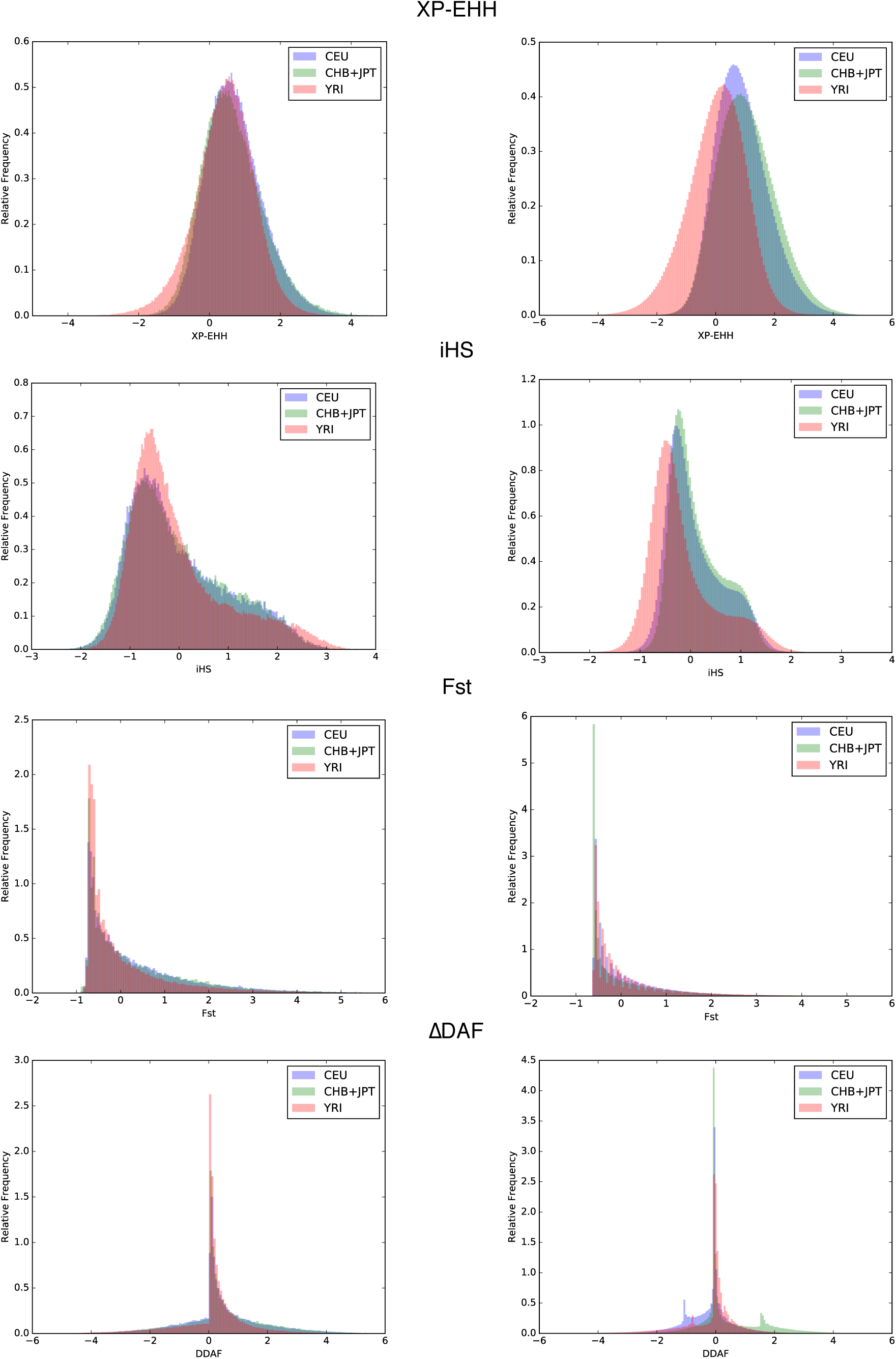
Component statistic distributions for 1000 Genomes data and simulations suggest reasons for discrepancy in number of SWIF(r) targets across populations. The left column shows histograms of component statistic values in simulated data using the demographic model from Schaffner *et al.*^9^, and the right column has the corresponding histograms for the 1000 Genomes data. XP-EHH shows greater bias by population in real data than in simulated data, even after normalizing with population-specific mean and variance based on simulations, with more negative values in the African population than in the other two. iHS shows a similar pattern, despite having been designed to minimize this bias^4^. The distributions of ΔDAF in real data also deviate substantially from simulations; the East Asian population in particular has values that are skewed higher than in the other two populations, possibly as a result of increased genetic drift^22^. These differences may be a consequence of the fact that our simulations did not include very recent population expansions, because of limitations of the simulation software `cosi` regarding the overlap of selective sweeps and demographic events^9^. Most of these differences likely have the effect of leading to an increase in the number of predicted sweep sites in East Asia, and a decrease in West Africa, as we observe in our scan using the 1000 Genomes data, and as has been observed previously^21^. Note that x- and y-axis limits vary for each panel.

**Supplementary Figure 21.**
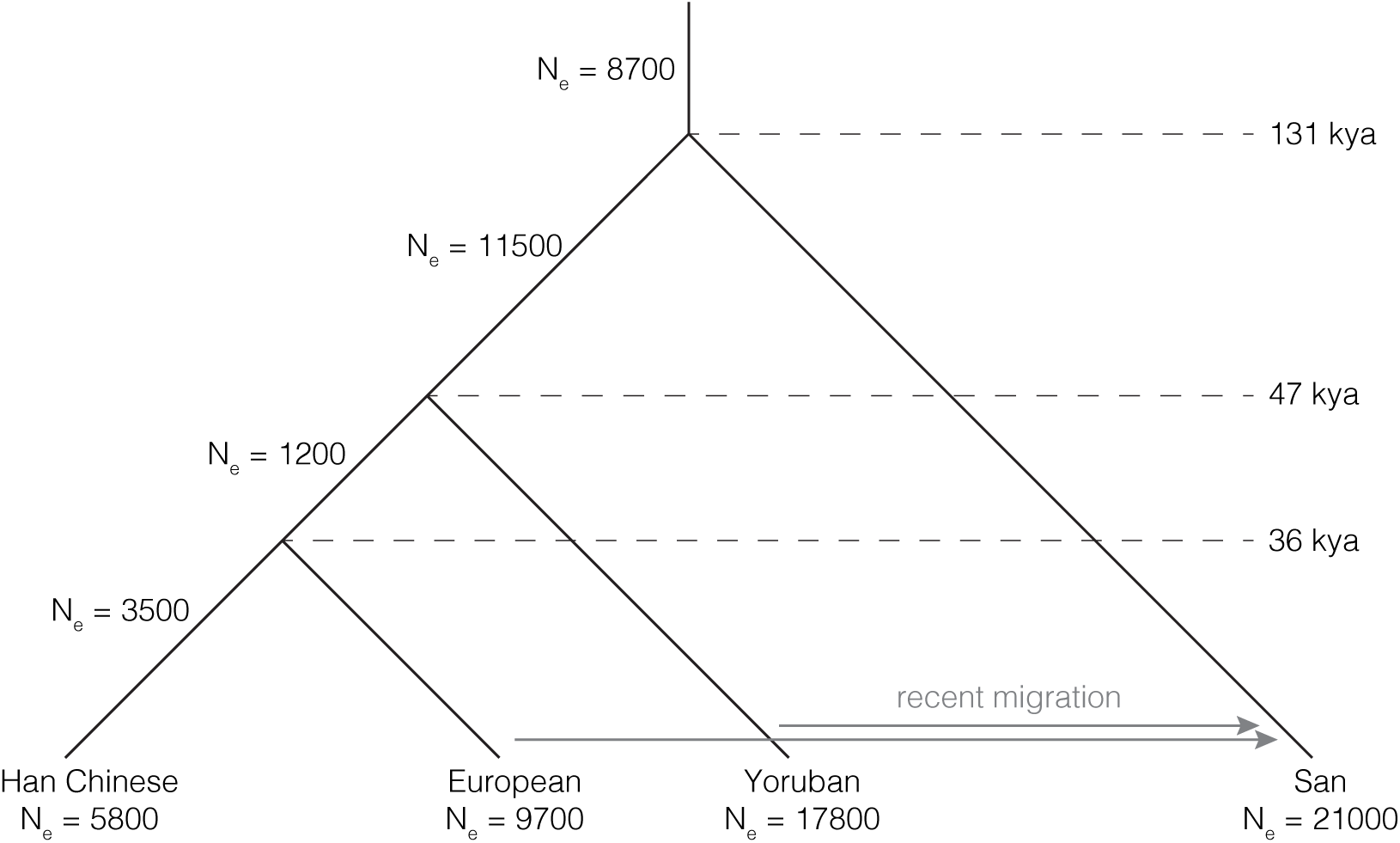
Demographic model used in analyses of the ‡Khomani San. Effective population sizes, coalescent times, and migration rate and times used for simulating haplotype data in analyses including the ‡Khomani San, adapted from Gronau *et al.*^8^. We adapted migration rates from Uren *et al.*^23^: a one-generation pulse from Europe with rate 0.197/chromosome/generation 7 generations ago, and a one-generation pulse from Yoruba with rate 0.227/chromosome/generation 14 generations ago (both shown with arrows in this diagram).

**Supplementary Figure 22.**
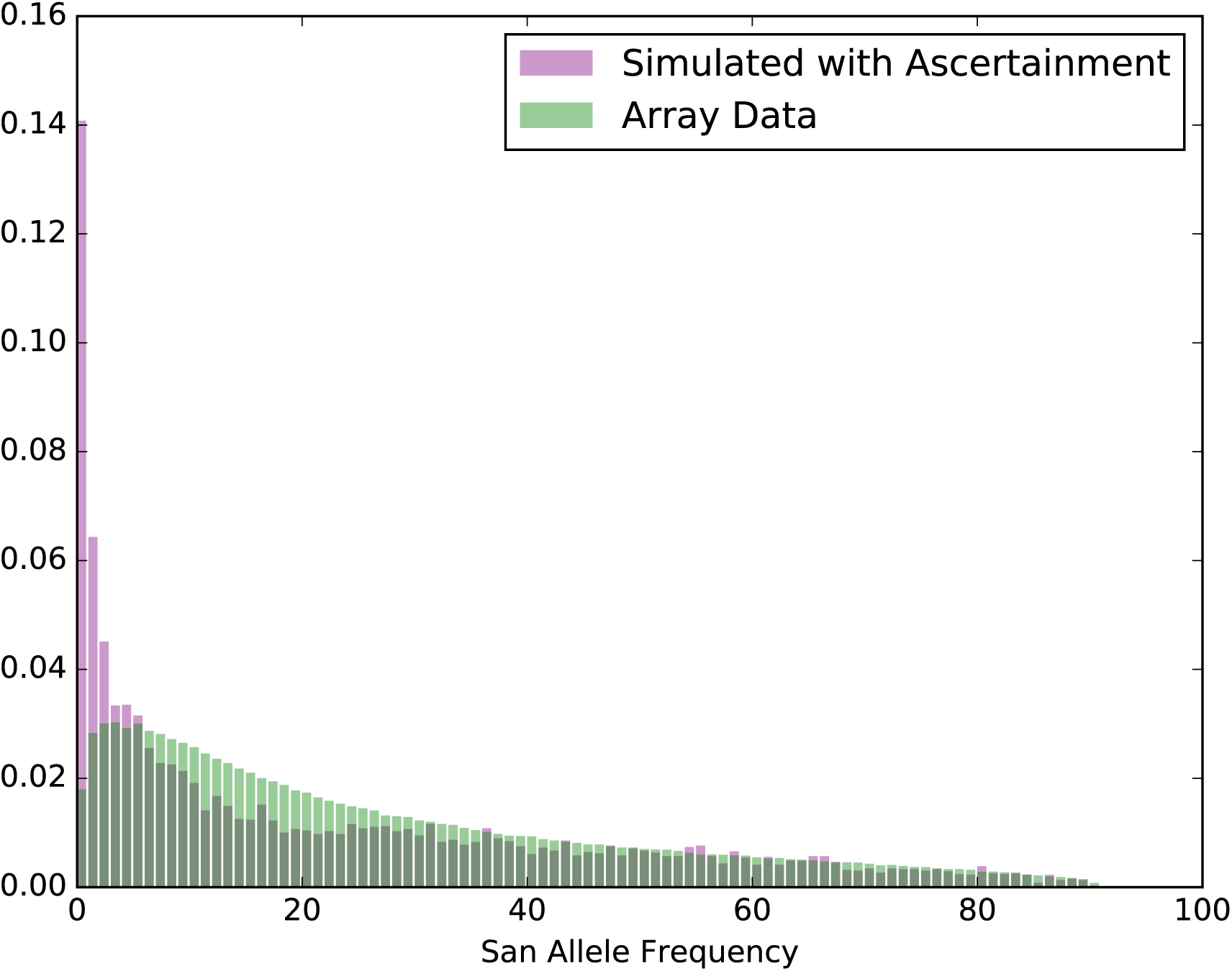
SIte Frequency Spectrum of observed array data and simulated data with ascertainment for ‡Khomani San. After ascertainment on simulated haplotypes, we see fairly good agreement in these site frequency spectra, except for the abundance of very low-frequency derived alleles in simulations. We believe this to be because of standard quality filtering steps taken with array data that are not modeled in the ascertainment process. This difference should not affect the output of SWIF(r) for two reasons: first, we only consider derived mutations in the human lineage as potential sweep targets; and second, SNPs with very low derived allele frequencies are unlikely to register strong signatures of adaptive evolution. Supplementary Figure 23 shows that low-frequency SNPs do not affect the summary statistics used in our implementation of SWIF(r), including haplotype-based statistics like iHS.

**Supplementary Figure 23.**
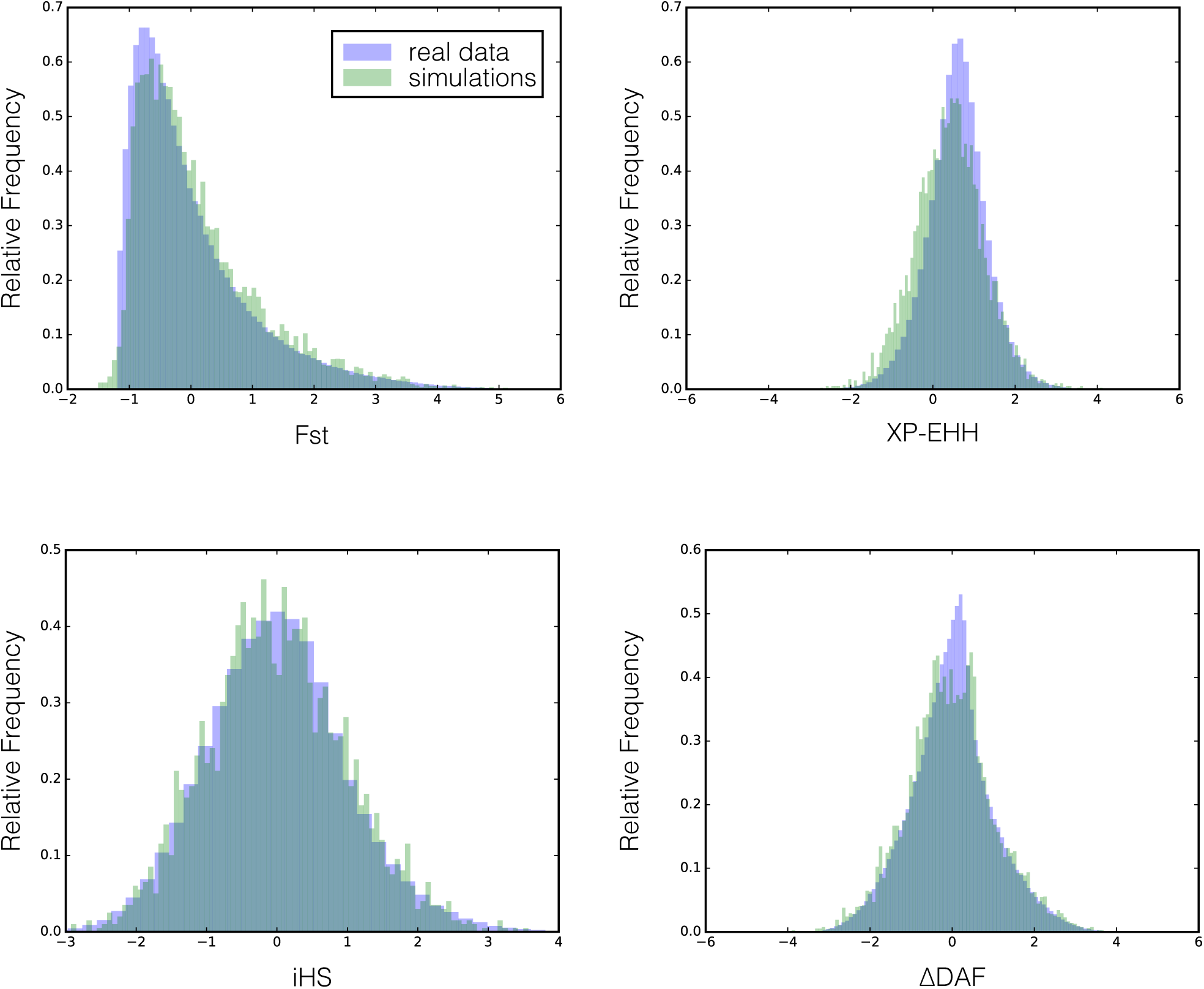
Component statistic distributions for ‡Khomani San simulations and observed data. Despite an excess of very low-frequency derived alleles in the simulated data (Supplementary Figure 22), the distributions of component statistics in simulations and in observed data are quite similar. We are therefore confident that this discrepancy does not inhibit the ability of SWIF(r) to detect selection in this population.

**Supplementary Figure 24.**
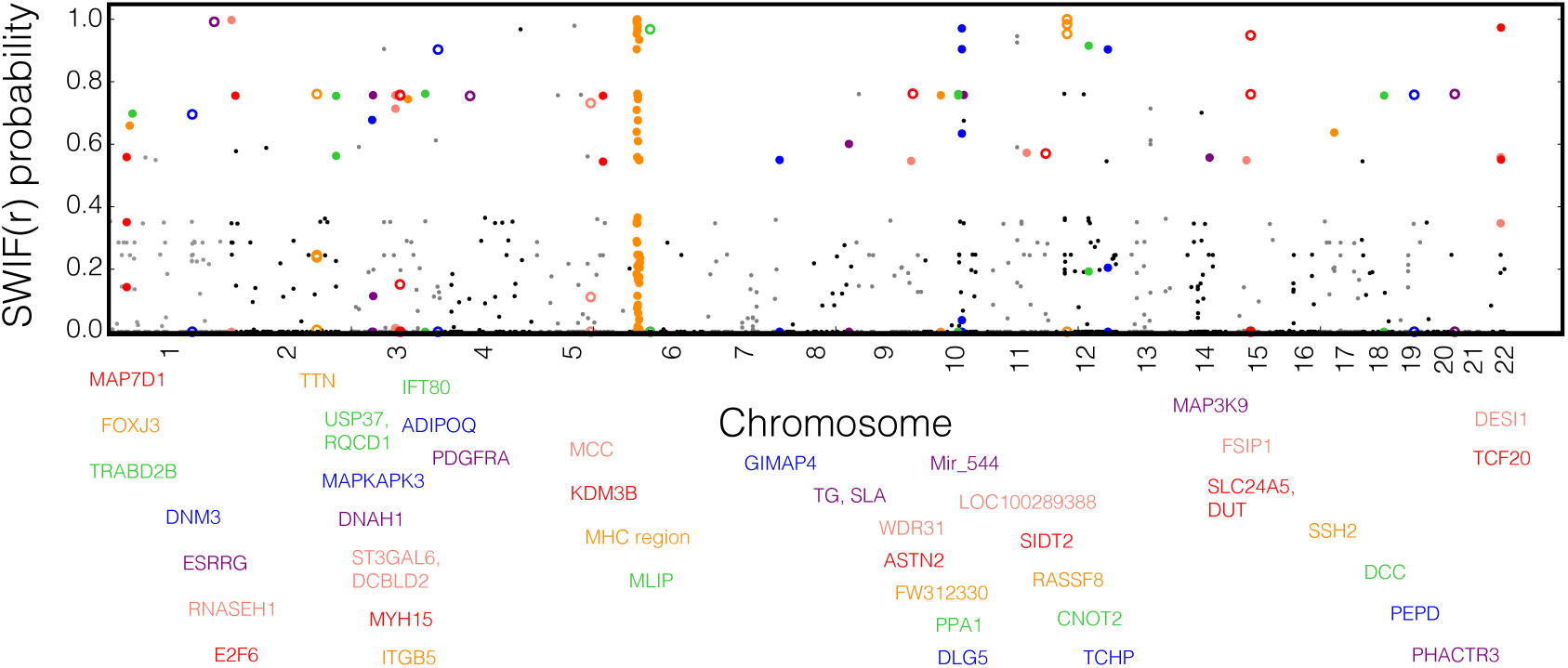
Full selection scan results from analyses of 45 ‡Khomani San samples. The value plotted for each position along the genome is the calibrated posterior sweep probability computed by SWIF(r), with per-site prior *π* = 10^−4^ (Supplementary Figure 2) in order to detect signals of relatively old sweeps given the high long-term *N_e_* of the ‡Khomani San. Only SNPs with posterior sweep probability greater than 1% are plotted. All genes containing at least one SNP with posterior sweep probability over 50% are labeled below the plot, and all variants within those genes are colored to match the gene label. A subset of these results are shown in Figure 3B; genes highlighted in that figure are denoted by open circles.

**Supplementary Figure 25.**
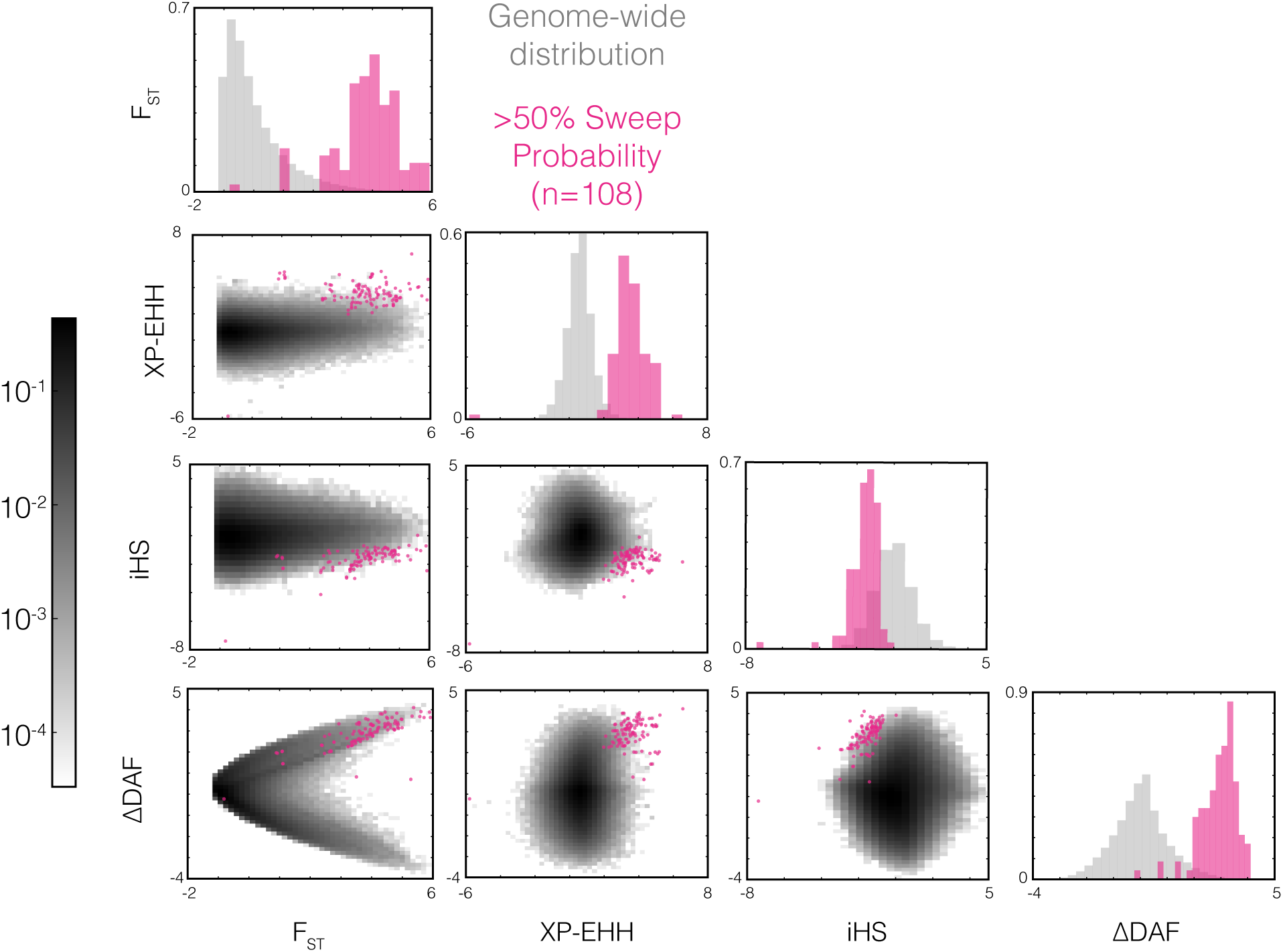
Genome-wide univariate and joint distributions of component statistics used to identify adaptive mutations in the ‡Khomani San, highlighting variants classified by SWIF(r) as adaptive. Distributions along the diagonal show the univariate empirical distributions for each component statistic used when applying SWIF(r) to genomic data from the ‡Khomani San, in gray for the whole genome, and in pink for sites classified as adaptive with a posterior sweep probability over 50%. Both histograms in each plot are normalized, with the genome-wide distributions made up of 628,032 sites for ΔDAF and *F_ST_*, 624,834 for XP-EHH, and 538,928 for iHS. XP-EHH and iHS are calculated at fewer sites because the component statistics are undefined in cases where ΔDAF and *F_ST_* are well-defined. Off-diagonal plots show the joint distributions of each pair of component statistics, again with the empirical genome-wide distribution in gray with a log scale shown in the colorbar, and the classified sweep sites in pink (SWIF(r) posterior probability ≥ 0.5). Any given joint distribution is plotted for only those sites at which both component statistics are defined.

**Supplementary Figure 26.**
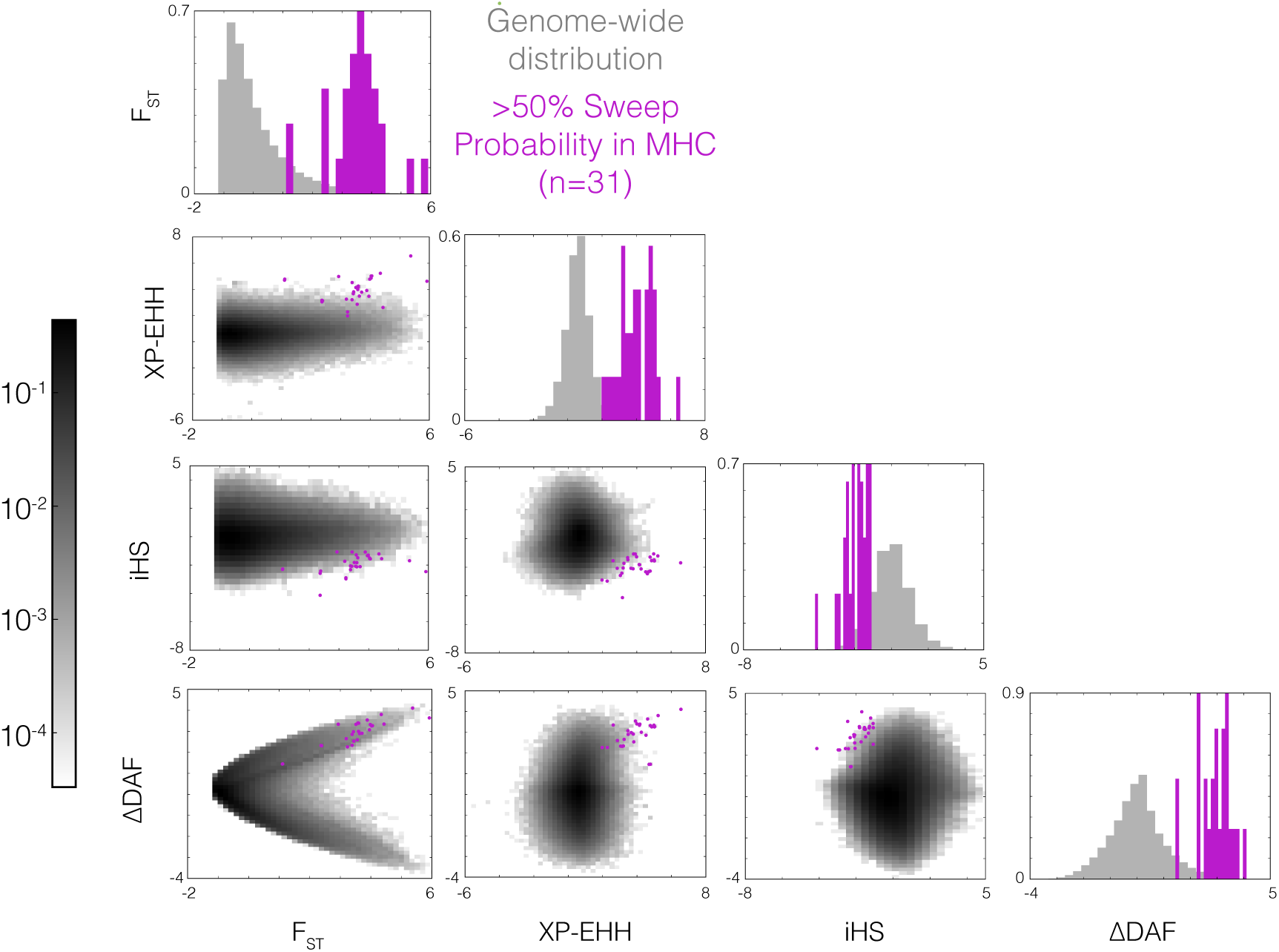
Univariate and joint distributions of component statistics used to identify adaptive mutations in the ‡Khomani San, highlighting SWIF(r) signals in MHC. Distributions along the diagonal show the univariate empirical distributions for each component statistic used when applying SWIF(r) to genomic data from the ‡Khomani San, in gray for the whole genome, and in green for sites within MHC classified as adaptive with a posterior sweep probability over 50%. Off-diagonal plots show the joint distributions of each pair of component statistics, again with the empirical genome-wide distribution in gray with a log scale shown in the colorer, and the classified MHC sites in green. Any given joint distribution is plotted for only those sites at which both component statistics are defined.

**Supplementary Figure 27.**
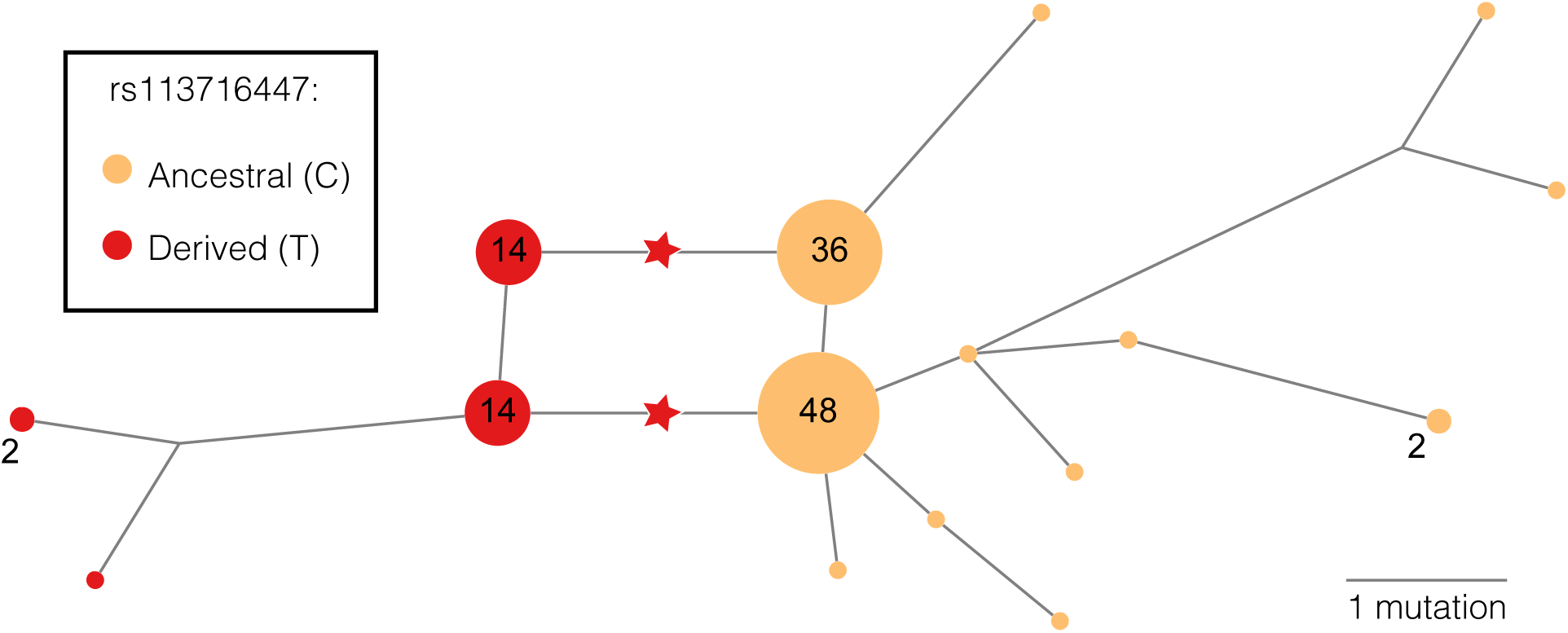
Haplotype network for *ADIPOQ* gene region in the ‡Khomani San dataset. The median-joining haplotype network^24^ is built using the Network software package v4.6.1.1 (fluxus-engineering.com) from combined exome and SNP array data for the ‡Khomani San population. Haplotypes span 17 SNPs over ∼4kb from chr3:186571486-186575536, an interval containing *ADIPOQ*, pruned to avoid recombination hotspots. Nodes represent specific haplotypes and are annotated with the number of times each haplotype appears in the population. Node sizes are scaled relative to the number of individuals represented. Nodes without labels appear once in the population. Edge lengths are proportional to the number of mutations that distinguish the two nodes on either end, and starred edges are those that are defined by the missense mutation at rs113716447. Nodes are colored by whether the haplotype carries the ancestral or derived allele of that gene. Although the available data for this gene region is limited, this network is consistent with selection on the derived allele.

**Supplementary Figure 28.**
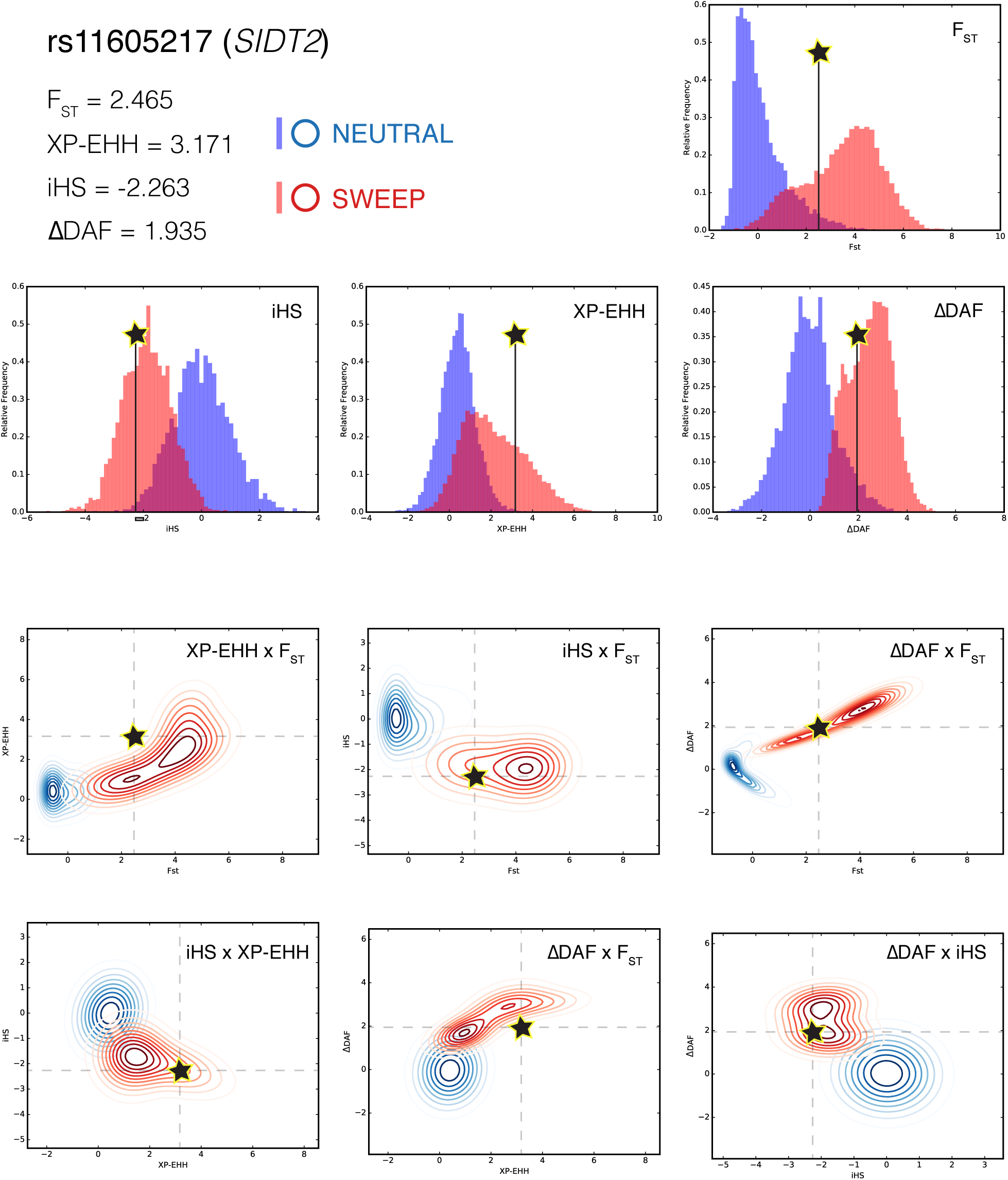
Knowledge of joint distributions allows for identification of *SIDT2* as a target of selection in the ‡Khomani San. The SNP identified by SWIF(r) as the site of an adaptive mutation in *SIDT2*, rs11605217 (posterior sweep probability 61%), had the following values for each summary statistic: *F_ST_* = 2.456, XP-EHH = 3.171, iHS = −2.263, ΔDAF = 1.935. If we do not learn joint distributions and instead only learn univariate distributions (resulting in a Naive Bayes classifier), the posterior probability at this SNP is only 30%. While these statistics individually show moderate evidence for a sweep, their accumulated evidence in a Naive Bayes framework is not enough to overcome a low prior probability, and *F_ST_* and ΔDAF in particular land well within normal limits for their respective neutral distributions (top). However, when we look at the pairwise joint distributions (bottom), in each case, the mutation in *SIDT2* lands within or nearby the sweep distribution, and relatively far from the neutral distribution.

**Supplementary Figure 29.**
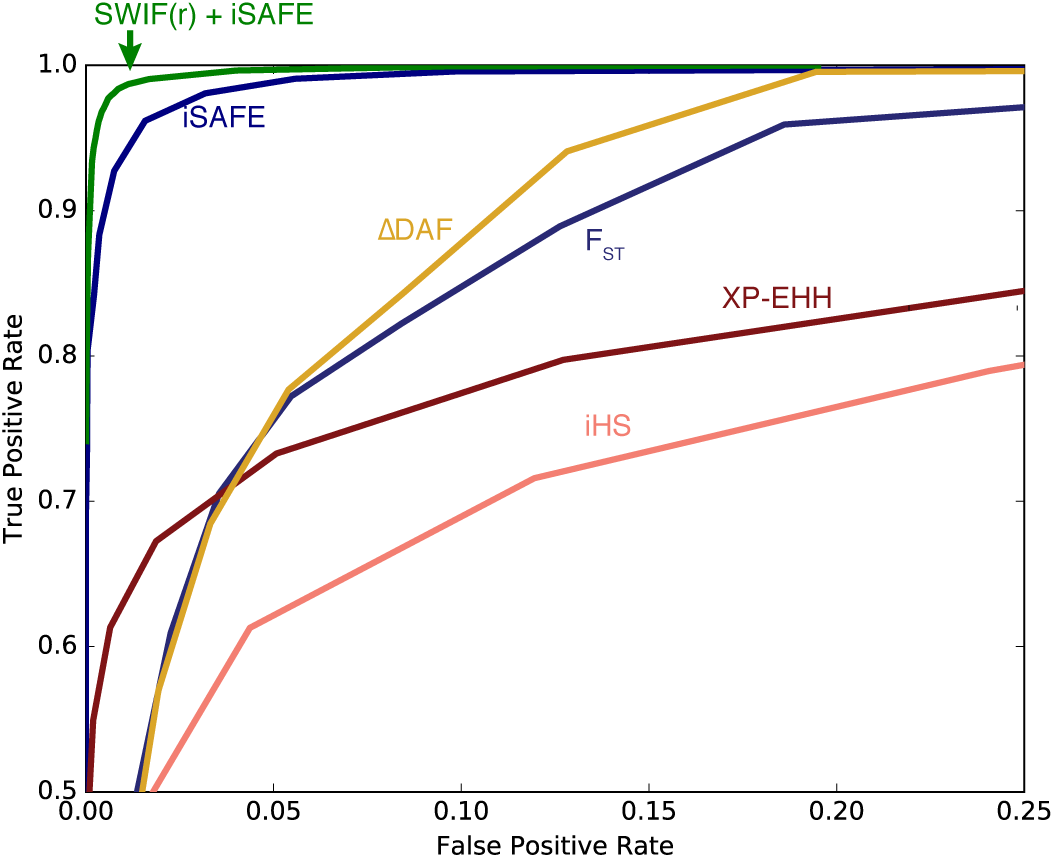
SWIF(r)’s framework can easily incorporate new statistics for increased power. A major feature of SWIF(r) is that it is completely generalizable to any set of component statistics, which means that SWIF(r) can take immediate advantage of more powerful statistics as they are designed. One such statistic, iSAFE ^25^ (preprint available at https://www.biorxiv.org/content/early/2017/10/01/139055), has proven to be much more powerful than current component statistics at localizing the specific site of a beneficial mutation. By incorporating iSAFE as a component statistic, the overall performance of SWIF(r) improves and exceeds that of iSAFE, because SWIF(r) leverages the joint information of iSAFE with other component statistics.

**Supplementary Figure 30.**
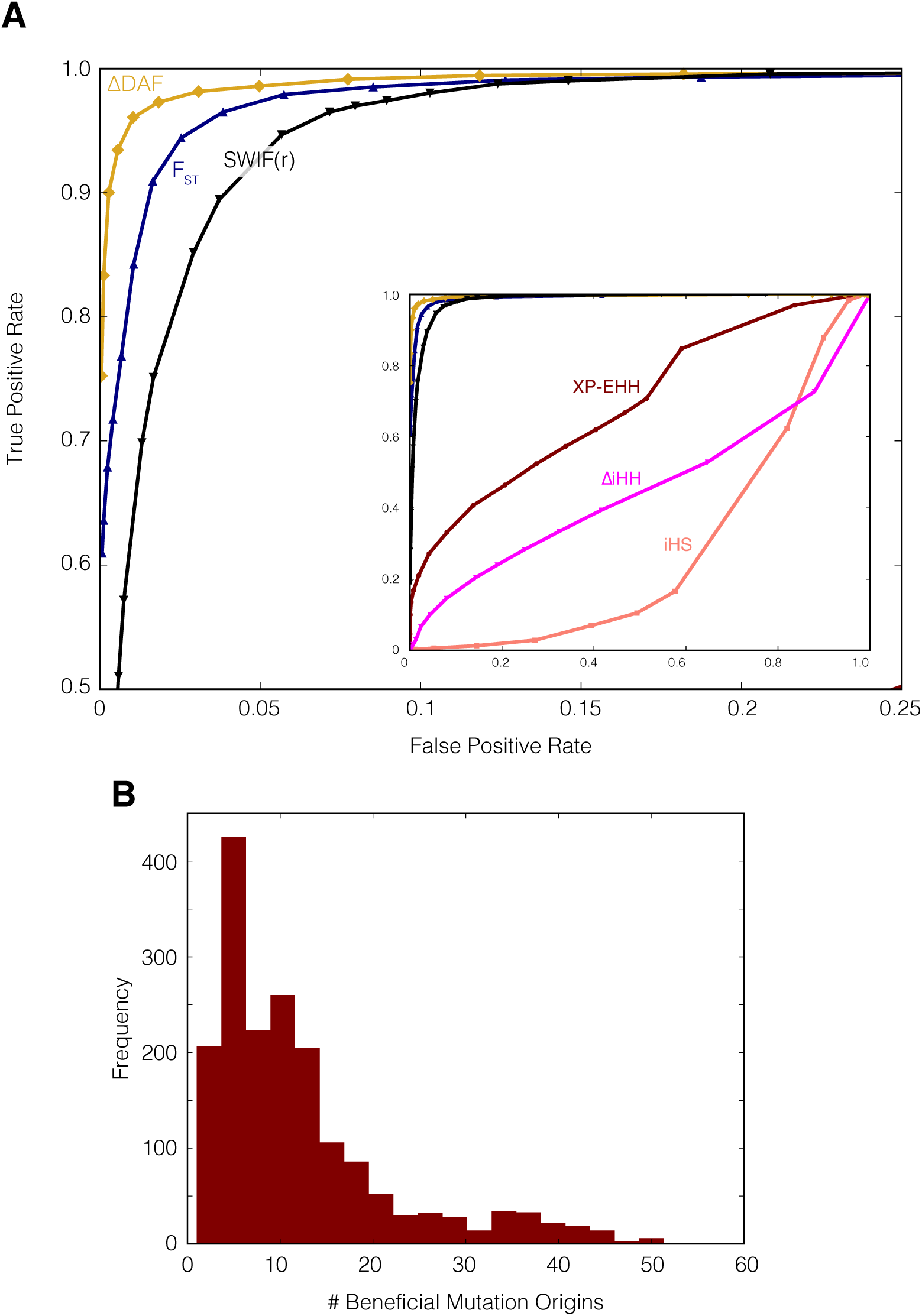
SWIF(r) also detects signatures of soft sweeps from standing variation, despite being trained on hard sweep simulations. Neutral and soft sweeps simulations were generated using `msms`^26^ with parameters designed to match the Schaffner demographic model^9^ (command line code: java -jar msms/msms.jar -N 10000 -ms 360 1 -I 3 120 120 120 -t 600 -r 600 -n 1 2.4 -n 2 0.77 -n 3 0.77 -m 1 2 1.28 -m 2 1 1.28 -m 1 3 0.32 -m 3 1 0.32 -en 0.049975 2 0.00247491582263 -en 0.049975 3 0.000720979721217 -ej 0.05 3 2 -en 0.05 2 0.77 -en 0.087475 1 0.00622496653265 -en 0.087475 2 0.00056286521273 -ej 0.0875 2 1 -en 0.0875 1 2.4 -en 0.425 1 1.25), with the initial frequency of the beneficial allele set to 0.02 for soft sweep simulations. **A)** ROC curves are generated as in Figure 1, with false positive rate being the fraction of neutral variants incorrectly classified as adaptive, and true positive rate being the fraction of adaptive mutations originating as a standing variant (soft sweep site) that are correctly classified as such. As might be expected, population differentiation component statistics perform well at distinguishing between neutrality and soft sweeps, while performance of iHS, XP-EHH, and ΔiHH suffers dramatically at this task relative to the task of distinguishing between neutrality and hard sweeps (inset panel). Nonetheless, SWIF(r) maintains power to detect the genomic signatures of adaptive standing variants. **B)** Although the initial frequency of the beneficial mutations is 2% in these simulations, the vast majority of these simulations result in soft sweeps in the population of interest at the time of sampling, with more than one (and in some cases many) of the original beneficial mutation origins still present at the time of sampling. In approximately 1% of our simulations, we observe a “hardening” of the sweep (i.e. only one mutation origin is present at the time of sampling). Note that we do not include any simulations in which the beneficial mutation is lost.

**Supplementary Figure 31.**
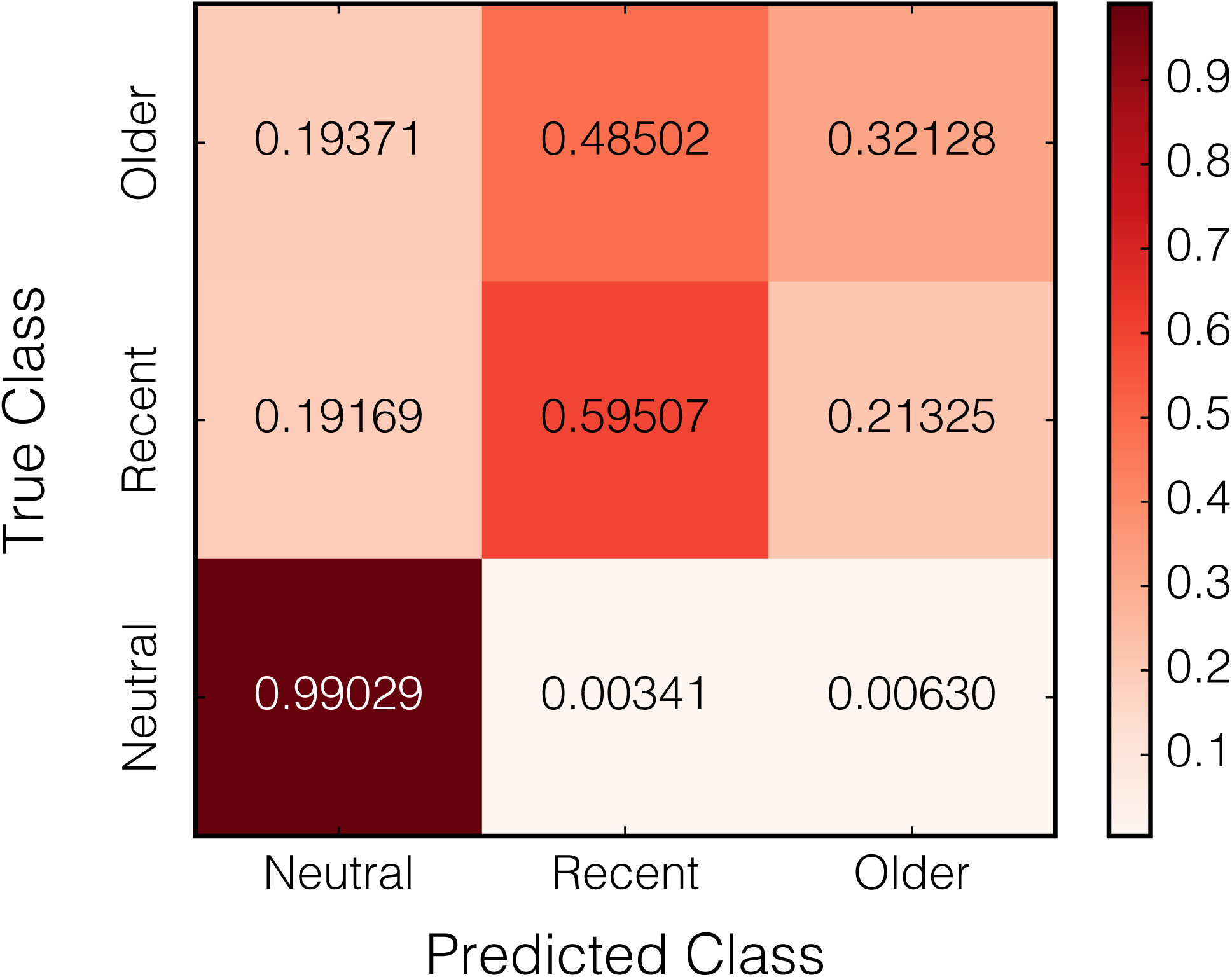
SWIF(r) extends trivially to multi-class classification. In addition to neutral simulations, SWIF(r) was trained on simulations of “recent” sweeps in the ‡Khomani San (5-30kya), and “older” sweeps (30-60kya). Training SWIF(r) on these three classes (“neutral”, “recent”, and “older”) merely requires computing three likelihoods instead of two (see Equation 2), with all three included in the denominator of Equation 3. Posterior probabilities can then be computed for each of the three classes. We classified loci by assigning them to the class with the highest posterior probability. Values in the confusion matrix are the conditional probability of classifying a site in the predicted class, given that the site belongs to the true class. Priors are 0.9 for neutral, and 0.05 for each of the two adaptive classes. In this case, the component statistics do not carry enough power to reliably distinguish between these two sweep timings, likely because of confounding effects of sweep strength and present-day allele frequency. It is plausible that by incorporating other component statistics, SWIF(r) may be able to achieve this timing inference.

**Supplementary Figure 32.**
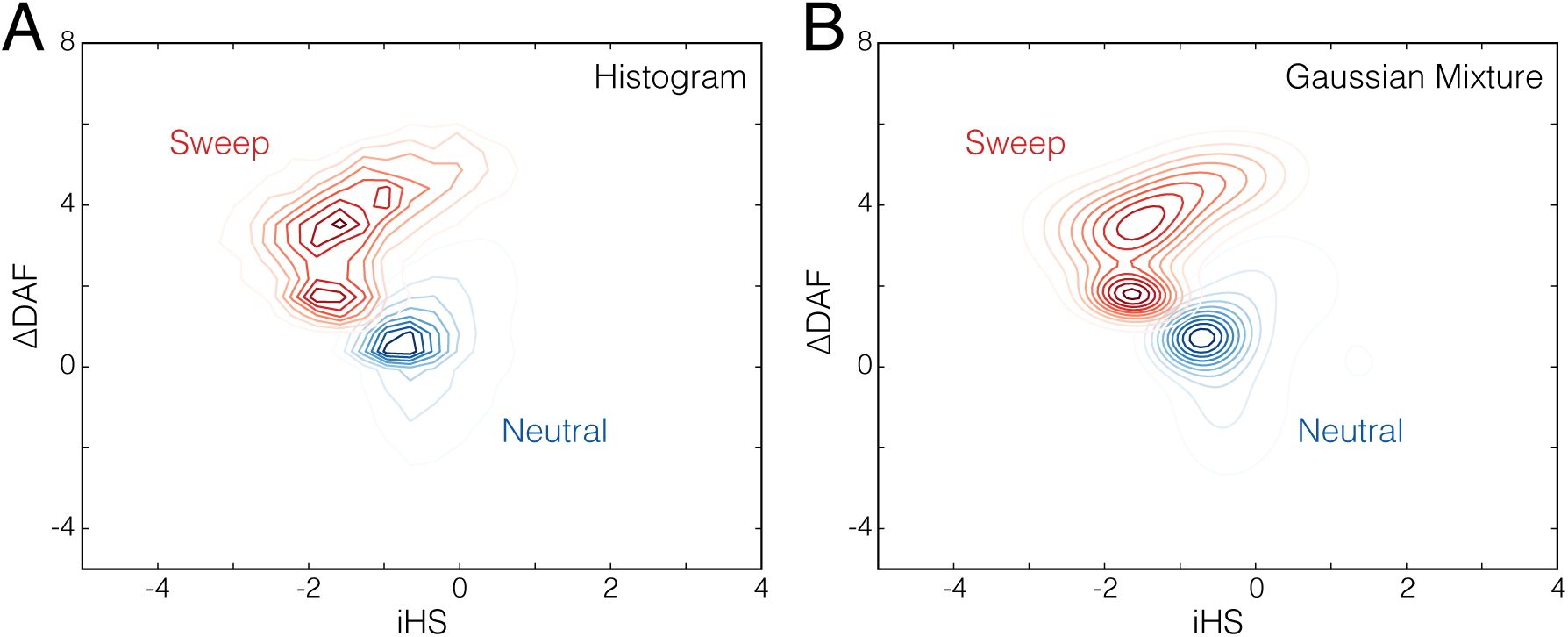
Using Gaussian mixture models to approximate joint likelihoods of selection statistics. The joint distributions of iHS and ΔiHH under neutral and sweep scenarios constructed with **A)** a 30×30 binned histogram approach, and **B)** Gaussian mixture models. The mixture models adequately smooth the distributions while maintaining their general shape and modality, thereby avoiding over-fitting.

**Supplementary Figure 33.**
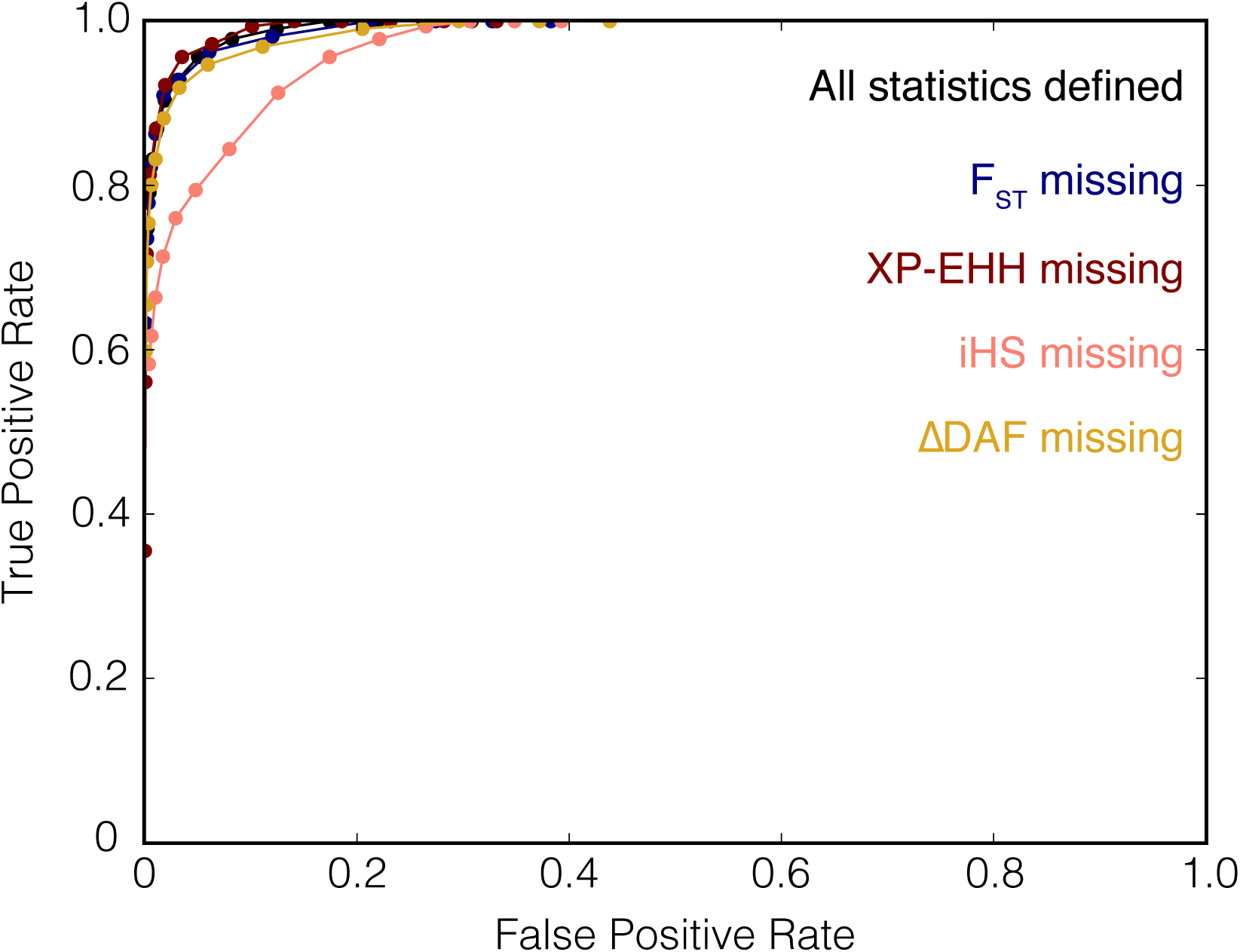
Effect of undefined statistics on classification power. In simulation, we gathered all sites with no missing statistics, and then calculated SWIF(r) sweep probabilities for the complete set of statistics, and with each statistic missing. The ROC curves show the effect of each of these missing statistics on the power of SWIF(r). XP-EHH, *F_ST_*, and ΔDAF all have very little effect, but a missing value for iHS does result in lower power to distinguish adaptive mutations from neutral mutations. This differential loss of power is not surprising, since iHS is extremely useful for identifying incomplete sweeps, while XP-EHH, ΔDAF, and *F_ST_* can more easily compensate for each other, as they are all most powerful for sweeps that are complete in the population of interest (Supplementary Figure 7).

**Supplementary Figure 34.**
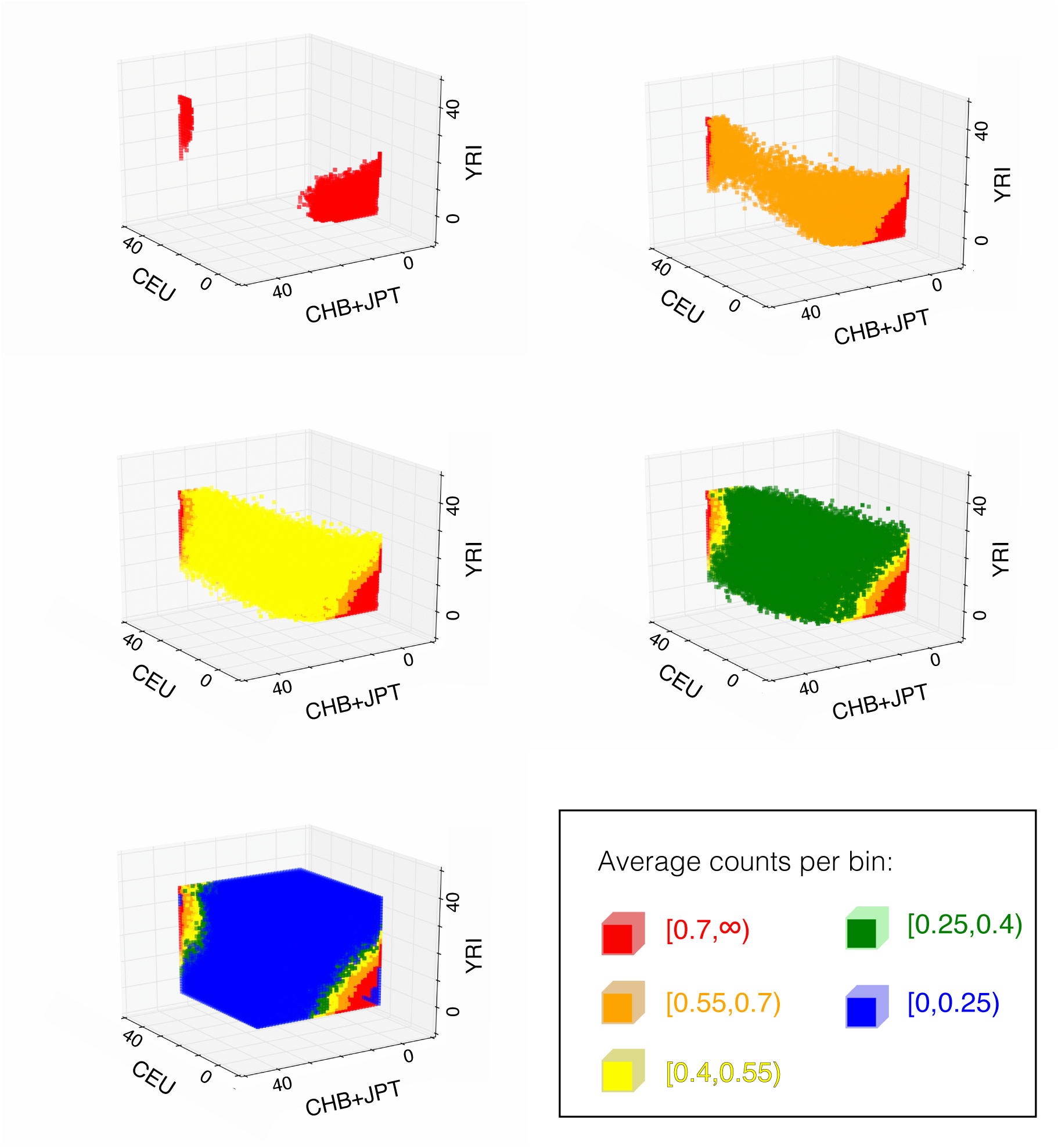
Illustration of step 2 of ascertainment modeling algorithm. For modeling ascertainment in the SNP array used to sequence the ‡Khomani San individuals in our selection scan, we defined five intervals that span the range of average count-per-bin of 40 × 40 × 40 SFS space, from less than 0.25 in blue, to greater than 0.7 in red. The set of bins defined by a given color are the “SFS regions” referred to in the Online Methods. The five plots are shown because lower-count regions tend to conceal the higher-count regions in the two-dimensional projection.

**Supplementary Table 1.** Undefined statistics in 1000 Genomes. Fraction of variant sites, out of 11 million, for which each of four component statistics are undefined within each of three populations.

| Population | $F_{ST}$ | XP-EHH | iHS | $\Delta DAF$ |
| --- | --- | --- | --- | --- |
| CEU | 0.5% | 0.2% | 77.5% | 0% |
| CHB+JPT | 0.5% | 0.3% | 78.9% | 0% |
| YRI | 0.5% | 0.3% | 67.4% | 0% |

**Supplementary Table 2**

**Selective sweep targets identified by SWIF(r) in the 1000 Genomes phase 1 dataset.** Spreadsheet contains all SNPs that have a posterior sweep probability greater than 10% for each of the three populations. SNPs are identified by rsid (column A), chromosome (B), and position in genome build hg19 (C). The uncalibrated posterior sweep probability calculated by SWIF(r) is shown in column D, the calibrated probability using isotonic regression is shown in E, and the calibrated value using smoothed isotonic regression is in F (Supplementary Figure 1, Supplementary Figure 3). SNPs are annotated by gene (G), mutation type (H), and genes within 100kb (I).

**Supplementary Table 3**

**Support for positive selection in genes identified by SWIF(r) in the 1000 Genomes phase 1 dataset.** For each of the three populations (**CEU, CHB+JPT, YRI**), we list every gene containing a SNP with posterior sweep probability over 10% (column A), the number of such SNPs (B), and the maximum sweep probability over all such SNPs (C). The citations in the remaining columns are studies in which the gene was implicated in a positive selection scan in the population of interest. Information about these citations can be found in the last sheet of this spreadsheet.

**Supplementary Table 4.** False Discovery and Positive Predictive Value rates for SWIF(r) predictions in the 1000 Genomes dataset. Estimating empirical false discovery rates in observed genomic data requires defining a set of neutrally evolving genomic sites, which is extremely difficult to do. We use two proxies for defining these sites: first, we use “non-conserved non-coding” sites, defined as those lying at least 2kb away from annotated genes, and having a phastcons conservation score of zero, following Hernandez *et al.*^27^. We consider this to be an extremely permissive definition, as sites with low conservation scores may well be tagging nearby variants with higher scores that may or may not be present in the genotype dataset, and intergenic regions contain many regulatory and functional elements. The second proxy we use is a set of 15 putatively neutral regions comprising ∼216kb identified by Gazave *et al.*^28^ using a range of criteria indicating neutrality. For the first proxy, we calculated the false positive rate among these sites at different sweep probability thresholds, then extended this false positive rate genome-wide to obtain a prediction for the total number of false positives that would be expected in a genome scan in each population of interest (YRI, CEU, CHB+JPT). Dividing the number of predicted false positives by the total number of sites that SWIF(r) identifies above a given threshold in each population gives us an estimate of the false discovery rate (FDR), and inversely of the positive predictive value (PPV) of SWIF(r). We find in general that we predict lower FDR at higher posterior probability cutoffs, indicating that high-probability SNPs are more likely to be true signals than lower-probability SNPs. The last column contains the fraction of total predictions made by SWIF(r) that we estimate to be true, based on these values. For YRI, we only show results up to a sweep probability threshold of 0.5 because we only observe one false positive at this threshold, and we thus lack the resolution to estimate FDR or PPV for higher thresholds. Again, we note that some SNPs labeled as false positives by this scheme may nonetheless tag nearby selective sweeps, and we consider these false discovery rates to be very conservative. Using the much stricter definition of neutrally evolving sequence developed by Gazave *et al.*^28^, we see no false positives in any of the three populations, even at a sweep probability threshold of 10%.

| Population | Sweep Probability Threshold | FDR | PPV | Estimated True/Total |
| --- | --- | --- | --- | --- |
| YRI | 0.5 | 0.514 | 0.486 | 7/15 |
| CEU | 0.5 | 0.421 | 0.579 | 138/238 |
|  | 0.6 | 0.469 | 0.531 | 96/181 |
|  | 0.7 | 0.459 | 0.541 | 91/168 |
|  | 0.8 | 0.504 | 0.496 | 76/153 |
|  | 0.9 | 0.367 | 0.633 | 80/126 |
|  | 0.95 | 0.350 | 0.650 | 71/110 |
|  | 0.98 | 0.339 | 0.661 | 60/91 |
| CHB+JPT | 0.5 | 1.00 | 0.00 | 0/342 |
|  | 0.6 | 0.801 | 0.199 | 46/231 |
|  | 0.7 | 0.806 | 0.194 | 39/201 |
|  | 0.8 | 0.878 | 0.122 | 19/158 |
|  | 0.9 | 0.461 | 0.539 | 63/117 |
|  | 0.95 | 0.308 | 0.692 | 69/100 |
|  | 0.98 | 0.214 | 0.786 | 72/58 |

**Supplementary Table 5.** Selection coefficients for simulations. Selection coefficients for each sweep scenario are fully determined by the start time of the sweep (all sweeps end at the time of sampling), the number of years per generation, the final allele frequency of the beneficial allele, and the effective population size for the population within which the sweep occurs. As in Schaffner *et al.*^9^, *N_e_* is 24000 for West Africa (YRI), 7700 each for Europe (CEU) and East Asia (CHB and JPT), and as in Gronau et *al.^8^, N_e_* is 21000 for the ‡Khomani San. The demographic model we use for the 1000 Genomes populations (YRI, CEU, CHB, JPT) assumes 20 years/generation^9^, while the one we use for the ‡Khomani San study assumes 25 years/generation^8^.

| Start Time | Final Allele Frequency | Selective Strength (Europe, East Asia) | Selective Strength (West Africa) | Selective Strength (‡Khomani San) |
| --- | --- | --- | --- | --- |
| 5kya-present | 0.2 | 0.033 | 0.0376 | 0.0463 |
|  | 0.4 | 0.0369 | 0.0415 | 0.0512 |
|  | 0.6 | 0.0402 | 0.0447 | 0.0553 |
|  | 0.8 | 0.0441 | 0.0487 | 0.0602 |
|  | 1 | 0.0771 | 0.0862 | 0.1065 |
| 10kya-present | 0.2 | 0.0165 | 0.0188 | 0.0231 |
|  | 0.4 | 0.0185 | 0.0207 | 0.0256 |
|  | 0.6 | 0.0201 | 0.0224 | 0.0276 |
|  | 0.8 | 0.0221 | 0.0243 | 0.0301 |
|  | 1 | 0.0386 | 0.0431 | 0.0532 |
| 15kya-present | 0.2 | 0.011 | 0.0125 | 0.0154 |
|  | 0.4 | 0.0123 | 0.0138 | 0.0171 |
|  | 0.6 | 0.0134 | 0.0149 | 0.0184 |
|  | 0.8 | 0.0147 | 0.0162 | 0.0201 |
|  | 1 | 0.0257 | 0.0287 | 0.0355 |
| 20kya-present | 0.2 | 0.0083 | 0.0094 | 0.0116 |
|  | 0.4 | 0.0092 | 0.0104 | 0.0128 |
|  | 0.6 | 0.01 | 0.0112 | 0.0138 |
|  | 0.8 | 0.011 | 0.0122 | 0.0150 |
|  | 1 | 0.0193 | 0.0216 | 0.0266 |
| 25kya-present | 0.2 | 0.0066 | 0.0075 | 0.0093 |
|  | 0.4 | 0.0074 | 0.0083 | 0.0102 |
|  | 0.6 | 0.008 | 0.0089 | 0.0111 |
|  | 0.8 | 0.0088 | 0.0097 | 0.0120 |
|  | 1 | 0.0154 | 0.0172 | 0.0213 |
| 30kya-present | 0.2 | 0.0057 | 0.0065 | 0.0080 |
|  | 0.4 | 0.0064 | 0.0072 | 0.0088 |
|  | 0.6 | 0.0069 | 0.0077 | 0.0095 |
|  | 0.8 | 0.0076 | 0.0084 | 0.0103 |
|  | 1 | 0.0133 | 0.0149 | 0.0184 |
| 41kya-36kya | 0.2 |  |  | 0.0462 |
|  | 0.4 |  |  | 0.0512 |
|  | 0.6 |  |  | 0.0553 |
|  | 0.8 |  |  | 0.0601 |
|  | 1.0 |  |  | 0.1064 |
| 46kya-36kya | 0.2 |  |  | 0.0231 |
|  | 0.4 |  |  | 0.0256 |
|  | 0.6 |  |  | 0.0276 |
|  | 0.8 |  |  | 0.0301 |
|  | 1.0 |  |  | 0.0532 |
| 52kya-47kya | 0.2 |  |  | 0.0463 |
|  | 0.4 |  |  | 0.0512 |
|  | 0.6 |  |  | 0.0553 |
|  | 0.8 |  |  | 0.0602 |
|  | 1.0 |  |  | 0.1065 |
| 57kya-47kya | 0.2 |  |  | 0.0231 |
|  | 0.4 |  |  | 0.0256 |
|  | 0.6 |  |  | 0.0276 |
|  | 0.8 |  |  | 0.0301 |
|  | 1.0 |  |  | 0.0532 |

**Supplementary Table 6.** False Discovery and Positive Predictive Value rates for SWIF(r) predictions in the ‡Khomani dataset. Estimating empirical false discovery rates in observed genomic data requires defining a set of neutrally evolving genomic sites, which is extremely difficult to do. We use two proxies for defining these sites: first, we use “non-conserved non-coding” sites, defined as those lying at least 2kb away from annotated genes, and having a phastcons conservation score of zero, following Hernandez *et al.*^27^. We consider this to be an extremely permissive definition, as sites with low conservation scores may well be tagging nearby variants with higher scores that may or may not be present in the genotype dataset, and intergenic regions contain many regulatory and functional elements. The second proxy we use is a set of 15 putatively neutral regions comprising ∼216kb identified by Gazave *et al.*^28^ using a range of criteria indicating neutrality. For the first proxy, we calculated the false positive rate among these sites at different sweep probability thresholds, then extended this false positive rate genome-wide to obtain a prediction for the total number of false positives that would be expected in a genome scan. Dividing the number of predicted false positives by the total number of sites that SWIF(r) identifies above a given threshold in the ‡Khomani San gives us an estimate of the false discovery rate (FDR), and inversely of the positive predictive value (PPV) of SWIF(r). The last column contains the fraction of total predictions made by SWIF(r) that we estimate to be true, based on these values. For YRI, we only show results up to a sweep probability threshold of 0.5 because we only observe one false positive at this threshold, and we thus lack the resolution to estimate FDR or PPV for higher thresholds. As in Supplementary Table 4, we note that some SNPs labeled as false positives by this scheme may nonetheless tag nearby selective sweeps, and we consider these false discovery rates to be very conservative. Using the much stricter definition of neutrally evolving sequence developed by Gazave *et al.*^28^, we see no false positives in any of the three populations, even at a sweep probability threshold of 10%.

| Sweep Probability Threshold | FDR | PPV | Estimated True/Total |
| --- | --- | --- | --- |
| 0.5 | 0.563 | 0.437 | 47/108 |
| 0.6 | 0.615 | 0.385 | 35/90 |
| 0.7 | 0.560 | 0.440 | 35/79 |
| 0.8 | 0.439 | 0.561 | 35/63 |
| 0.9 | 0.474 | 0.526 | 18/35 |
| 0.95 | 0.572 | 0.428 | 12/29 |
| 0.98 | 0.251 | 0.749 | 16/22 |

**Supplementary Table 7**

**Adaptive loci identified by SWIF(r) in the ‡Khomani San array dataset.** Spreadsheet contains all SNPs that have posterior sweep probability greater than 10% in the ‡Khomani array dataset^23^. SNPs are identified by rsid (column A), chromosome (B), and position in hg19 (D). The derived allele frequency of the SNP in the {Khomani is in column C. The uncalibrated posterior sweep probability calculated by SWIF(r) is in column E, the calibrated probability using isotonic regression is shown in F, and the calibrated value using smoothed isotonic regression is in G (Supplementary Figure 2, Supplementary Figure 3). The un-calibrated posterior sweep probability is broken down into posterior probabilities for “recent” (<30kya) and “ancient” (36-47kya) in columns H and I, respectively. SNPs are also annotated by gene (J), mutation type (K), and genes within 100kb (L).

**Supplementary Table 8**

**Exome data from 45 ‡Khomani San individuals reveals variants that have functional consequence and large allele frequency differences relative to other worldwide populations within SWIF(r)-identified genes involved in metabolism and obesity.** Spreadsheet contains SNPs of interest in exome data (Martin *et al.*^29^) within genes highlighted in Figure 3B. Each SNP is annotated with rsid and chromosome position in genome build hg19. The variant type was determined using the UCSC Genome Browser^30^, and frequencies in worldwide populations were taken from phase 3 of the 1000 Genomes project^16^ where available, otherwise, frequencies were taken from the Human Genome Diversity Project and/or the ExAC browser^31^.

## Supplementary Note

### cosi demographic parameter file for 1000 Genomes dataset

`length 1000000`

`mutation_rate 1.5e-8`

`recomb_file <filename>`

`gene_conversion_rate 4.5e-9`

`pop_define 1 european`

`pop_define 2 asian`

`pop_define 3 african`

`#initial sizes and sample sizes`

`pop_size 1 7700`

`sample_size 1 120`

`pop_size 2 7700`

`sample_size 2 120`

`pop_size 3 24000`

`sample_size 3 120`

`#Migration start`

`pop_event migration_rate “afr to eur migration” 3 1 1505 .000032`

`pop_event migration_rate “eur to afr migration” 1 3 1504 .000032`

`pop_event migration_rate “afr to as migration” 3 2 1503 .000008`

`pop_event migration_rate “as to afr migration” 2 3 1502 .000008`

`#Migration end`

`pop_event migration_rate “afr to eur migration” 3 1 1996 0`

`pop_event migration_rate “eur to afr migration” 1 3 1995 0`

`pop_event migration_rate “afr to as migration” 3 2 1994 0`

`pop_event migration_rate “as to afr migration” 2 3 1993 0`

`#Recent Bottlenecks:`

`pop_event bottleneck “african bottleneck” 3 1997 .008`

`pop_event bottleneck “asian bottleneck” 2 1998 .067`

`pop_event bottleneck “european bottleneck” 1 1999 .02`

`#Population splits:`

`pop_event split “asian and european split” 1 2 2000`

`pop_event split “out of Africa” 3 1 3500`

`#Out-of-africa bottleneck`

`pop_event bottleneck “OoA bottleneck” 1 3499 .085`

`#Ancestral expansion`

### cosi demographic parameter file for ‡Khomani dataset

`length 1000000`

`mutation_rate 1.5e-8`

`recomb_file <filename>`

`gene_conversion_rate 4.5e-9`

`pop_define 1 european #E`

`pop_define 2 han_chinese #H`

`pop_define 3 yoruban #Y`

`pop_define 4 khomani #K`

`#initial sizes and sample sizes`

`pop_size 1 9700`

`sample_size 1 164`

`pop_size 2 5800`

`sample_size 2 372`

`pop_size 3 17800`

`sample_size 3 174`

`pop_size 4 21000`

`sample_size 4 90`

`#population size changes`

`pop_event change_size “H” 3 1240 3500`

`pop_event change_size “HE” 2 1441 1200`

`pop_event change_size “HEY” 3 1881 11500`

`pop_event change_size “HEYK” 4 5241 8700`

`#migration to Khomani`

`pop_event migration_rate “YRI to Khomani migration” 3 4 14 0.227`

`pop_event migration_rate “YRI to Khomani migration end” 3 4 15 0`

`pop_event migration_rate “CEU to Khomani migration” 1 4 6 0.179`

`pop_event migration_rate “CEU to Khomani migration end” 1 4 7 0`

`#population splits:`

`pop_event split “CE” 2 1 1440`

`pop_event split “CEY” 3 2 1880`

`pop_event split “CEYK” 4 3 5240`

### Selection Statistic Calculations

For each segregating site within the neutral simulations, and for the adaptive site in sweep simulations, we computed component selection statistics (*F_st_*, XP-EHH, iHS, ΔiHH, ΔDAF) with respect to a population of interest (the population undergoing a sweep in sweep simulations, and a population chosen uniformly at random in the case of neutral simulations), ignoring sites for which the derived allele frequency in the population of interest was zero. *F_ST_* for each of the pairwise population comparisons involving the population of interest was computed as in Weir *et al.*^32^, and then averaged together. iHS was computed with selscan^33^. ΔiHH was calculated as defined in Grossman *et al.*^15^: ΔiHH|iHH_ancestral_-iHH_derived_|, where iHH is the integrated haplotype homozygosity defined in Voight et al.^4^. ΔDAF was also calculated as defined in Grossman *et al.*^15^: 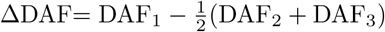 where DAF_1_ is the derived allele frequency in the population of interest, and DAF_2_ and DAF_3_ are the derived allele frequencies in the other two populations. XP-EHH was computed with a minor alteration of `selscan` in which *EHH* is computed as in Wagh *et al.*^34^ (see “Adaptation of `selscan` for better performance on incomplete sweeps”), and XP-EHH values were also normalized with population-specific mean and standard deviation, learned from neutral simulations, to correct for inherent biases based on linkage disequilibrium structure.

Small adjustments were required for computing these component statistics using the ‡Khomani San simulations and genotype array data^23^. In these analyses, there are three outgroups (western Africa, Europe, and eastern Asia) instead of two: XP-EHH was defined as the maximum XP-EHH value across the three comparisons; and ΔDAF was defined as

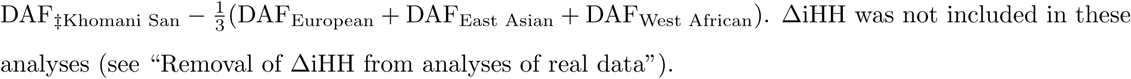

Results for each component statistic were normalized within 1MB regions, with iHS and ΔiHH being normalized first within frequency bins as in Voight *et al.*^4^. For the CMS, we learned one-dimensional probability distributions for each (scenario, component statistic) pair in 60 evenly spaced bins, with minimum and maximum values below, chosen to encompass the full range of values observed across all neutral and sweep simulations:

| component statistic | minimum | maximum |
| --- | --- | --- |
| mean $F_{ST}$ | -1 | 9 |
| maximum XP-EHH | -3 | 10 |
| iHS | -5 | 4 |
| $\Delta iHH$ | -3 | 10 |
| $\Delta DAF$ | -5 | 8 |

When computing statistics for either 1000 Genomes or ‡Khomani San genotype data, component statistics were normalized genome-wide, following Grossman *et al*.^14^.

### Adaptation of selscan^33^ for better performance on incomplete sweeps

XP-EHH is defined as log 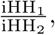 where iHH_1_ and iHH_2_ are “integrated haplotype homozygosities” for the population of interest and a reference population respectively. iHH is computed as the integral under the EHH curve, where EHH at a distance *x* from the core SNP is canonically computed as follows:

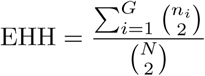

where *N* is the total number of chromosomes, *G* is the number of distinct haplotypes, and *n_i_* is the number of chromosomes of distinct type *i*. In the case of incomplete sweeps from a *de novo* mutation, this definition somewhat counterintuitively leads to negative values of XP-EHH. This is due to the fact that the numerator has an upper bound of 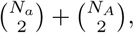 where *N_a_* and *N_A_* are the number of chromosomes that have each of the two possible alleles at the core SNP. In the reference population, there is a larger upper bound of 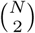, leading to a larger value of iHH for the reference population than for the population with the adaptive mutation.

Following Wagh *et al.*^34^, we instead define EHH as follows:

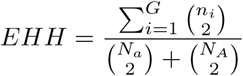

When XP-EHH is defined this way, it is far more powerful for detecting sweeps, and does not return negative values for incomplete sweeps. We modified `selscan`’s source code to implement this change. These modifications can be found at https://github.com/lasugden/selscan (original software at https://github.com/szpiech/selscan).

### Removal of ΔiHH from analyses of real data

While other statistics are distributed approximately normally after normalization, ΔiHH maintains a very long tail (1-2 orders of magnitude longer than the tails of other statistics), which is exacerbated by normalization. This problem is compounded by the fact that ΔiHH is an absolute value, leading to high scores even when the ancestral haplotype is longer than the derived haplotype. In Supplementary Figure 5, we show the distribution of ΔiHH and iHS values for sites with SWIF(r) sweep probability ≥ 50% in the 1000 Genomes dataset using SWIF(r) with ΔiHH included as a component statistic. Note the scale of both statistics; while iHS ranges from −4 to 4, ΔiHH ranges from 0 to 100 (in simulations, the probability mass for ΔiHH lies mostly below 5, and entirely below 14). The plot shows that ΔiHH can be extremely high, even for values of iHS that are positive and thus provide evidence against selective sweeps. Furthermore, the more negative iHS values correspond to more moderate ΔiHH values, and not the most extreme ones, thus leading to a large number of false positives. For these reasons, we removed ΔiHH as a component statistic for analysis of simulations and genotype data from the 1000 Genomes and the ‡Khomani San.

### Processing of 1000 Genomes Data

We performed a genome-wide scan for selective sweeps in four populations (YRI, CEU, CHB, JPT) using phase 1 of the 1000 Genomes Project (May 2011 release), with CHB and JPT grouped together and representing East Asia. We filtered the samples to omit children from parent-child pairs and trios (NA07048, NA10847, and NA10851 from CEU and NA19129 from YRI), and we only analyzed single-nucleotide variants. We also removed loci that were monomorphic within the filtered set of unrelated individuals across the four populations analyzed. We used ancestral allele information provided by the 1000 Genomes Project.

### Calculation of migration rates from YRI and CEU to ‡Khomani San

Uren *et al.*^23^ use the software package Tracts^35^ to infer the magnitude of genetic contributions to present-day ‡Khomani San individuals from three source populations: KhoeSan (data presented in Uren *et al.*^23^ and Schuster *et al.*^36^), LWK (1000 Genomes), and CEU (1000 Genomes). In our simulations, we use YRI as a proxy population for LWK (i.e. migration rates learned in Uren et al. from LWK are implemented in our demographic model as migration rates from YRI), as both are Bantu-speaking populations, which all originate from west central Africa. Therefore both populations provide an appropriate source for the Bantu ancestry in the ‡Khomani San. The migration rates reported by the Tracts software represent the proportion of the target population that is replaced at a given generation by a source population, starting in this case 14 generations ago. Prior to 14 generations ago, we assume that the ‡Khomani San population is made up of 100% KhoeSan ancestry, and we converted the Tracts migration rates into migration rates from only two source populations (YRI and CEU) into the third (‡Khomani San). We achieved this by going through an intermediate step in which we calculated a matrix of ancestry proportions at each generation. The table below shows the tracts output from Uren *et al.*^23^ (migration rates *m_S_*, *m_Y_*, and *m_C_* at each generation from 14 generations ago to present. We then calculate the ancestry proportions at each generation, initializing at generation 14 (*m_Y_* and *m_S_* at generation 14 add to 1, indicating that the ancestry proportions at this time point are equal to the migration rates), and working backwards. If we define 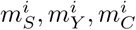 to be the migration rates at generation *i* in this 3-source population model, and 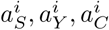 to be the ancestry proportions at generation *i*, then we can calculate these by the following recursive formulas, where 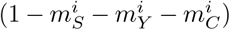 represents the fraction of the population not being replaced at the given generation.

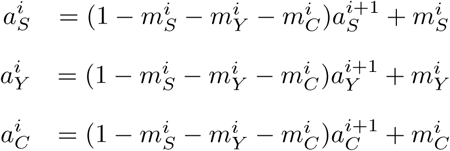

The entries in the following table for *a_S_*, *a_Y_*, and *a_C_* are calculated using these formulas. Finally, we need to convert these ancestry proportions into migration rates for a model with two population sources (YRI and CEU). For simplicity, we assume a one-generation migration pulse from YRI at generation 14, and a one-generation migration pulse from CEU at generation 7. We note that in order to achieve a present-day CEU ancestry proportion of 0.179, the CEU migration rate must be 0.179 (the migration rate is again defined as the fraction of the population being replaced by the source population). Finally, to achieve a present-day YRI ancestry proportion of 0.186, the migration rate from YRI must be 0.227, since (1 – 0.179) × 0.227 = 0.186 (this accounts for the proportion of YRI ancestry that gets replaced by CEU ancestry in generation 7).

| generation | $m_S$ | $m_Y$ | $m_C$ | $a_S$ | $a_Y$ | $a_C$ |
| --- | --- | --- | --- | --- | --- | --- |
| 1 | 0 | 0 | 0 | 0.635 | 0.186 | 0.179 |
| 2 | 0 | 0 | 0 | 0.635 | 0.186 | 0.179 |
| 3 | 0 | 0 | 0 | 0.635 | 0.186 | 0.179 |
| 4 | 0.240 | 0.031 | 0 | 0.635 | 0.186 | 0.179 |
| 5 | 0.052 | 0.005 | 0 | 0.542 | 0.212 | 0.246 |
| 6 | 0 | 0 | 0.151 | 0.520 | 0.219 | 0.261 |
| 7 | 0 | 0 | 0.130 | 0.612 | 0.258 | 0.130 |
| 8 | 0 | 0 | 0 | 0.703 | 0.297 | 0 |
| 9 | 0 | 0 | 0 | 0.703 | 0.297 | 0 |
| 10 | 0 | 0 | 0 | 0.703 | 0.297 | 0 |
| 11 | 0 | 0 | 0 | 0.703 | 0.297 | 0 |
| 12 | 0 | 0 | 0 | 0.703 | 0.297 | 0 |
| 13 | 0.527 | 0.222 | 0 | 0.703 | 0.297 | 0 |
| 14 | 0.703 | 0.297 | 0 | 0.703 | 0.297 | 0 |

### Generation of exome and array data in San population

‡Khomani San individuals were sampled in 2006 in Upington, South Africa and neighboring villages. Institution Review Board (IRB) approval for assessment of genetic diversity and ancestry inference was obtained from Stanford University [Protocol 13829] and Stony Brook University [Protocol 727494-5]. Still-living individuals were re-consented in 2011 (IRB approved from Stanford University and Stellenbosch University, South Africa). {Khomani N|u-speaking individuals, local community leaders, traditional leaders, non-profit organizations and a legal counselor were all consulted about the project aims before DNA collection commenced. All individuals initially orally consented to participate in the project in the presence of a witness fluent in the native language, and were re-consented with written consent. DNA was collected via saliva (Oragene kits). Ancestry and genotyping details of individuals included here can be found in Uren *et al.*^23^.

90 KhoeSan DNA samples were captured with 3 exome platforms: 74 samples on an Agilent SureSelect Human All Exon V2 44Mb, 8 samples on an Agilent SureSelect Human All Exon 50Mb, and 8 samples on an Agilent SureSelect Human All Exon V4+UTRs 71Mb (Martin *et al.*^29^). Illumina short-read sequencing data were jointly processed according to the best practice pipeline of the 1000 Genomes Project^16^. Reads were aligned to the hg19 reference genome using bwa-mem 0.7.10^37^. The resultant BAM files were then sorted and marked for duplicate reads using the Picard v1.92 toolkit (http://broadinstitute.github.io/picard/). The following programs were then run with GATKv3.2.2^38^: RealignerTargetCreator, IndelRealigner, BaseRecalibrator, PrintReads, HaplotypeCaller, GenotypeGVCFs, and VariantRecalibrator, and ApplyRecalibration. During the HaplotypeCaller step, we filtered reads to include only the Agilent capture regions ± 100 bp of padding.

For phasing, exome data were merged with Illumina SNP arrays for each of 87 individuals to improve accuracy by providing a broader SNP scaffold. After merging and filtering to 5% genotyping missingness using vcftools, 759,586 SNPs remained. Data were phased using a two-step phasing process as follows: first, 25 related individuals, consisting of 3 trios and 8 duos, were used to create a family reference panel; after phasing unrelated individuals using default protocols in SHAPEIT2^39^, pedigree information was used via duo-trio phasing with duoHMM^40^ to inform the phasing of the unrelated individuals in a second step, improving phasing accuracy overall by correcting phase switch errors.

{Khomani San individuals were genotyped on two SNP array platforms: the Illumina OmniExpress and OmniExpressPlus chips. Only SNPs shared between these two platforms^23^ were retained for investigation of adaptive sites, in order to avoid allele frequency biases related to platform choice; there is broad overlap between these arrays, with the OmniExpressPlus SNP array containing an additional 250k sites. 86 individuals were genotyped and were phased using only pedigree information. This pedigree-phased dataset provides the highest possible SNP density for these individuals and, after quality control filtering, included just under 650k SNPs.

For the analyses presented here, we selected 45 unrelated ‡Khomani San individuals with low rates of recent European and Bantu-derived admixture from the phased datasets. These individuals were identified by running ADMIXTURE v1.2^41^ on a joint dataset of Illumina SNP arrays consisting of diverse populations of individuals^23^ and selecting those KhoeSan individuals with >90% Khoesan ancestry at *K* = 6. SIx was determined to be the K value that best fit the data, using both cross-validation procedures and cluster appearance.

### SWIF(r) signals associated with muscle-based phenotypes

Both *MYH15* and *TTN* have been associated with obesity and metabolism phenotypes (Table 1). Additionally, both genes encode striated muscle. *MYH15* has been associated with coronary heart disease^42^, and *TTN* mutations have been associated with cardiomyopathy^43^, with RNAseq data indicating that expression is highest in heart tissue^44^. Associations for *MYH15* and TTN with the obesity and metabolism phenotype may be a consequence of these other functions and associations. Exome support for selection acting within these genes is given below; allele frequencies for populations other than the ‡Khomani San refer to frequencies found in the 1000 Genomes phase 3 dataset^16^.

*TTN* (titin): We find 6 missense mutations within 50kb of the SWIF(r) signal; in itself, this may not be unusual given the exon richness in this gene. However, many of the mutations have a population frequency of approximately 50% in the Khomani San, while being absent or having much lower frequencies in other worldwide populations. For example, the derived asparagine to isoleucine (rs11900987) mutation is conserved amongst mammals^30^, <1% in other human populations, but present at 48% in our San sample. Other mutations segregating at similar frequency within 50kb suggest that a high frequency haplotype is under adaptive evolution the ‡Khomani San.

*MYH15* (myosin heavy chain 15): Our exome analysis found two missense mutations in *MYH15* with large allele frequency deviations. the G allele of rs9868484, which lies ∼4kb from the SWIF(r) signal, is at a frequency of 71% in our sample, relative to a maximum frequency of 40% elsewhere in Africa. The T allele of rsi078456, ∼50kb from the SWIF(r) signal, is at a frequency of 22%, relative to a maximum of 4% worldwide. In addition, a splice region variant ∼38kb from the SWIF(r) signal, rs113330737, has derived allele frequency 46% in the {Khomani sample relative to a maximum of 1% worldwide.

